# Generation of human nociceptor-enriched sensory neurons for the study of pain-related dysfunctions

**DOI:** 10.1101/2022.02.19.480828

**Authors:** Anna-Katharina Holzer, Christiaan Karreman, Ilinca Suciu, Lara-Seline Furmanowsky, Harald Wohlfarth, Dominik Loser, Wilhelm G Dirks, Emilio Pardo González, Marcel Leist

## Abstract

*In vitro* models of the peripheral nervous system would benefit from further refinements to better support studies on neuropathies. In particular, the assessment of pain-related signals is still difficult in human cell cultures. Here, we harnessed induced pluripotent stem cells (iPSCs) to generate peripheral sensory neurons enriched in nociceptors. The objective was to generate a culture system with signaling endpoints suitable for pharmacological and toxicological studies. Neurons generated by conventional differentiation protocols expressed moderate levels of P2X3 purinergic receptors and only low levels of TRPV1 capsaicin receptors, when maturation time was kept to the upper practically-useful limit of 6 weeks. As alternative approach, we generated cells with an inducible *NGN1* transgene. Ectopic expression of this transcription factor during a defined time window of differentiation resulted in highly-enriched nociceptor cultures, as determined by functional (P2X3 and TRPV1 receptors) and immunocytochemical phenotyping, complemented by extensive transcriptome profiling. Single cell recordings of Ca^2+^-indicator fluorescence from >9,000 cells were used to establish the “fraction of reactive cells” in a stimulated population as experimental endpoint, that appeared robust, transparent and quantifiable. To provide an example of application to biomedical studies, functional consequences of prolonged exposure to the chemotherapeutic drug oxaliplatin were examined at non-cytotoxic concentrations. We found (i) neuronal (allodynia-like) hypersensitivity to otherwise non-activating mechanical stimulation that could be blocked by modulators of voltage-gated sodium channels; (ii) hyper-responsiveness to TRPV1 receptor stimulation. These findings and several other measured functional alterations indicate that the model is suitable for pharmacological and toxicological studies related to peripheral neuropathies.

## Introduction

*In vitro* models of the human peripheral nervous system (PNS) are still relatively scarce. They are required to study chemotherapy-induced peripheral neuropathy (CIPN) and other impairments of the PNS. Of particular interest are systems that allow the assessment of agents that functionally impair sensory neurons.

Cell-based model systems for the PNS are still mostly based on non-human cells, like rat dorsal root ganglion (DRG) neurons. Such DRG cultures have drawbacks concerning e.g., their comparability, and human-specific functions may only be modelled partially [1]. In the past decade, stem cell technology has provided novel alternatives. The fundamental principles of generating peripheral neurons from human induced pluripotent stem cells (iPSCs) were described in 2012 by the Studer laboratory [2]. This protocol uses neuralization of iPSCs by dual SMAD inhibition. The fine-tuning of differentiation towards the sensory neuron fate is subsequently achieved by small molecule inhibitors combined with neurotrophins.

*In vitro* model systems for the PNS are indispensable for toxicity testing, as peripheral neurotoxicants are often not identified by models of the central nervous system (CNS) [3, 4]. The sensory neuronal subclass of nociceptors is of specific interest in CIPN research. Neuropathies involving this particular subpopulation [5–7] are amongst the side effects that most profoundly decrease the quality of life of chemotherapy-receiving patients [8, 9].

Sensory neurons can be classified into nociceptors, mechanoceptors and proprioceptors. The first group expresses the nerve growth factor (NGF)-receptor TRKA (encoded by *NTRK1*) [10] during maturation, while the others depend on TRKB and TRKC tyrosine kinase signaling. While all peripheral neurons are derived from neural crest progenitors, the TRKA-expressing neurons develop from the subgroup of NGN1-positive neural crest cells [11, 12]. They can be further divided into nociceptor-subgroups: Peptidergic neurons release the neuropeptides substance P and calcitonin gene-related peptide and maintain TRKA expression. Non-peptidergic neurons lose expression of TRKA upon maturation, and express the RET neurotrophin receptor instead [10]. Nociceptors can also be distinguished according to their expression of different cation channels like the transient receptor potential (TRP) channels or the purinergic receptor ion channels. Notable members of the TRP family are TRPV1, TRPM8 and TRPA1 channels. The respective major functions are the sensing of heat, cold or electrophilic chemicals [7]. Temperature-sensing TRP channels are polymodal and can respond to chemical agonists. A prominent example is the TRPV1 channel, which is activated by increased temperatures exceeding the threshold of ∼43°C, but also by vanilloid compounds like capsaicin [13]. The purinoceptor P2X3 is the main ATP-activated pain-related channel on nociceptors. As TRPV1 and P2X3 are only found on nociceptors and not on other sensory neurons (e.g. stretch receptors), they can serve as characteristic functional biomarkers [14–16].

De-regulations of ligand-activated and voltage-gated ion channels on peripheral neurons are known to contribute to the dose-limiting side-effects induced by the chemotherapeutic drug oxaliplatin [17, 18]. General neuronal hyperexcitability [18–21], thermal hyperalgesia and mechanical allodynia [18, 22–24] are characteristic features of acute oxaliplatin-induced peripheral neuropathy (OXAIPN). While all platinum drugs lead to structural damage upon prolonged treatment, the acute form of OXAIPN occurs largely independent of neurodegeneration [25].

The example of OXAIPN demonstrates the need for *in vitro* model systems that can identify functional impairments of the PNS. While assays to detect chemicals acting on neurite growth, neuroprogenitor migration or central neuronal signaling are well established [3, 4, 26–31], PNS systems optimized to detect functional impairments are still scarce, and data on the modulation of pain receptors are mostly not included [32].

Therefore, the aim of this study was to establish an *in vitro* system able to detect signaling alterations relevant for CIPN. A protocol to generate peripheral neurons with nociceptor features (PNN) from iPSCs was established. After an extensive phenotypic profiling, Ca^2+^-imaging was chosen as a quantitative endpoint for the assessment of pain-receptor signaling. A case study of oxaliplatin treatment was performed to demonstrate the relevance of our novel PNS model, which offers pain-receptor related functional endpoints for CIPN research. Thus, our study explored whether complex functional impairments of nociceptors are reliably detectable and quantifiable *in vitro*.

## Materials and Methods

### Materials

Unless mentioned otherwise, all chemicals and cell culture reagents were from Merck (Darmstadt, Germany). All antibodies and PCR primers used are compiled in dedicated tables in the supplementary materials file. There, also an extensive chapter on supplementary methods is included.

### Differentiation of sensory neurons from iPSC

We used the iPSC line Sigma iPSC0028 and derived from this the transgenic iPSC line Sigma-NGN1. Maintenance of the iPSCs was performed under xeno-free conditions [33] as detailed in the supplementary methods. The differentiation was performed according to Hoelting et al. (2015) [4, 30] with small modifications as shown in figure 1 and described in Klima et al. (2021) [4, 30].

**Figure 1:**
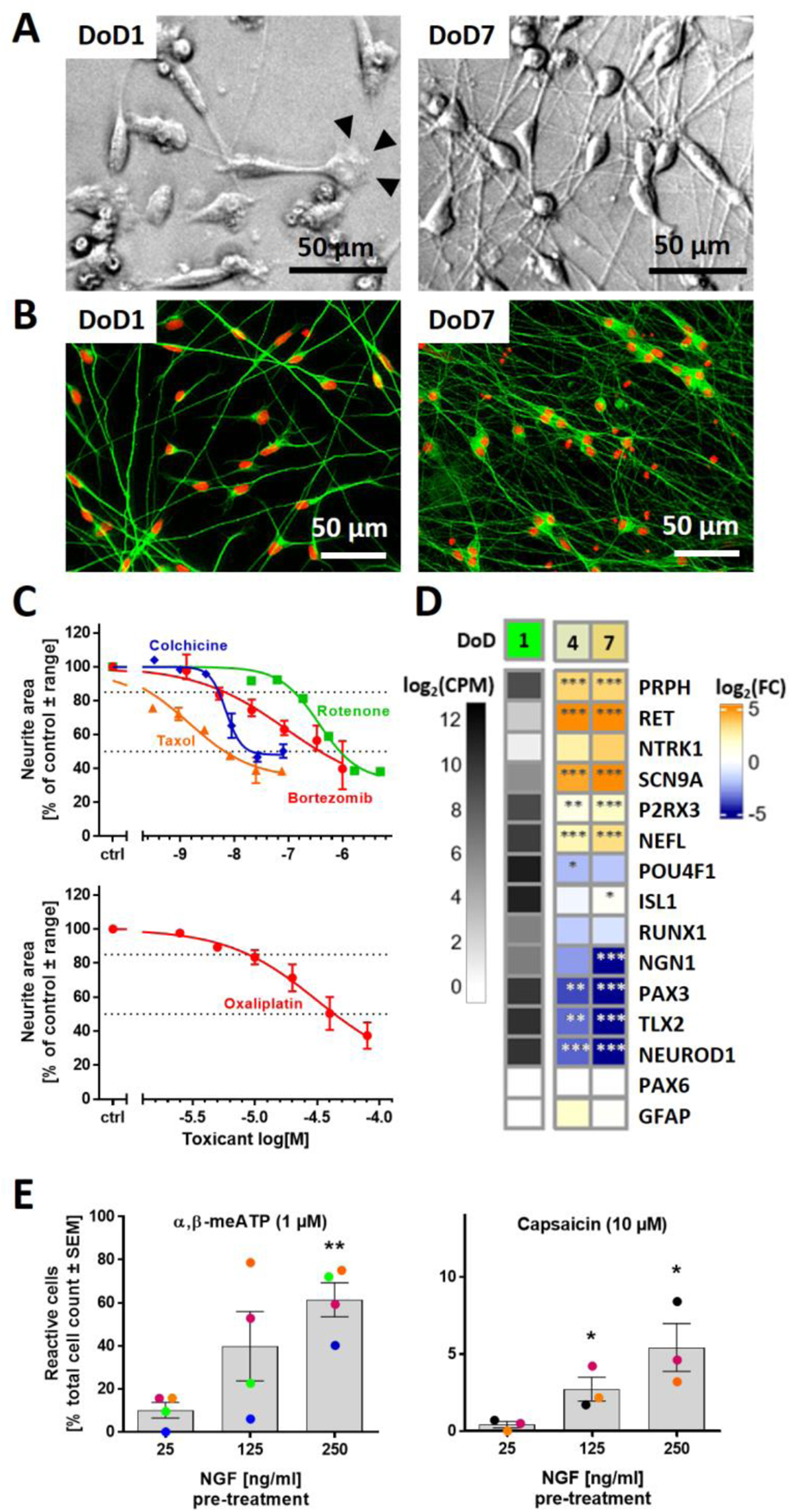
Human sensory neurons derived from iPSCs. Cells were pre-differentiated for 9 days and frozen (Fig. S1A). Counting of peripheral neuronal age in days of differentiation (DoD) started after thawing and plating (= DoD0). **(A)** Phase-contrast images of DoD1/DoD7 neurons. Arrowheads indicate a growth cone. **(B)** DoD1/DoD7 cultures stained for the neuronal cytoskeletal marker β-III-tubulin (green); DNA is shown in red. **(C)** Neurons were used in the PeriTox test to assess the effects of toxicant exposure (24h) on neurites. Data are given as mean ± range of 2-3 biological replicates. Viability was not significantly affected at the tested drug test concentrations (see Fig. S1B). **(D)** Gene expression levels were determined by the TempO-Seq method. The left column shows the absolute expression levels of selected marker genes on DoD1 in counts of the corresponding gene per 1 million reads (CPM). The data for DoD4/DoD7 show the fold change (FC) of the expression levels versus DoD1. The color scale uses log2FC units (see supplementary files for complete data sets). **(E)** Mature neurons (DoD25-45) were used for Ca^2+^-imaging. Cells reacting to the application of α,β-methylene ATP (α,β-meATP) and capsaicin were quantified. The effect of pre-treatment (48 h) with increased concentrations of nerve growth factor (NGF) on the percentage of reactive cells was assessed. Data displayed as bars are means ± SEM of 3-4 biological replicates. Color matching data points are derived from the same experiment. * p < 0.05, ** p < 0.005, when tested vs. control conditions (25 ng/ml NGF).

The detailed differentiation procedure is described in the supplementary methods (also see annex). In brief, iPSCs underwent neuralization induced by dual SMAD inhibition. Differentiation towards the sensory neuron fate was achieved by small molecule inhibition following the established literature [2]. After 9 days of differentiation (on DoD9’), immature peripheral neurons were frozen in 90% fetal bovine serum (FBS) (Thermo Fisher Scientific, Waltham, MA, USA) and 10% dimethyl sulfoxide (DMSO). Further maturation after thawing was driven by a growth factor cocktail. For the differentiation of PNN, doxycycline (2 µg/ml) exposure from DoD4’-9’ and DoD1-14 was integrated in the standard small molecule differentiation protocol, as detailed in the results chapter.

### PeriTox test

Immature peripheral neurons were thawed and used on DoD0 to assess the effects of test compounds on neurite area and cell viability (supplementary methods) as previously described [3, 4, 34, 35].

### Generation of a gene-edited iPSC line

The human Sigma iPSC0028 line was infected with the lentivirus described in figure 2A (also see supplementary methods). In brief, infected cells underwent hygromycin (Carl Roth, Karsruhe, Germany) selection followed by manual picking and expansion of the colonies. Stocks of the clones were cryopreserved in 90% FBS and 10% DMSO. Short tandem repeat (STR) DNA typing (described in detail in [36]) was performed for cell line authentication. To evaluate the clone’s NGN1 expression properties, iPSCs were seeded as single cells in E8 medium and exposed to doxycycline (2 µg/ml) for up to 5 days (Fig. 2B).

**Figure 2:**
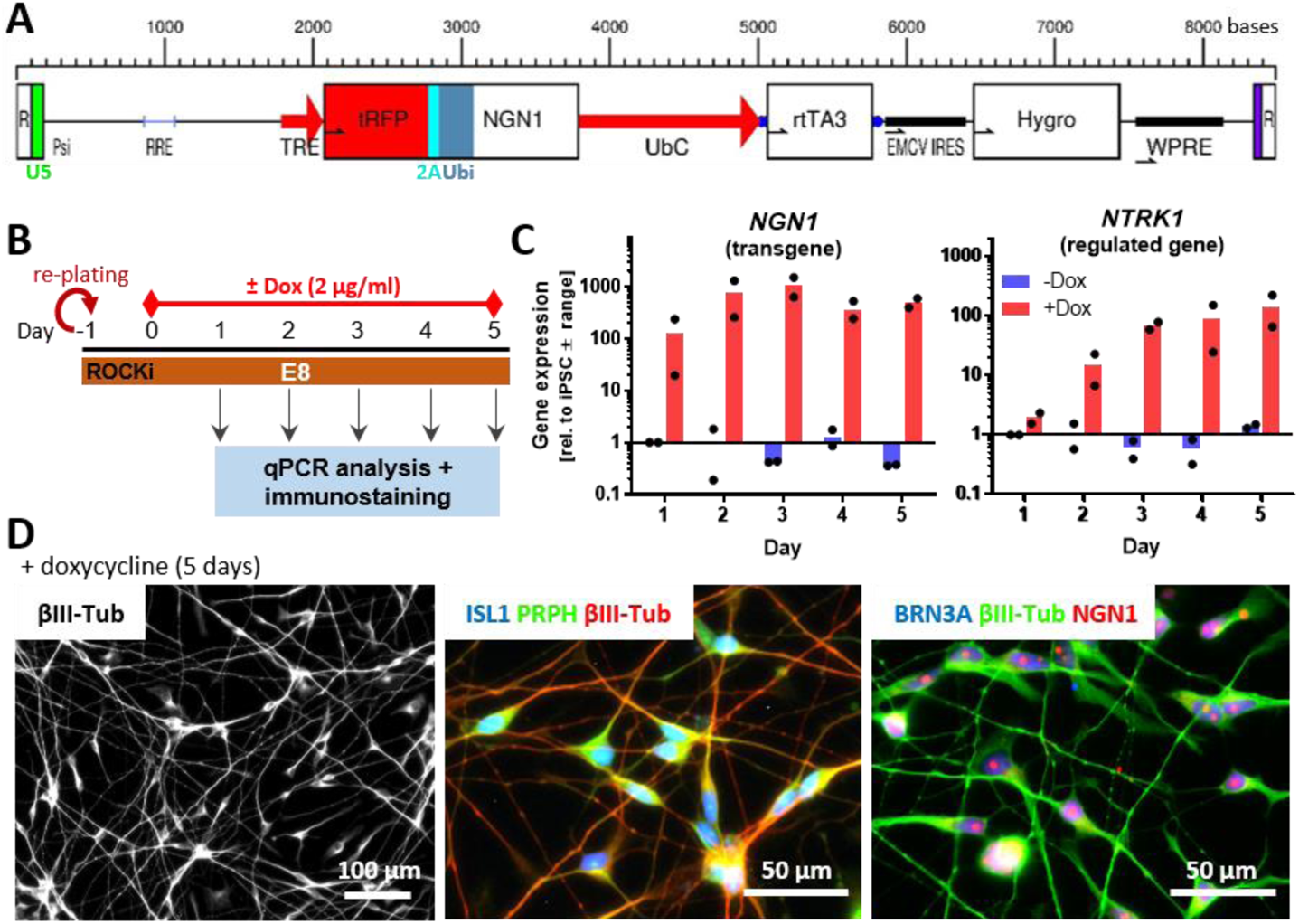
Generation of human iPSCs with inducible expression of ectopic NGN1. **(A)** Structure of the lentiviral construct used to generate a NGN1-overexpressing iPSC line. NGN1 is expressed as fusion protein with turbo red fluorescent protein (tRFP). tRFP and NGN1 are linked via a 2A region and ubiquitin (Ubi) to ensure the precise cleavage of NGN1 inside cells. Gene expression is controlled via the tetracycline-on system (TRE promotor). This system uses a reverse tetracycline-controlled transactivator (rtTA3) driven by the UbC promoter. R, repeat region of the HIV long terminal repeat (LTR) region; U5 (green), U5’ region of the HIV LTR; Psi, packaging sequence; RRE, Rev response element; TRE, tetracycline response element; UbC, Ubiquitin C promotor; EMCV IRES, encephalomyocarditis virus internal ribosomal entry site; Hygro, hygromycin resistance; WPRE, woodchuck hepatitis virus post-translational regulatory element. **(B)** Experimental setup to assess the NGN1 transgene expression. Cells were exposed for 5 days to doxycycline (0 or 2 µg/ml). **(C)** Gene expression of NGN1 and its downstream-regulated gene NTRK1 were monitored daily in control (blue) and doxycycline exposed cells. Gene expression was quantified by RT-qPCR. Data are given relative to iPSC (-Dox, day1). Data displayed as bars are means of two biological replicates (black dots). **(D)** Immunofluorescence images of cells treated with doxycycline for 5 days. Cells were labelled with antibodies against β-III tubulin (βIII-Tub), peripherin (PRPH) and the sensory neuronal transcription factors NGN1, ISL1 and BRN3A. Color code and scale bars are given in the images. More detail is given in figure S2E,F.

### Assessment of gene and protein expression

Gene expression was investigated by quantitative reverse transcriptase PCR (RT-qPCR) using SsoFast^TM^ EvaGreen® Supermix (Bio-Rad). Protein expression was assessed via immunofluorescence staining and microscopy. All samples were prepared, and analyzed exactly as described before [30, 37], using primers and antibodies as detailed in supplementary methods.

### Transcriptome data generation and analysis

Sample lysates were prepared as described [28, 30]. Measurements were performed at Bioclavis (BioSpyder Tech., Glasgow, UK) via the TempO-Seq targeted sequencing technology applied to the whole transcriptome set [38]. For data processing, the R package DESeq2 (v1.32.0) was used for quality control, normalization and determination of differentially expressed genes (DEGs) [39]. A Benjamini-Hochberg-adjusted threshold of P < 0.05 and a fold change of 2 were used as filter for DEGs. Analysis of gene ontology (GO) term over-represenation was done with g:profiler software [40]. All procedures are detailed in supplementary methods, and the data on numbers of reads for each gene analyzed, and the fold-changes for DEGs are provided in Supplement file1, organized as Excel workbook.

### Electrophysiological data

For electrophysiological characterization of the PNN, manual patch-clamp recordings and multielectrode array measurements were performed as described [29, 30]. Details are given in supplementary methods.

### Measurement of changes in intracellular Ca^2+^ concentration [Ca^2+^]i

Sensory neurons were cultured in 96-well plates after thawing. Cells were loaded with the Ca^2+^-indicator Fluo-4 (Thermo Fisher Scientific). Monitoring of [Ca^2+^]i was performed using a VTI HCS microscope (Thermo Fisher Scientific) equipped with an automated pipettor and an incubation chamber providing an atmosphere with 5% CO2 and 37°C. Cells were imaged for 45 s. Test compounds were automatically applied after baseline recording (10 s). The images were exported as *.avi video files and analysed with the CaFFEE software. Details are given in a dedicated technology paper [41] and in supplementary methods.

### Statistics

If not stated otherwise, experiments were performed on 3 or more independent cell preparations (here called biological replicates). In each cell preparation at least three different wells (here called technical replicates) were measured. Quantitative Ca^2+^-imaging data were derived from time-dependent series of images by using the CaFFEE software [41]. The binary endpoint of reactive/non-reactive cells was defined primarily by a well-specific, noise level-based threshold of changes in fluorescence intensity: (mean(ΔF) + 3x SD(ΔF)), with an upper limit set to 18 (ΔF: fluorescent change by negative control stimulation). Information concerning descriptive statistics and experimental variability is included in the figure legends or the figures themselves. GraphPad Prism 5 software (Version 7.04, Graphpad Software, Inc, San Diego, USA) was used for significance testing and data display. Data were evaluated by ANOVA plus appropriate post-hoc testing method or by t-test for binary comparisons. *p*-values < 0.05 were regarded as statistically significant.

## Results

### Characterization of human sensory neurons generated from non-modified iPSCs

For the generation of sensory neuronal cultures, we optimized a previously published two-step differentiation protocol starting from iPSC [4] (Fig. 1A). The time point of freezing of the cells was adapted (DoD9’), and the culture medium was supplemented with cytarabine from DoD3 until DoD14 to remove any mitotic, potentially non-neuronal cells. This procedure yielded pure neuronal cultures that develop an extensive neurite network (Fig. 1B, C). The capacity of such cells to grow neurites within 24 h forms the basis for the established PeriTox test [3, 4]. This assay was used to verify that typical neurotoxicants exhibit a specific neurite-damaging effect. The pesticide rotenone, the gout medication colchicine and the three chemotherapeutics taxol, bortezomib and oxaliplatin all reduced the neurite area at concentrations that did not affect general neuronal viability (Fig. 1D, S1B).

The sensory neuronal phenotype was confirmed by gene expression analysis. Markers like *PRPH*, *SCN9A* and *P2RX3* were expressed on DoD1 and further up-regulated over time. Indicators of the neural crest cell intermediary stage (*PAX3*, *TLX2*) were down-regulated (Fig. 1F). Markers for central neuron precursors (*PAX6*) or glial cells (*GFAP*) were absent. To investigate the functional expression of pain-related receptors, we used selective agonists of TRPV1 (capsaicin) and P2X3 (α,β-methylene ATP (α,β-meATP) [42]). Under normal conditions, only 10% of the cells showed P2X3- and 1% TRPV1-signaling. This was not considerably changed by maturation of up to 40 days (Fig. 1F). By mimicking inflammatory conditions with increased NGF concentrations [43–47] we obtained 60% of α,β-meATP-responsive cells. However, the capsaicin-responsive subpopulation did not exceed 5%. In summary, the optimized protocol generated largely pure, fully post-mitotic sensory neurons (Fig. S1A), but the functional properties were not suitable for CIPN research related to altered pain sensation, e.g., through the TRPV1 receptor system.

### Generation of an iPSC line with inducible NGN1 expression

The standard differentiation protocols did not yield a sufficiently large nociceptor subpopulation to allow functional studies. Consequently, we investigated an alternative approach. NGN1 is a key transcription factor in the development of the here-desired neurons [11, 12, 48]. Therefore, we hypothesized that its time-controlled overexpression would improve differentiation success [32].

An iPSC line was generated, in which NGN1 expression can be controlled by adding doxycycline to the medium. After the NGN1 expression construct (Fig. 2A, S2A) was stably inserted into the genome, the newly generated iPSC line Sigma-NGN1 was authenticated by the established method of STR analysis [36]. On this basis, the Sigma-NGN1 line and the commercially available Sigma iPSC 0028 line were declared identical (Fig. S3). The pluripotency of the newly generated iPSC population was assessed by immunofluorescence imaging. The expression of several pluripotency markers (e.g., Nanog, OCT4) (Fig. S2B-D) as well as the absence of the neuroectodermal markers PAX6 and SOX10 (data not shown) were similar to that of the pluripotent parent cell line. Further, the cells’ NGN1 expression properties were verified (Fig. 2B). Gene expression of *NGN1* was found to be inducible by doxycycline (Fig. 2C). The functionality of the *NGN1* transgene was derived from the control of its downstream target *NTRK1*. Moreover, transgene expression for five days leads to the complete conversion of iPSCs into cells expressing the pan-neuronal marker βIII-tubulin (βIII-Tub) and exhibiting neuronal morphology. Furthermore, these cells expressed the PNS markers peripherin (PRPH), BRN3A and ISL1 (Fig. 2D, S2E,F)). Such a staining pattern is typical for neurons that have exited the cell cycle [11]. Taken together, these data confirm the successful generation of an iPSC line carrying an inducible *NGN1* transgene with the expected functionality of the gene product, NGN1.

### Integration of NGN1 overexpression in the standard small molecule differentiation protocol

In a next step, it was tested, which time window of NGN1-overexpression was most suitable to improve the standard differentiation protocol. Three different doxycycline exposure schedules (S1– S3), integrated into the standard differentiation, were investigated (Fig. 3A). Gene expression of the sensory neuronal markers *NGN1*, *NTRK1*, *RUNX1*, *PRPH*, *POU4F1* (*BRN3A*) and *ISL1* was monitored daily until the day of freezing (Fig. 3B, S4B). Early induction of *NGN1* expression on DoD2’ in condition S2 led to an earlier expression of *NGN1*, *NTRK1*, *PRPH* and *ISL1* compared to cells not exposed to doxycycline (S1). However, up-regulation of *RUNX1* expression, a gene crucial for nociceptor specification [48, 49], was poor, while the rate of cell death after thawing was increased (Fig. 3C). Differentiation condition S3 resulted in the highest gene expression levels for *RUNX1*, and shifted *PRPH* and *NTRK1* expression to earlier time points. Immunofluorescence images on DoD3 showed that all three exposure schedules yielded peripheral neurons (PRPH^+^ neurites). The sensory neuronal markers ISL1 and BRN3A were expressed to a large extent in conditions S1 and S3 (74-100% positive cells), but not in S2 (Fig. 3C, S4A,C,D), which was decisive to exclude S2. Eight days after thawing uniformity of neuronal cultures was further assayed using tRFP fluorescence as an internal reporter of NGN1 expression (Fig. 3D). Quantification of red fluorescent cells showed S3-derived cultures to be more uniform (96% tRFP-positive cells) than S1 cultures (36% tRFP-positive cells) (Fig. 3E). Therefore, all future experiments were conducted using exposure schedule S3, which yields neuronal cultures with the highest sensory neuron marker expression and uniformity.

**Figure 3:**
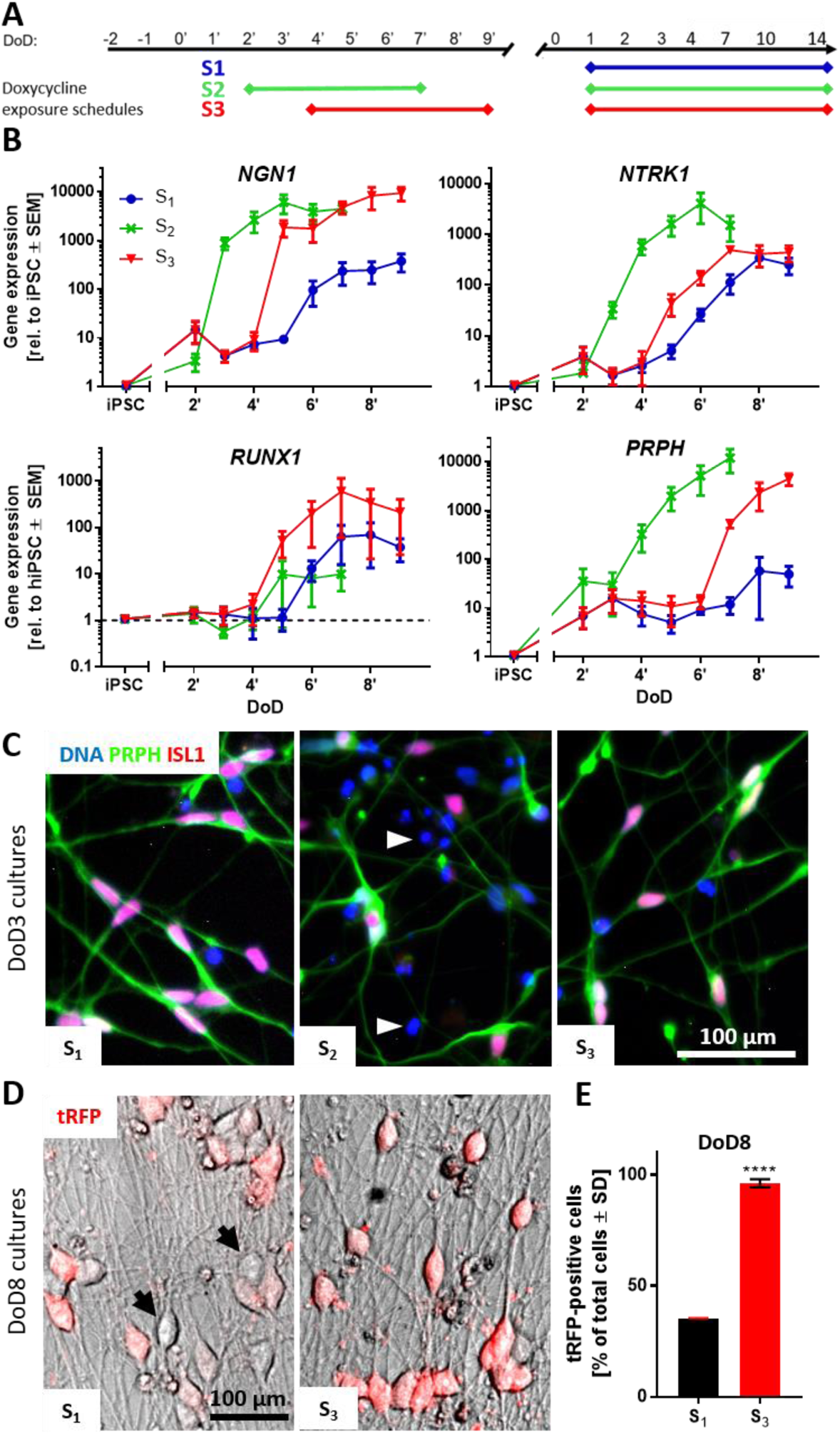
Integration of NGN1 overexpression in the standard small molecule differentiation protocol. **(A)** Schematic representation of three different doxycycline exposure schedules (S1, S2, S3) incorporated in the standard differentiation protocol. Condition S1 did not receive doxycycline treatment before freezing. Cells of condition S2 were treated with doxycycline from DoD2’ until DoD7’ with subsequent freezing of the cells. Condition S3 included doxycycline treatment from DoD4’ until DoD9’ with subsequent freezing. After thawing, all three conditions were exposed to doxycycline from DoD1 until DoD14. **(B)** Gene expression analysis of the nociceptor marker genes NGN1, NTRK1, RUNX1 and the general sensory neuronal marker gene peripherin (PRPH) for all three exposure situations. Data are expressed relative to expression levels in iPSCs and given as means ± SEM of 3-4 biological replicates. **(C)** Immunofluorescence images of cultures (S1-3) on DoD3 (after thawing). Cells were stained for peripherin (PRPH) and ISL1. Nuclei were stained with H33342 (blue). Details are displayed in figure S4A. White arrowheads indicate exemplary dead cells. **(D)** Overlay of phase contrast and tRFP fluorescence images of condition S1 and S3 neuronal cultures on DoD8. Black arrows indicate exemplary tRFP-negative cells. **(E)** Quantification of tRFP positive cells in cultures of differentiation conditions S1 and S3 on DoD8. Data are shown as percentage of total cell count ± SD. DoD, day of differentiation; tRFP, turbo red fluorescent protein.

### Transcriptomics-based characterization of mature iPSC-derived sensory neurons

We used time-dependent transcriptome profiling as broad and unbiased approach to describe the differentiation process of iPSC-derived sensory neurons. Transcript levels of about 19,000 genes were measured for 7 differentiation stages (Suppl. File2). A principal component analysis (PCA) was used as a first overview of the data structure. Independent biological replicates clustered closely together, and the first principle component coincided with increasing time of maturation (Fig. 4A). The absolute expression levels of the sensory neuron marker genes *ISL1* and *PRPH* were high (>1000 transcripts per 1 million reads) from DoD1 until DoD49. Nociceptor marker genes like *P2X3*, *RET* and *SCN9A* also reached high absolute levels (Suppl. File2). However, some essential genes (*NTRK1*, *SCN10A*, *TRPV1)* were not captured well by the transcriptome mapping approach. Consequently, their gene expression was investigated via RT-qPCR. The levels of *NGN1*, *RET*, *NTRK1*, *RUNX1* and *SCN10A* peaked at DoD3-7 and then declined until DoD21 (Fig. 4B). For *P2RX3*, *SCN9A*, *TRPM8* and *TRPV1* we found increased expression on DoD3-7, and thereafter largely stable levels until DoD21 (Fig. 4C).

**Figure 4:**
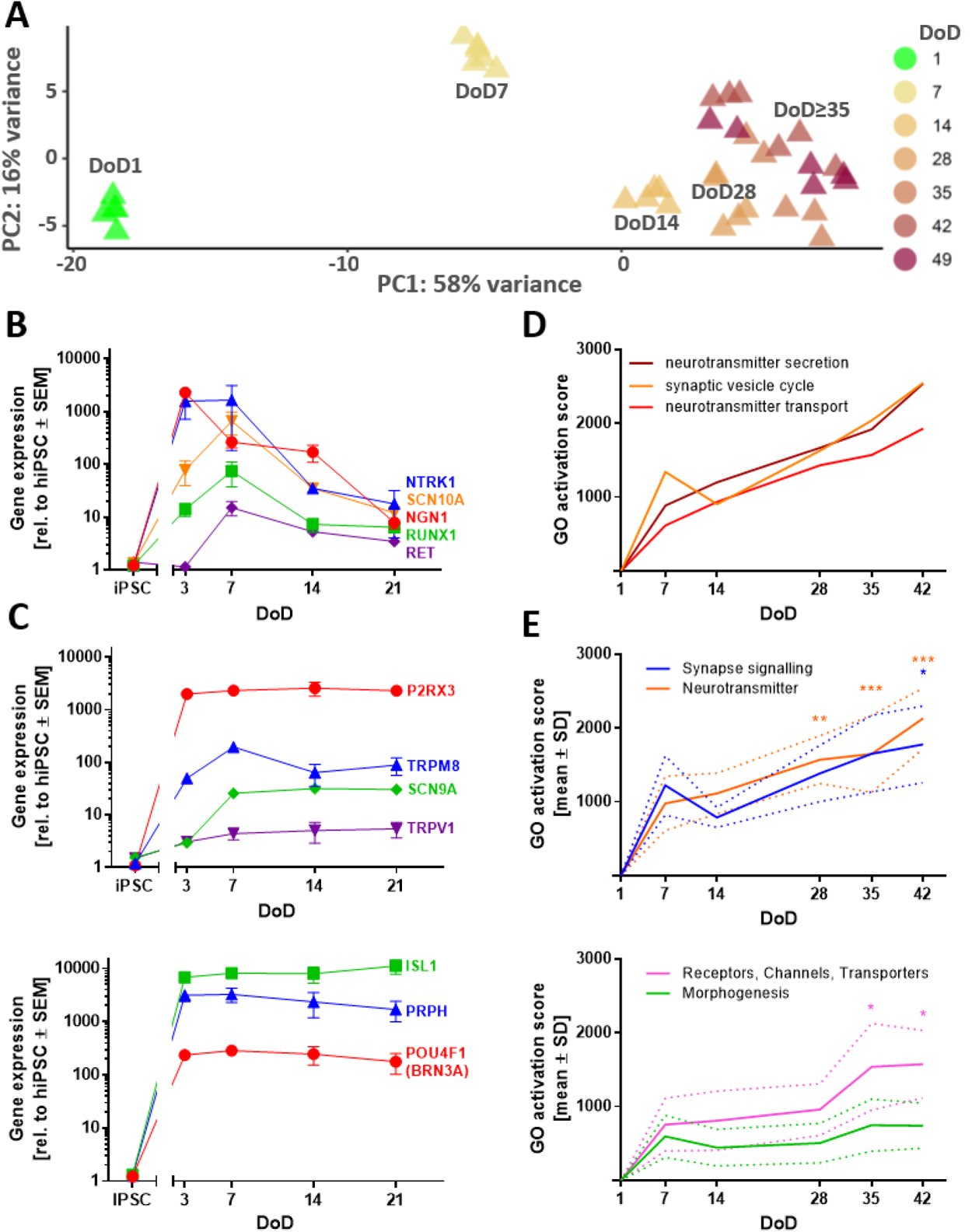
Time-dependent transcriptome profiling of iPSC-derived sensory neurons. **(A)** TempO-Seq whole transcriptome analysis (19,000 genes). For the top 500 variable genes of this data set (full data in Supplementary file1) a PCA was performed. In the two-dimensional PCA display, seven maturation stages of PNN are color-coded according to their DoD. Data points are derived from three independent differentiations. **(B,C)** Gene expression levels of sensory neuron and nociceptor marker genes were assessed via RT-qPCR. Data are means ± SEM, n = 3-4. Error bars smaller than the data point symbols are not shown. **(D)** Over-represented gene ontology (oGO) terms were determined for the significant DEGs on DoD42. Quantitative activation scores of the Top50 oGOs were calculated for all time points by “multiplying the percentage of genes within the GO that was found to be significantly regulated with the average fold change of these regulations” [51]. Activation scores for the GO terms “neurotransmitter secretion”, “synaptic vesicle cycle” and “neurotransmitter transport” are shown over time. **(E)** oGO terms were assigned to the superordinate groups “Synapse signaling”, “Neurotransmitter”, “Receptors, Channels, Transporters” and “Morphogenesis” (see Fig. S5). Means of the activation scores of all oGOs belonging to one group are shown to visualize the development of these biological categories over time. The dotted lines indicate the upper and lower bounds of the SEM. Significance was tested against the respective mean activation scores on DoD7. * p < 0.05, ** p < 0.001, *** p < 0.0001.

The expression kinetics of these pre-selected transcripts are in good agreement with our objective of generating nociceptor-enriched sensory neuron cultures. For further transcriptome data mining, DEGs were determined for all sampling time points (Suppl. file2). For the 600 DEGs of DoD42 altogether 200 over-represented gene ontology (oGO) term groups were identified [40]. The 50 oGOs with the lowest *p-*values mainly fell into the superordinate groups “synapse signaling”, “neurotransmitters”, “receptors, channels, transporters” and “morphogenesis”. They were also analyzed for the other time points (Fig. S5A,B) and quantitative GO activation scores [50, 51] were calculated for all time points (Fig. 4D, S5C). Activation scores for “synapse signaling” and “neurotransmitters” showed a continuous increase until DoD42 (Fig. 4E). The activation scores of receptor/channel-related genes showed a plateau for DoD7-28 and then jumped to a higher level at late differentiation stages (DoD35-42) (Fig. 4F). In summary, analysis of gene expression patterns over large biological categories confirmed that the here-established differentiation protocol yields peripheral neurons with nociceptor features (PNN). While most general neuronal markers were well established after 1-3 weeks of differentiation, genes linked to particular PNN functions continued to be up-regulated until at least DoD35-42.

### Electrophysiological characterization of PNN

The basic functional characterization of the PNN also included a check for general neuronal electrophysiological features. Patch-clamp measurements provided evidence for all major classes of voltage-gated cation channels (KV, NaV and CaV) (S7). All cells recorded showed that they could fire action potentials (Fig. 5A). Half of the cells showed a phasic firing pattern (Fig. 5B, left), while the other half displayed tonic firing behavior (Fig. 5B, right). This distribution is consistent with the current literature on the characterization of primary rat DRG neurons [52]. After confirmation of these basic neuronal properties, we moved on to establish a neuronal signaling endpoint, more suitable for broader toxicological/pharmacological evaluation.

**Figure 5:**
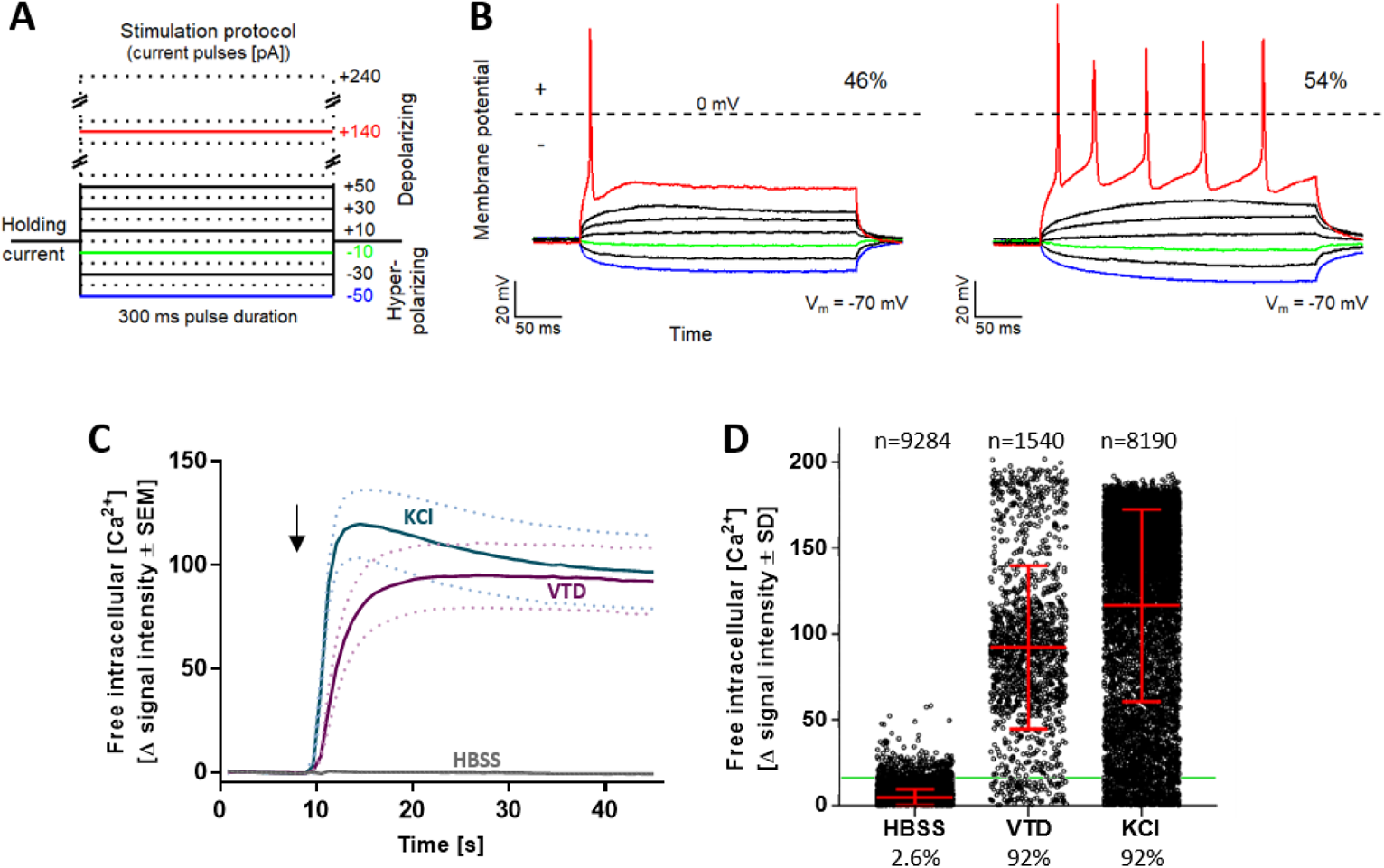
Characterization of neuronal excitability of PNN. PNN (DoD28-35) were used for current-clamp recordings and Ca^2+^-imaging experiments. **(A)** Schematic representation of the stimulation protocol used for current-clamp recordings. Current pulses were applied with a pulse duration of 300 ms and a pulse frequency of 0.2 Hz, starting at -50 pA and increasing in steps of +10 pA. Example pulses are colored. **(B)** Current-clamp measurements of PNN (n=28). Cells exhibit phasic (46%) (B, left) or tonic (54%) (B, right) action potential firing behavior. **(C)** Representative traces of changes in Ca^2+^ indicator fluorescence (=Δ signal intensity) in response to the negative control HBSS (Hanks’ balanced salt solution), the positive control KCl [50 mM] and the voltage-gated sodium (NaV) channel opener veratridine (VTD) [3 µM]. The arrow indicates the time point of stimulus addition. Data are shown as means of 4 biological replicates. The dotted lines indicate the upper and lower bounds of the SEM. **(D)** Quantification of the percentage of reactive cells according to their Δ signal intensity values upon HBSS, VTD or KCl addition. The green line indicates the noise boundary of the Δ signal intensity. Each dot represents the Δ signal intensity of an individual cell. The mean ± SD of all cells is shown graphically in red. The percentage of reactive cells is indicated below the diagram and the exact number of measured cells is given above. A total of more than 10,000 cells was individually measured in 14 experiments.

### Establishment of intracellular Ca^2+^-measurement as test endpoint

We decided on the use of Ca^2+^-imaging [53] as signaling endpoint for our PNN. General proof-of-concept for the feasibility of this approach was obtained by recording strong signals triggered by increased K^+^ concentrations in the medium or by opening of NaV channels by veratridine (VTD) (Fig. 5C,D). As PNN are a mixed neuronal population, it was important to establish the Ca^2+^-signaling endpoint on a single cell level. As practical approach to work with the multi-dimensional information provided by the recording of Ca^2+^ fluorescence time courses of thousands of cells, we decided to use a binary endpoint of “responsive” *versus* “non-responsive” cells. For this, we thoroughly investigated and defined suitable response thresholds (Fig. S8). Based on the extensive evaluation (signal intensity changes (Δ) for >9200 cells), a robust algorithm was chosen to define responsive cells in Ca^2+^-signaling experiments.

### Functional characterization of PNN cultures regarding nociceptive features

A hallmark of nociceptive neurons is the expression of ion channels responsible for the sensation of pain. In this study, we focused on TRPV1 and P2X3. Immunofluorescence staining revealed the presence of both receptors in virtually all cells on DoD42 (Fig. 6A, S9A,B). The specific agonist of P2X3 receptors α,β-meATP and the TRPV1-agonist capsaicin were used as tool compounds for functional characterization. The cultures did not react to the stimuli during the first 4 weeks after thawing. From then on, the percentage of reactive cells continuously increased until DoD42 (Fig. S9C, left, right). For comparison, functional NaV channels were found to be present from DoD7 on and maximum culture responsiveness towards the NaV opener VTD was reached on DoD21 (Fig. S9C, middle).

**Figure 6:**
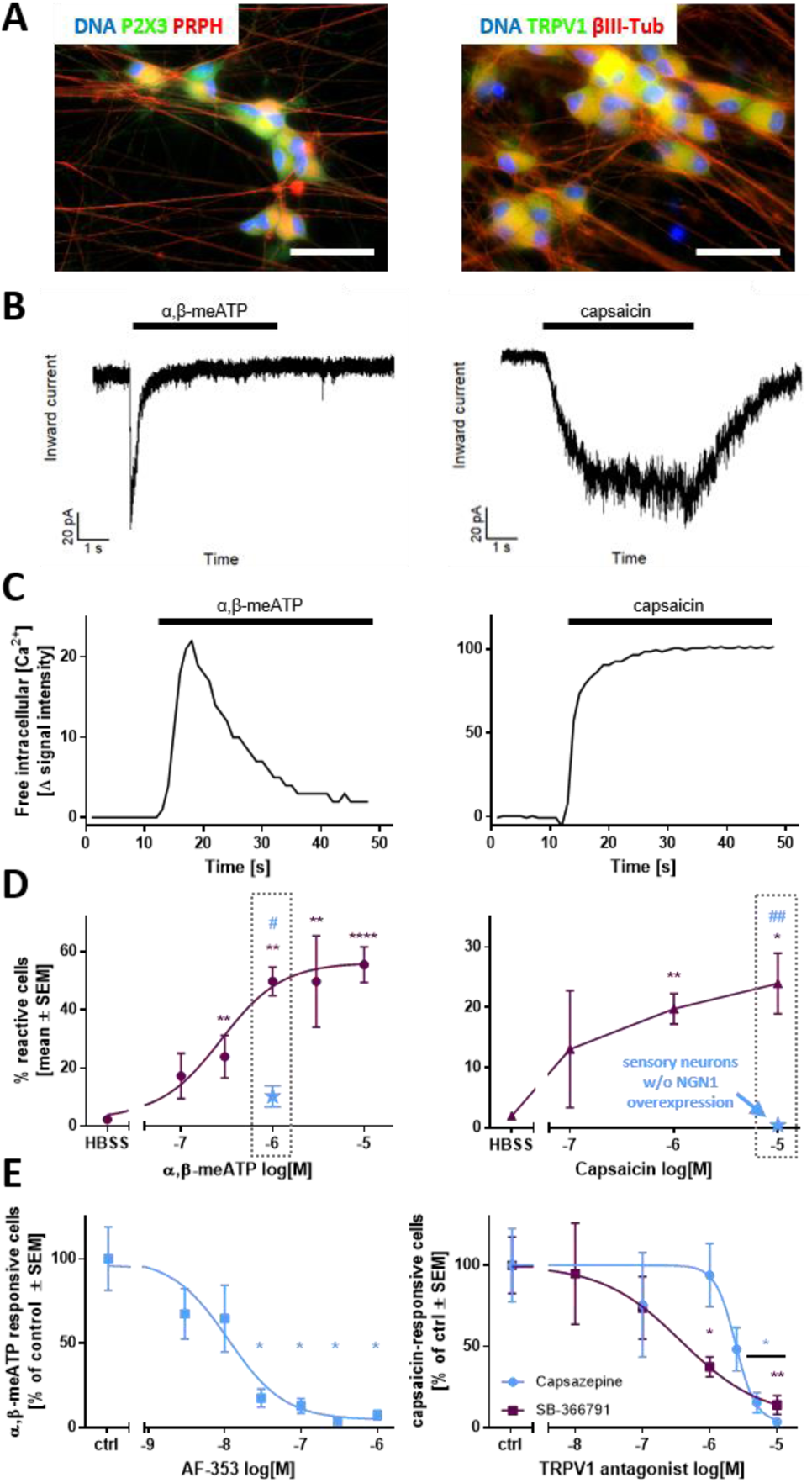
Functional characterization of the nociceptor-ion channels P2X3 and TRPV1 in PNN. Sensory neurons were differentiated for at least 37 days after thawing. **(A)** Representative immunofluorescence images of fixed cells stained for P2X3 and peripherin (PRPH) (left), as well as TRPV1 and βIII-tubulin (βIII-Tub) (right). Nuclei were stained using H33342 (DNA). Color code and scale bars are given in the images, and further details are shown in figure S9A,B. **(B)** Representative voltage-clamp recordings of cells exposed to α,β-methylene ATP (α,β-meATP) [10 µM] (left) and capsaicin [1 µM] (right). The black bars indicate the time of compound exposure. **(C)** Representative traces of changes in intracellular Ca^2+^ concentration of single cells upon addition of α,β-meATP [1µM] (left) and capsaicin [1 µM] (right). The black bars indicate the time of compound exposure. **(D)** Concentration-dependency of the percentage of reactive cells towards a stimulus of α,β-meATP (left) and capsaicin (right). The blue data point depicts the respective percentage of reactive cells in sensory neurons generated traditionally (without transient NGN1 overexpression). **(E)** Concentration-dependency of the P2X3-specific antagonist AF-353 (left) and the TRPV1-specific antagonists capsazepine (blue) and SB-366791 (purple) (right). All data are means ± SEM of at least 3 biological replicates. Significance was tested against control/HBSS (*) or against cells w/o NGN1 overexpression (#). */# *p* < 0.05, **/## *p* < 0.005.

Responses induced by α,β-meATP were characterized by fast-inactivating inward currents typical for P2X3 receptors (Fig. 6B,C, left) [54–56]. Capsaicin, in contrast, evoked sustained inward currents throughout the exposure period, as is typical for TRPV1 receptors (Fig. 6B,C, right) [57, 58]. The expression of functional TRPV1 receptors was further substantiated, as treatment with two other TRPV1-agonists, olvanil and piperine, induced Ca^2+^ influx in a subset of neurons (Fig. S9G). Quantification of reactive cells revealed a concentration-dependency of both P2X3 and TRPV1 responses (Fig. 6E), which also makes this endpoint a useful model for pharmacological intervention studies in PNN. The reactivity of PNN towards nociceptive stimuli was clearly superior to the one observed in peripheral neurons differentiated conventionally (without transient NGN1 overexpression) (Fig. 6D, grey boxes).

As a next step, we performed double-stimulation studies to investigate the overlap of P2X3 and TRPV1 receptor-expressing cell populations (Fig. S9D). PNN were treated with α,β-meATP followed by a capsaicin stimulus and vice versa. Independent of the sequence, we found that 40% of the cells reacted to a P2X3 stimulus only, while about 10% reacted towards capsaicin only. One quarter of the whole population responded to both stimuli (Fig. S9E). These sequential stimulation experiments also demonstrated that there was no cross-(de)sensitization of TRPV1 and ATP receptors, as has sometimes been claimed [59–61]. This finding also significantly increased the throughput of this method, as double-stimulations can be used as the standard experimental design.

To ensure that the measured responses are P2X3- and TRPV1-specific, the cells were pre-incubated with the P2X3-selective antagonists AF-353 (Fig. 6E, left) or A-317491 (data not shown). A concentration-dependent decrease in Ca^2+^ influx was observed at ≥30 nM AF-353 and ≥2.5 µM A-317491, confirming P2X3 as the main P2X subtype expressed in PNN [62, 63]. To prove the specificity of TRPV1-responses the well-known antagonist capsazepine was used. We also tested SB-366791, which exhibits improved selectivity and potency [64, 65]. Both antagonists blunted the capsaicin responses. Selectivity of the receptor-antagonists A-317491 and capsazepine was confirmed by double-stimulation experiments demonstrating that only the respective target receptor was inhibited, but not the response of the other receptor (Fig. S9F).

### Modelling CIPN-related alterations in pain receptor functions *using PNN*

Acute painful CIPN is often attributable to alterations in pain signaling, but not necessarily to morphological damage. We performed here an oxaliplatin case study to investigate the potential of PNN to model acute chemotherapy-related functional alterations *in vitro*. On the basis of the PeriTox test (DoD0 cells) [4], two oxaliplatin test concentrations were selected. This assay has been used broadly for identifying neurite-damaging agents [3, 34]. Based on the test data (Fig. S10A), we selected 5 µM (no effect) and 20 µM (moderately decreased neurite area, but no cell death) for further experiments.

In PNN, matured for several weeks, neither concentration affected the neurite integrity or viability (Fig. S10B). Basic neuronal function (spontaneous firing) was also maintained (Fig. S10C). Using Ca^2+^-signaling as endpoint, we examined whether pain-related excitability was affected independent of morphological effects. First, we established a simple model of mechanical allodynia. Pre-treatment with 20 µM oxaliplatin (24 h) made PNN react to a mechanical stimulus (mild shear forces) with increased Ca^2+^ influx (Fig. 7A,B, Fig. S11A,B). As NaV channels have been implied in OXAIPN-mechanical allodynia [66], we applied the inhibitors TTX and carbamazepine (Carb). They fully blocked Ca^2+^ signals following mechanical stress (Fig. 7B). This dampening effect was specific for the mechanical stress model, as the same inhibitors did not affect signaling triggered by direct TRPV1 activation (Fig. S11D). P2X3 is not involved in this *in vitro* mechanical allodynia, as inhibition by A-317491 had no effect (Fig. 7B). We were interested in learning, whether the bare presence of oxaliplatin is sufficient to alter neuronal responsiveness (allodynia). The washout of oxaliplatin did not restore normal functions and a shortened pre-treatment time (1 h) did not lead to the same de-regulations as observed with 24 h incubation time (Fig. 7C,D). These data argue against a direct interaction of oxaliplatin with NaV channels as a cause for the observed signaling changes.

**Figure 7:**
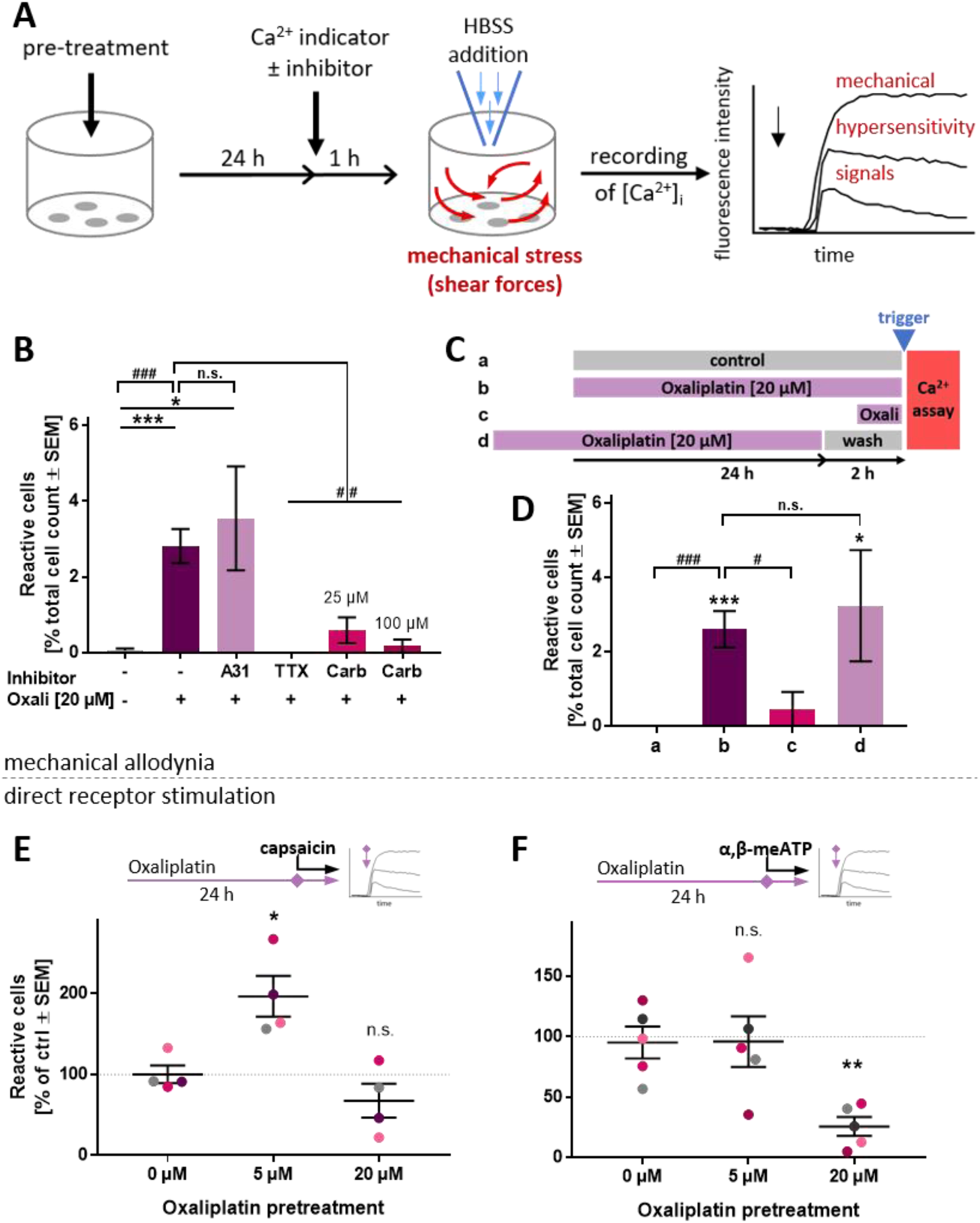
Functional impairment of PNN exposed to oxaliplatin. PNN (>DoD37) were used for Ca^2+^-imaging experiments to assess oxaliplatin-induced functional impairments. **(A)** Schematic representation of the experimental setup to assess responses towards a mechanical stimulus. **(B)** PNN were pre-treated with oxaliplatin (Oxali) according to **(A)**. Cells received the P2X3 antagonist A-317491 (A31, 10 µM), or the NaV channel inhibitors tetrodotoxin (TTX, 3 µM) or carbamazepine (Carb). The fractions of cells reacting with Ca^2+^-influx (ΔF > 18) were quantified. **(C)** Schematic representation of oxaliplatin exposure scenarios **a**-**d** before stimulation of cells by HBSS addition (see trigger mark). Some cells (**a**) did not receive oxaliplatin pre-treatment, while others (**b**) were pre-treated with oxaliplatin for 24 h prior to Ca^2+^ measurements. Condition **c** included pre-treatment with oxaliplatin for 1 h only. In scenario **d,** a 24 h oxaliplatin pre-treatment was ended 2 h before Ca^2+^-measurement. **(D)** PNN were treated with oxaliplatin according to the scenarios **a**-**d**, and reactive cells were quantified. Data in B, D are means ± SEM, from 3-6 biological replicates. Significance was tested against condition **a** (*) or against condition **b** (#). */# *p* < 0.05, ***/### *p* < 0.001. **(E, F)** Effect of oxaliplatin pre-treatment (24 h) on **(E)** TRPV1 and **(F)** P2X3 PNN reactivity upon capsaicin or α,β-meATP stimulation (both 1 µM), respectively. Data points belonging to the same cell lot are color-matched. Means ± SEM are given. Significance was tested against the respective control (0 µM). * *p* < 0.05, ** *p* < 0.005, n.s. not significant.

As second approach to understand functional impairments triggered by oxaliplatin, we studied potentially modified receptor responses. Triggered by the pertinent literature [17, 22] we focused on TRPV1. A significant hypersensitivity to capsaicin was observed at low oxaliplatin (5 µM) concentrations (Fig. 7E). The control stimulus (P2X3 receptors) was not affected at this concentration. However, the response of P2X3 receptors was found to be decreased at higher (20 µM) oxaliplatin pre-treatment (Fig. 7F). As the depression of P2X3 responses by oxaliplatin pre-treatment was quite pronounced, and some direct receptor inactivation by oxaliplatin may be conceived, we performed a series of experiments varying the presence of the chemotherapeutic drug during receptor stimulation. Only prolonged pre-treatment was effective, while direct presence of oxaliplatin was not required to attenuate P2X3 responses (Fig. S11C).

In contrast to oxaliplatin, cisplatin treatment usually is not associated with acute pain effects [9]. We investigated whether this was replicable in the *in vitro* model. In cisplatin pre-treated cells, we did neither observe mechanical allodynia-like signals nor TRPV1 hyper-responsiveness. However, a decrease in P2X3 responsiveness (Fig. S11A,E) was observed as seen similarly for oxaliplatin.

In summary, these data suggest that CIPN-relevant alterations of ion channel functions can be observed and studied in PNN. As both hyper- and hypo-sensitivity to different stimuli can be simultaneously assessed in a concentration-dependent manner, the PNN-based test system allows for novel approaches to study CIPN *in vitro*.

## Discussion

We present here a robust method to generate PNN. Moreover, the study provides a full characterization of a signaling endpoint that can be used to assess normal and disturbed neuronal signaling in such cultures. Finally, we demonstrate in an exemplary case study the applicability of PNN to assess pain-related altered neuronal excitability after exposure to a chemotherapeutic drug.

Altogether, this paper contains Ca^2+^-signaling data for more than 60,000 individual neurons. The recording of intracellular Ca^2+^ concentrations over time provides a wealth of data (considering different curve shapes, peak heights, areas-under-the curve, relaxations times etc.). Extraction of robust information from such multidimensional data sets can be extremely difficult. Often it is not possible at all, unless the evaluation method is adapted and optimized from experiment to experiment. The latter procedure has a relatively large risk-of-bias. We explored here as alternative the use of a binary readout of “responsive” *versus* “non-responsive” cells. This allowed the clear and accessible display of the multidimensional data recorded by high-throughput imaging. With this test method in hand, we demonstrated, using the example of oxaliplatin, that drug-induced receptor hyper-sensitivity or mechanical allodynia can be assessed *in vitro*.

Until few years ago, the major models to study peripheral pain-related neuropathies were experimental animals and patients [1, 67]. The few *in vitro* studies mainly focused on structural defects, and the main test systems for this were rodent neurons. Robust quantitative studies on nociceptor modulation are thus quite limited [68]. *In vivo* studies often assess behavioral outcome measures that are the result of a complex integration of peripheral, central, and glial cell type activities. In such situations, specific mechanisms or receptors are hard to assess. Within the published mechanistic studies, only a small fraction focused on functional neuronal properties [69]. Morphology-based test methods are more wide-spread and better-established, but they may miss signaling changes [28]. This is at present an important gap in CIPN-research, as it is known that chemotherapeutic drugs like oxaliplatin can alter neuronal excitability/function without structural damage [25]. Test systems based on nociceptor functions are therefore required in this area.

Since pluripotent stem cells have been established as readily-available resources, it became possible to generate complex human cell types not easily available from other sources. Protocols to generate peripheral neurons from iPSCs have paved the way for new human-relevant test systems. *In vitro* test methods offer many advantages for the study of specific mechanistic and pharmacological aspects of peripheral neurotoxicity due to their relative “simplicity”, and as environmental factors can be very tightly controlled. Indeed, iPSC-based *in vitro* test methods have been repeatedly used to measure the effects of chemotherapeutics on cellular viability or morphology [3, 4, 27, 70–72]. However, there is still a dearth of studies that employ human iPSC-derived nociceptor cultures to assess alterations of signaling endpoints. The cell system, together with the endpoints we characterized may help to fill this gap for toxicological or pharmacological studies.

Although we have several years of experience in the use of sensory neuronal cultures [3, 4, 30, 73], it was not possible to significantly improve the nociceptor character of these cells by using modifications of conventional protocols [2]. Functional sensory neurons can indeed be obtained after differentiation times of > 60 days [32, 74]. However, such time-demanding protocols severely limit the usefulness and robustness of the resulting cultures. Therefore, we harnessed here the nowadays widely used method of transcriptional programming [32, 75–78] to enhance the fate specification towards nociceptive neurons by transient NGN1 overexpression. The integration of this approach into the traditional small molecule differentiation protocol yielded PNN with a high abundance of P2X3 and TRPV1 receptors. It was interesting to note that gene expression patterns quickly resembled those of PNN, but the cells required considerably more time to acquire functional properties of nociceptors. For instance, high transcript levels for P2X3 were detected already on DoD1, while responses to P2X3 agonists were measured earliest from DoD28 onwards. This may be attributable to the continuing changes in several components of signaling pathways [79]. Our study therefore also demonstrates that the expression of receptor-encoding genes does not necessarily imply the functionality of these receptors.

We made here use of the fact that recording of Ca^2+^-signaling on the level of single cells allows the assessment of the composition of functionally heterogeneous populations. This is important for nociceptors, which are known to be a phenotypically mixed population (e.g., TRPV1 expression is only found on half of the peptidergic nociceptors [80]). Mixed populations also require large numbers of cells to be monitored to obtain robust results. In this sense, the endpoint presented here offers possibilities to re-evaluate other approaches that originally had to use < 20 individual cells for important statements [22].

## Conclusion

The *in vitro* model of PNN opens new possibilities for the study of functional aspects of peripheral neuropathies. However, our study does not close all gaps. An important future goal is the further shortening of the culture time, and the generation of several, highly defined subpopulations of sensory neurons. For instance, nociceptors expressing TRPA1 [17], or other specific receptors and ion channels would be desirable. It should also not be forgotten that toxicity often is a network phenomenon that may involve interactions between several glial cells, nociceptors and central neurons [81, 82]. Next steps might therefore involve co-culturing of PNN with e.g., Schwann cells [83]. In the more distant future, it is likely that network recording tools, like multi-electrode arrays, will reach cellular resolution, and thus allow recording of single cell responses in mixed cultures.

## Abbreviations

α,β-meATP: α,β-methylene ATP
βIII-Tub: βIII-tubulin
AraC: cytarabine
ATP: adenosine triphosphate
BDNF: brain-derived neurotrophic factor
CIPN: chemotherapy-induced peripheral neuropathy
CNS: central nervous system
CPM: counts per million
DEG: differentially expressed gene
DMSO: dimethyl sulfoxide
DRG: dorsal root ganglion
FBS: fetal bovine serum
FGF: fibroblast growth factor
GDNF: glia-derived neurotrophic factor
GO: gene ontology
HBSS: Hanks’ Balanced Salt Solution
iPSC: induced pluripotent stem cell
NGF: nerve growth factor
oGO: over-represented gene ontology
OXAIPN: oxaliplatin-induced peripheral neuropathy
PNN: peripheral neurons with nociceptor features
PNS: peripheral nervous system
PRPH: peripherin
RT-qPCR: quantitative reverse transcriptase polymerase chain reaction
ROCKi: ROCK inhibitor
tRFP: turbo red fluorescent protein
TRP: transient receptor potential
TRK: tyrosine receptor kinase

## Acknowledgements

This work was supported by CEFIC, the BMBF, EFSA, and the DK-EPA (MST-667-00205). It has received funding from European Union’s Horizon 2020 research and innovation program under grant agreements No. 964537 (RISK-HUNT3R), No. 964518 (ToxFree) and No. 825759 (ENDpoiNTs). This work received financial support from the State Ministry of Baden-Wuerttemberg for Economic Affairs, Labour and Tourism.

## Data Availability Statement

Raw data can be requested from the corresponding author.

## References

1. Lehmann HC, Staff NP, Hoke A (2020) Modeling chemotherapy induced peripheral neuropathy (CIPN) in vitro: Prospects and limitations. Exp Neurol 326, 113140.

2. Chambers SM, Qi Y, Mica Y et al. (2012) Combined small-molecule inhibition accelerates developmental timing and converts human pluripotent stem cells into nociceptors. Nat Biotechnol 30, 715–720.

3. Delp J, Gutbier S, Klima S et al. (2018) A high-throughput approach to identify specific neurotoxicants/ developmental toxicants in human neuronal cell function assays. ALTEX 35, 235– 253.

4. Hoelting L, Klima S, Karreman C et al. (2016) Stem Cell-Derived Immature Human Dorsal Root Ganglia Neurons to Identify Peripheral Neurotoxicants. Stem Cells Transl Med 5, 476–487.

5. Campbell JN, Meyer RA (2006) Mechanisms of neuropathic pain. Neuron 52, 77–92.

6. Boivie J, Leijon G, Johansson I (1989) Central post-stroke pain — a study of the mechanisms through analyses of the sensory abnormalities. Pain 37, 173–185.

7. Basbaum AI, Bautista DM, Scherrer G et al. (2009) Cellular and molecular mechanisms of pain. Cell 139, 267–284.

8. Shah A, Hoffman EM, Mauermann ML et al. (2018) Incidence and disease burden of chemotherapy-induced peripheral neuropathy in a population-based cohort. J Neurol Neurosurg Psychiatry 89, 636–641.

9. Staff NP, Grisold A, Grisold W et al. (2017) Chemotherapy-induced peripheral neuropathy: A current review. Ann Neurol 81, 772–781.

10. Snider WD, McMahon SB (1998) Tackling Pain at the Source: New Ideas about Nociceptors. Neuron 20, 629–632.

11. Ma Q, Fode C, Guillemot F et al. (1999) Neurogenin1 and neurogenin2 control two distinct waves of neurogenesis in developing dorsal root ganglia. Genes Dev 13, 1717–1728.

12. Ma Q, Chen Z, Del Barrantes IB et al. (1998) neurogenin1 Is Essential for the Determination of Neuronal Precursors for Proximal Cranial Sensory Ganglia. Neuron 20, 469–482.

13. Caterina MJ, Schumacher MA, Tominaga M et al. (1997) The capsaicin receptor: a heat-activated ion channel in the pain pathway. Nature 389, 816–824.

14. Chen CC, Akopian AN, Sivilotti L et al. (1995) A P2X purinoceptor expressed by a subset of sensory neurons. Nature 377, 428–431.

15. Cook SP, Vulchanova L, Hargreaves KM et al. (1997) Distinct ATP receptors on pain-sensing and stretch-sensing neurons. Nature 387, 505–508.

16. Immke DC, Gavva NR (2006) The TRPV1 receptor and nociception. Semin Cell Dev Biol 17, 582–591.

17. Calls A, Carozzi V, Navarro X et al. (2020) Pathogenesis of platinum-induced peripheral neurotoxicity: Insights from preclinical studies. Exp Neurol 325, 113141.

18. Adelsberger H, Quasthoff S, Grosskreutz J et al. (2000) The chemotherapeutic oxaliplatin alters voltage-gated Na+ channel kinetics on rat sensory neurons. Eur J Pharmacol 406, 25–32.

19. Wilson RH, Lehky T, Thomas RR et al. (2002) Acute oxaliplatin-induced peripheral nerve hyperexcitability. J Clin Oncol 20, 1767–1774.

20. Webster RG, Brain KL, Wilson RH et al. (2005) Oxaliplatin induces hyperexcitability at motor and autonomic neuromuscular junctions through effects on voltage-gated sodium channels. Br J Pharmacol 146, 1027–1039.

21. Lehky TJ, Leonard GD, Wilson RH et al. (2004) Oxaliplatin-induced neurotoxicity: acute hyperexcitability and chronic neuropathy. Muscle Nerve 29, 387–392.

22. Anand U, Otto WR, Anand P (2010) Sensitization of capsaicin and icilin responses in oxaliplatin treated adult rat DRG neurons. Mol Pain 6, 82.

23. Chen K, Zhang Z-F, Liao M-F et al. (2015) Blocking PAR2 attenuates oxaliplatin-induced neuropathic pain via TRPV1 and releases of substance P and CGRP in superficial dorsal horn of spinal cord. J Neurol Sci 352, 62–67.

24. Chukyo A, Chiba T, Kambe T et al. (2018) Oxaliplatin-induced changes in expression of transient receptor potential channels in the dorsal root ganglion as a neuropathic mechanism for cold hypersensitivity. Neuropeptides 67, 95–101.

25. Park SB, Krishnan AV, Lin CS-Y et al. (2008) Mechanisms underlying chemotherapy-induced neurotoxicity and the potential for neuroprotective strategies. Curr Med Chem 15, 3081–3094.

26. Nyffeler J, Dolde X, Krebs A et al. (2017) Combination of multiple neural crest migration assays to identify environmental toxicants from a proof-of-concept chemical library. Arch Toxicol 91, 3613–3632.

27. Wing C, Komatsu M, Delaney SM et al. (2017) Application of stem cell derived neuronal cells to evaluate neurotoxic chemotherapy. Stem Cell Res 22, 79–88.

28. Loser D, Hinojosa MG, Blum J et al. (2021) Functional alterations by a subgroup of neonicotinoid pesticides in human dopaminergic neurons. Arch Toxicol 95, 2081–2107.

29. Loser D, Schaefer J, Danker T et al. (2021) Human neuronal signaling and communication assays to assess functional neurotoxicity. Arch Toxicol 95, 229–252.

30. Klima S, Brüll M, Spreng A-S et al. (2021) A human stem cell-derived test system for agents modifying neuronal N-methyl-D-aspartate-type glutamate receptor Ca2+-signalling. Arch Toxicol 95, 1703–1722.

31. Stiegler NV, Krug AK, Matt F et al. (2011) Assessment of chemical-induced impairment of human neurite outgrowth by multiparametric live cell imaging in high-density cultures. Toxicol Sci 121, 73–87.

32. Boisvert EM, Engle SJ, Hallowell SE et al. (2015) The Specification and Maturation of Nociceptive Neurons from Human Embryonic Stem Cells. Sci Rep 5, 16821.

33. Chen G, Gulbranson DR, Hou Z et al. (2011) Chemically defined conditions for human iPSC derivation and culture. Nat Methods 8, 424–429.

34. Krebs A, van Vugt-Lussenburg BMA, Waldmann T et al. (2020) The EU-ToxRisk method documentation, data processing and chemical testing pipeline for the regulatory use of new approach methods. Arch Toxicol 94, 2435–2461.

35. Klose J, Pahl M, Bartmann K et al. (2021) Neurodevelopmental toxicity assessment of flame retardants using a human DNT in vitro testing battery. Cell Biol Toxicol.

36. Dirks WG, Drexler HG (2013) STR DNA typing of human cell lines: detection of intra- and interspecies cross-contamination. Methods Mol Biol 946, 27–38.

37. Dreser N, Madjar K, Holzer A-K et al. (2020) Development of a neural rosette formation assay (RoFA) to identify neurodevelopmental toxicants and to characterize their transcriptome disturbances. Arch Toxicol 94, 151–171.

38. House JS, Grimm FA, Jima DD et al. (2017) A Pipeline for High-Throughput Concentration Response Modeling of Gene Expression for Toxicogenomics. Front Genet 8, 168.

39. Love MI, Huber W, Anders S (2014) Moderated estimation of fold change and dispersion for RNA-seq data with DESeq2. Genome Biol 15, 550.

40. Raudvere U, Kolberg L, Kuzmin I et al. (2019) g:Profiler: a web server for functional enrichment analysis and conversions of gene lists (2019 update). Nucleic Acids Res 47, W191–W198.

41. Karreman C, Klima S, Holzer A-K et al. (2020) CaFFEE: A program for evaluating time courses of Ca2+ dependent signal changes of complex cells loaded with fluorescent indicator dyes. ALTEX 37, 332–336.

42. North RA (2003) The P2X3 subunit: a molecular target in pain therapeutics. Curr Opin Investig Drugs 4, 833–840.

43. Bonnington JK, McNaughton PA (2003) Signalling pathways involved in the sensitisation of mouse nociceptive neurones by nerve growth factor. J Physiol 551, 433–446.

44. Chuang HH, Prescott ED, Kong H et al. (2001) Bradykinin and nerve growth factor release the capsaicin receptor from PtdIns(4,5)P2-mediated inhibition. Nature 411, 957–962.

45. D’Arco M, Giniatullin R, Simonetti M et al. (2007) Neutralization of nerve growth factor induces plasticity of ATP-sensitive P2X3 receptors of nociceptive trigeminal ganglion neurons. J Neurosci 27, 8190–8201.

46. Namer B, Schick M, Kleggetveit IP et al. (2015) Differential sensitization of silent nociceptors to low pH stimulation by prostaglandin E2 in human volunteers. Eur J Pain 19, 159–166.

47. Zhang X, Huang J, McNaughton PA (2005) NGF rapidly increases membrane expression of TRPV1 heat-gated ion channels. EMBO J 24, 4211–4223.

48. Lallemend F, Ernfors P (2012) Molecular interactions underlying the specification of sensory neurons. Trends Neurosci 35, 373–381.

49. Chen C-L, Broom DC, Liu Y et al. (2006) Runx1 determines nociceptive sensory neuron phenotype and is required for thermal and neuropathic pain. Neuron 49, 365–377.

50. Waldmann T, Grinberg M, König A et al. (2017) Stem Cell Transcriptome Responses and Corresponding Biomarkers That Indicate the Transition from Adaptive Responses to Cytotoxicity. Chem Res Toxicol 30, 905–922.

51. Waldmann T, Rempel E, Balmer NV et al. (2014) Design principles of concentration-dependent transcriptome deviations in drug-exposed differentiating stem cells. Chem Res Toxicol 27, 408–420.

52. Yu Y-Q, Chen X-F, Yang Y et al. (2014) Electrophysiological identification of tonic and phasic neurons in sensory dorsal root ganglion and their distinct implications in inflammatory pain. Physiol Res 63, 793–799.

53. Tsien RW, Tsien RY (1990) Calcium channels, stores, and oscillations. Annu Rev Cell Biol 6, 715–760.

54. Bianchi BR, Lynch KJ, Touma E et al. (1999) Pharmacological characterization of recombinant human and rat P2X receptor subtypes. Eur J Pharmacol 376, 127–138.

55. Koshimizu TA, van Goor F, Tomić M et al. (2000) Characterization of calcium signaling by purinergic receptor-channels expressed in excitable cells. Mol Pharmacol 58, 936–945.

56. North RA (2002) Molecular physiology of P2X receptors. Physiol Rev 82, 1013–1067.

57. Ursu D, Knopp K, Beattie RE et al. (2010) Pungency of TRPV1 agonists is directly correlated with kinetics of receptor activation and lipophilicity. Eur J Pharmacol 641, 114–122.

58. Starkus J, Jansen C, Shimoda LMN et al. (2019) Diverse TRPV1 responses to cannabinoids. Channels (Austin) 13, 172–191.

59. Ambrosino P, Soldovieri MV, Russo C et al. (2013) Activation and desensitization of TRPV1 channels in sensory neurons by the PPARα agonist palmitoylethanolamide. Br J Pharmacol 168, 1430–1444.

60. Jancsó N, Jancsó-Gábor A, Szolcsányi J (1967) Direct evidence for neurogenic inflammation and its prevention by denervation and by pretreatment with capsaicin. Br J Pharmacol Chemother 31, 138–151.

61. Jarvis MF (2010) The neural-glial purinergic receptor ensemble in chronic pain states. Trends Neurosci 33, 48–57.

62. Jarvis MF, Burgard EC, McGaraughty S et al. (2002) A-317491, a novel potent and selective non-nucleotide antagonist of P2X3 and P2X2/3 receptors, reduces chronic inflammatory and neuropathic pain in the rat. Proc Natl Acad Sci U S A 99, 17179–17184.

63. Gever JR, Soto R, Henningsen RA et al. (2010) AF-353, a novel, potent and orally bioavailable P2X3/P2X2/3 receptor antagonist. Br J Pharmacol 160, 1387–1398.

64. Gunthorpe MJ, Rami HK, Jerman JC et al. (2004) Identification and characterisation of SB-366791, a potent and selective vanilloid receptor (VR1/TRPV1) antagonist. Neuropharmacology 46, 133–149.

65. Varga A, Németh J, Szabó A et al. (2005) Effects of the novel TRPV1 receptor antagonist SB366791 in vitro and in vivo in the rat. Neurosci Lett 385, 137–142.

66. Deuis JR, Zimmermann K, Romanovsky AA et al. (2013) An animal model of oxaliplatin-induced cold allodynia reveals a crucial role for Nav1.6 in peripheral pain pathways. Pain 154, 1749–1757.

67. Kanzawa-Lee GA, Knoerl R, Donohoe C et al. (2019) Mechanisms, Predictors, and Challenges in Assessing and Managing Painful Chemotherapy-Induced Peripheral Neuropathy. Semin Oncol Nurs 35, 253–260.

68. Brüning T, Bartsch R, Bolt HM et al. (2014) Sensory irritation as a basis for setting occupational exposure limits. Arch Toxicol 88, 1855–1879.

69. St Germain DC, O’Mara AM, Robinson JL et al. (2020) Chemotherapy-induced peripheral neuropathy: Identifying the research gaps and associated changes to clinical trial design. Cancer 126, 4602–4613.

70. Wheeler HE, Wing C, Delaney SM et al. (2015) Modeling chemotherapeutic neurotoxicity with human induced pluripotent stem cell-derived neuronal cells. PLoS One 10, e0118020.

71. Morrison G, Liu C, Wing C et al. (2016) Evaluation of inter-batch differences in stem-cell derived neurons. Stem Cell Res 16, 140–148.

72. Schinke C, Fernandez Vallone V, Ivanov A et al. (2021) Modeling chemotherapy induced neurotoxicity with human induced pluripotent stem cell (iPSC) -derived sensory neurons. Neurobiol Dis 155, 105391.

73. Klima S, Suciu I, Hoelting L et al. (2021) Examination of microcystin neurotoxicity using central and peripheral human neurons. ALTEX 38, 73–81.

74. Saito-Diaz K, Street JR, Ulrichs H et al. (2021) Derivation of Peripheral Nociceptive, Mechanoreceptive, and Proprioceptive Sensory Neurons from the same Culture of Human Pluripotent Stem Cells. Stem Cell Reports 16, 446–457.

75. García-León JA, Kumar M, Boon R et al. (2018) SOX10 Single Transcription Factor-Based Fast and Efficient Generation of Oligodendrocytes from Human Pluripotent Stem Cells. Stem Cell Reports 10, 655–672.

76. Hulme AJ, McArthur JR, Maksour S et al. (2020) Molecular and Functional Characterization of Neurogenin-2 Induced Human Sensory Neurons. Front Cell Neurosci 14, 600895.

77. Desiderio S, Vermeiren S, van Campenhout C et al. (2019) Prdm12 Directs Nociceptive Sensory Neuron Development by Regulating the Expression of the NGF Receptor TrkA. Cell Rep 26, 3522–3536.e5.

78. Nickolls AR, Lee MM, Espinoza DF et al. (2020) Transcriptional Programming of Human Mechanosensory Neuron Subtypes from Pluripotent Stem Cells. Cell Rep 30, 932–946.e7.

79. Isensee J, Schild C, Schwede F et al. (2017) Crosstalk from cAMP to ERK1/2 emerges during postnatal maturation of nociceptive neurons and is maintained during aging. J Cell Sci 130, 2134– 2146.

80. Rostock C, Schrenk-Siemens K, Pohle J et al. (2018) Human vs. Mouse Nociceptors - Similarities and Differences. Neuroscience 387, 13–27.

81. Wang X-M, Lehky TJ, Brell JM et al. (2012) Discovering cytokines as targets for chemotherapy-induced painful peripheral neuropathy. Cytokine 59, 3–9.

82. Carozzi VA, Canta A, Chiorazzi A (2015) Chemotherapy-induced peripheral neuropathy: What do we know about mechanisms? Neurosci Lett 596, 90–107.

83. Kraus D, Boyle V, Leibig N et al. (2015) The Neuro-spheroid--A novel 3D in vitro model for peripheral nerve regeneration. J Neurosci Methods 246, 97–105.

