## Supplementary File 1 for "Generation of human nociceptor-enriched sensory neurons for the study of pain-related dysfunctions"

**Table of Contents**

| | | |
|---|---|---|
| | Page2 (P2) | Supplementary Methods |
| Tab. S1 | P8 | Primary antibodies used in this study |
| Tab. S2 | P9 | Primers used in this study |
| Tab. S3 | P10 | Agonists and antagonists used in $Ca^{2+}$-imaging experiment |
| Fig. S1 | P11 | Characterization of iPSC-derived sensory neurons |
| Fig. S2 | P12 | Generation of the Sigma-NGN1 iPSC line with inducible expression of NGN1 |
| Fig. S3 | P14 | Short tandem repeat (STR) typing |
| Fig. S4 | P15 | Characterization of neurons resulting from different doxycycline exposure scenarios integrated in the standard differentiation protocol |
| Fig. S5 | P17 | Unbiased, transcriptomics-based approach to explore the differentiation of PNN |
| Fig. S6 | P19 | Driver genes of the oGO terms |
| Fig. S7 | P20 | Expression of functional voltage gated ion channels |
| Fig. S8 | P22 | Threshold setting approach for $Ca^{2+}$ imaging evaluation to define reactive cells |
| Fig. S9 | P24 | Characterization of PNN and their use in $Ca^{2+}$ imaging experiments |
| Fig. S10 | P26 | Effects of platinum compounds on viability parameters |
| Fig. S11 | P28 | Functional changes in mature PNN exposed to platinum compounds |
| | P30 | Supplementary references |
| | P33 | Appendix |

**Supplementary Methods**

***Maintenance of induced pluripotent stem cells (iPSCs)***

Maintenance of the iPSC lines Sigma iPSC0028 (Merck, Darmstadt, Germany) and the in-house gene-edited iPSC line Sigma-NGN1 was performed on human Laminin-521 (BioLamina, Sundbyger, Sweden) coating in essential 8 (E8) medium (Dulbecco's modified Eagle's medium/F12 [DMEM/F12] supplemented with 15 mM Hepes [Thermo Fisher Scientific, Waltham, MA, USA], 10 µg/ml holo-transferrin, 20 µg/ml insulin, 16 mg/ml L-ascorbic-acid, 0.7 mg/ml sodium selenite [all from Merck], 100 ng/ml bFGF [Thermo Fisher Scientific], 1.74 ng/ml TGFb [Bio-Techne, Minneapolis, MN, USA]) essentially as described [1]. Passaging of the iPSCs was performed every 7 days. Cells were incubated with EDTA for 2 min (37°C, 5% $CO_2$) to detach the cells, so that clumps remain (no single cell suspension). iPSCs were washed off the plate with DMEM/F12. Cells were re-seeded in E8 medium on freshly coated plates in a final dilution of 1:40-60.

***Differentiation of sensory neurons from iPSC***

The iPSC line chosen has often been used for the generation of various cell types [2–9]. The iPSCs were prepared for neural differentiation on day of differentiation minus 2 (DoD-2) by replating in a single cell suspension (90,000 cells/cm$^2$) onto Matrigel$^{TM}$ (Corning, Glendale, AZ, USA) coated 6-well plates in E8 medium supplemented with 10 µM Rock inhibitor (Y-27632 [Bio-Techne]).
On DoD0', E8 was replaced by neural differentiation medium KSR (knock out DMEM with 15% serum replacement, 1 x Glutamax, 1 x nonessential amino acids, and 50 µM β-mercaptoethanol [all from Thermo Fisher Scientific]) and the combination of five small molecule pathway inhibitors. From DoD0'-5', 17.5 ng/ml Noggin (Bio-Techne) and 10 µM SB-431642 (Bio-Techne) were added, and 1.5 µM CHIR99021 (Axon Medchem, Groningen, Netherlands), 5 µM SU5402 (Bio-Techne) and 5 µM DAPT (γ-Secretase inhibitor IX) (Merck) were added on DoD2'-9'. From DoD4' onwards, KSR medium was gradually replaced by N2-S medium (DMEM/F12, 1 x GlutaMax [both from Thermo Fisher Scientific], 0.1 mg/ml apotransferrin, 1.55 mg/ml glucose, 25 µg/ml insulin, 20 nM progesterone, 100 µM putrescine and 30 nM selenium [all from Merck]) in 25% increments. On DoD9' the cells were cryopreserved in 90% fetal bovine serum (FBS) (Thermo Fisher Scientific) and 10% dimethyl sulfoxide (DMSO) (Merck).
After thawing of the pre-differentiated cells, sensory neuron precursors were cultured in 25% KSR and 75% N2-S supplemented with CHIR99021 (1.5 µM), SU5402 (5 µM) and DAPT (5 µM). Cells were seeded at a density of 100.000 cells/cm² on Matrigel$^{TM}$ coated plates. For further differentiation and maturation, half of the medium was changed on DoD1 and DoD2. With the fresh culture medium on DoD2, Matrigel was added to the cells at a final dilution of 1:80. On DoD3, medium was changed to N2-S medium supplemented with 12.5 ng/ml brain-derived neurotrophic factor, 25 ng/ml glia-derived neurotrophic factor and 25 ng/ml nerve growth factor (all from Bio-Techne) and 2 µM cytarabin (AraC; Merck). Half of the medium was changed on DoD4, 7 and 10 with further Matrigel addition on DoD10. On DoD14, medium was changed to maturation medium (N2-S supplemented with BDNF [12.5 ng/ml], GDNF and

NGF [both 25 ng/ml]). Half medium exchanges are performed every three to four days. Matrigel is diluted in the culture medium at a final dilution of 1:80 every 10 days.

##### *PeriTox-test*

Immature peripheral neurons were thawed and seeded at a density of 100,000 cells /cm$^2$ [10]. Cells were left to attach for 1 h at 37°C, 5% $CO_2$ followed by treatment with the respective test compounds. Cells were exposed to the compounds for 24 h. One hour prior to analysis, peripheral neurons were stained with 1 µg/ml HOECHST-33342 (H-33342) and 1 µM calcein-AM (both from Merck). After incubation for 1 h at 37°C, 5% $CO_2$, image acquisition was performed automatically using an ArrayScan VTI HCS microscope (Thermo Fisher Scientific). Image analysis was performed as described previously [11]. In brief, the calcein stain was used to identify the neuronal area and the somatic area, defined by the enlarged area of H-33342 stain, was subtracted resulting in the neurite area. The same images were used to derive data on cell viability. Each H-33342 stained cell was checked for a double stain with calcein-AM. Double-positive cells were classified as viable, cells that were only H-33342 positive as dead.

##### *Lentiviral construct and the generation of a gene-edited iPSC line*

The lentiviral sequence was designed to yield a fusion protein consisting of turboRFP (tRFP), the 2A sequence, ubiqitin (Ubi) and NGN1. This construct enables the exact generation of the NGN1 protein by cleavage. Equimolar amounts of tRFP and NGN1 are produced. The expression is driven by a synthetic promoter (Tet-responsive element [TRE]) that is dependant on the presence of Doxycycline (Dox). Furthermore, the lentivirus carries a hygromycin restistance gene allowing the selection for cells that incorporated the lentiviral DNA upon infection (Fig. 2A). The vector and the principle of the fusion construct was published earlier [12].

The human Sigma iPSC0028 line was infected with the described lentivirus. Infected cells underwent hygromycin (Carl Roth, Karsruhe, Germany) selection. After selection, the cells were cultured for 5 days in E8 medium without hygromycin. This was followed by manual picking and expansion of the colonies. Stocks of the clones were cryopreserved in 90% FBS and 10% DMSO. Short tandem repeat (STR) DNA typing (described in detail in [36]) was performed for cell line authentication. Furthermore, bordering sequences of the inserted NGN1 expression virus were isolated by nested inverse PCR. Fragments were purified by gel isolation and sequenced. The construct was found to have integrated in an intron of the glutamate ionotropic receptor kainate type subunit 5 gene (GRIK5) on chromosome 19.

To evaluate the clone's NGN1 expression properties, iPSCs were seeded as single cells in E8 medium supplemented with 10 µM ROCKi, at a density of 10.000 cells/cm$^2$. After one day, medium was exchanged to E8 without ROCKi and cells were exposed to doxycycline (2 µg/ml) for up to 5 days (Fig. 2B).

##### *Immunofluorescence staining and microscopy*

Neurons, grown on glass coverslips coated with Matrigel$^{TM}$, were fixed with 4% paraformaldehyde at 4°C over night. All further steps were performed at room temperature. Paraformaldehyde was taken off and cells were washed (~1 min) with phosphate buffered saline (PBS) followed by permeabilization with 0.6% Triton X-100 in PBS for 7 min. Coverslips were washed (~1 min) with PBS and blocked for 1 h in PBS containing 5% FCS and 0.1% Triton X-100. Primary antibodies (see Tab. S1) were diluted in fresh blocking solution and applied for 1 h. Residual free primary antibodies are then washed off with PBS. Secondary antibodies and H-33342 are diluted in blocking solution and applied on the coverslips for 30 min. After washing with PBS, coverslips were placed upside-down on mounting medium on microscope slides.

***RNA extraction, cDNA synthesis and reverse transcriptase qPCR***

RNA was extracted with TRIzol (Thermo Fisher Scientific) according to the manufacturer's protocol. To produce cDNA, reverse transcription was performed with 1 µg RNA using iScript (Bio-Rad, Hercules CA, USA), following the manufacturer's protocol. SsoFast™ EvaGreen® Supermix (Bio-Rad) was used to quantify the cDNA. Determination of the threshold cycle ($C_T$) was done with the CFX data analysis software (Bio-Rad). Reference genes were used for normalization of the mRNA levels of the genes of interest which were then compared for different time points of differentiation according to the ΔΔ method [13]. Primers used in this study are listed in detail in table S2.

***Transcriptome data generation and analysis***

Sample lysates were prepared by medium removal, followed by a wash with 50 µl of phosphate buffered saline (PBS) (Thermo Fisher Scientific) and instant addition of 33 µl 1x Biospyder lysis buffer (BioSpyder Tech., Glasgow, UK). After incubation at RT for 10 minutes, the sample plates were stored at -80°C up to the time of dry ice shipping to Bioclavis (BioSpyder Tech., Glasgow, UK). The whole transcriptome was then measured via the TempO-Seq targeted sequencing technology [14]. The set of genes analysed, and the read data are detailed in Supplement file2, organized as Excel workbook. Labelling and clear explanations are included.

Downstream data interpretation was performed using the R package DESeq2 (v1.32.0) for the differential gene expression (DGE) analysis [15]. The raw probe counts were normalized to total sample counts per million (CPM). No library size threshold was used; samples with replicate correlation (Pearson R) to group average below 0.8 were removed from the analysis. Prior to the DGE, the low-count genes (less than 3 samples above 5 CPM) were discarded. The Wald statistics test was used for significance evaluation of each differentiation stage against DoD1. Selection of the most significant differentially expressed genes (DEG) was done using a (Benjamini-Hochberg) p-adjusted maximum threshold of 0.05 and a $\log_2$(fold change) minimum threshold of 1. Gene ontology (GO) terms were analysed for over-representation using g:profiler software [16], based on Fisher's *F* test. GO terms of the category biological process with a maximum size of 1000 genes were taken into account for further analysis. Quantitative activation scores were calculated according to Waldmann et al. (2014) by by "multiplying the percentage of genes within the GO that was found to be significantly regulated with the average fold change of these regulations" [17].

***Manual patch-clamp recordings***

Patch-clamp experiments were performed in the whole-cell mode [18] using an EPC 10 USB patch-clamp amplifier and the PATCHMASTER software (version 2x91; HEKA Elektronik, Lambrecht, Germany). The extracellular solution contained (in mM): 140 NaCl, 4 KCl, 1

$MgCl_2$, 1.8 $CaCl_2$, 10 HEPES, and 10 D-glucose, pH 7.4. The intracellular solution contained (in mM): 107 K-gluconate, 10 KCl, 1 $MgCl_2$, 10 HEPES, 5 EGTA, and 4 $Na_2ATP$, pH 7.2. For the experiments in which the voltage-gated potassium ion channel currents were blocked, the following solutions were used: the extracellular solution contained (in mM): 140 NaCl, 10 tetraethylammonium chloride (TEA), 1 $MgCl_2$, 1.8 $CaCl_2$, 10 HEPES, and 10 D-glucose, pH 7.4. The intracellular solution contained (in mM): 117 CsCl, 10 TEA, 1 $MgCl_2$, 10 HEPES, 5 EGTA, and 4 $Na_2ATP$, pH 7.2. All substances were obtained from Carl Roth except K-gluconate, EGTA, $Na_2ATP$, TEA, and CsCl, which were from Sigma Aldrich.
The recordings were performed at room temperature and the cells were kept at a holding potential of -70 mV in all experiments. To investigate the action potential firing behavior, the cells were stimulated in current-clamp mode by hyper- and depolarizing current pulses from -50 pA to +240 pA in +10 pA steps with a pulse duration of 300 ms. In voltage-clamp mode, voltage-gated ion channels were stimulated by voltage pulses ranging from -70 mV to +60 mV in +10 mV steps with a pulse duration of 300 ms. For leak subtraction, the P/4 algorithm of the PATCHMASTER software was used in voltage-clamp mode. Both pulse protocols were executed at 0.2 Hz. For agonist tests with α,β-methylene ATP (α,β-meATP) and capsaicin in voltage-clamp mode, cells were exposed to these compounds for 5 s. Tetrodotoxin (TTX), α,β-meATP, and capsaicin were obtained from Tocris.
The data of the manual patch-clamp recordings were analyzed and visualized with scripts written in R (version 3.6.3) [19]. The following r packages were utilized for data handling: cowplot [20], ephys2 [21], ggplot2 [22], lemon [23].

***Microelectrode array recordings***

Sensory neurons were seeded at a density of 120.000 cells/7 µl drop on Matrigel™ coated 24-well CytoView multielectrode array (MEA) plates with 16 electrodes per well (Axion Biosystems, Atlanta, USA). Neurons were left at 37°C, 5% $CO_2$ for 1 h to attach before the wells were filled with 500 µl DoD0 medium. Neurons were further differentiated and matured as described above. Measurement was performed on different days of differentiation for the same wells of a plate. After an equilibration time of 15 min, recordings with a Maestro Edge (Axion Biosystems) were started. Recordings were carried out for 15 min every hour for a total of 24 h to 38 h. In case of oxaliplatin treatment, the compound was applied in 50 µl of maturation medium. Control neurons were treated with 50 µl maturation medium alone. All recordings were captured using the Axion Integrated Studio Navigator (Axion Biosystems) with a recording chamber at 37°C and 5% $CO_2$. For raw data acquisition, signals from all electrodes were recorded simultaneously with a sampling frequency of 12.5 kHz/channel. The recorded raw files were converted offline from voltage traces into various time-dependent data sets, such as spiking frequency, etc. The threshold spike detector was set to 5.5x of the noise level (signal SD) on each electrode, using adaptive threshold crossing for spike detection [3].

***Measurement of changes in intracellular $Ca^{2+}$-concentration $[Ca^{2+}]_i$***

Immature sensory neurons were seeded on DoD0 in 96-well plates at a density of 100.000 cells/cm$^2$. They were differentiated as described above for up to 49 days. One hour before the measurement, neurons were loaded with Fluo-4 Direct™ Calcium Assay Kit (Thermo Fisher Scientific) and H-33342 by exchanging half of the medium with the staining solution. Preincubation with e.g. receptor antagonists was done together with Fluo-4 loading. Test compounds were diluted in Hanks' Balanced Salt Solution (HBSS). Agonists and antagonists used in this study are listed in detail in table S3.

Monitoring of $[Ca^{2+}]_i$ was performed using a VTI HCS microscope (Thermo Fisher Scientific) equipped with an automated pipettor and an incubation chamber. Neurons were kept at 37°C and 5% $CO_2$ during the experiments. Images were taken as fast as possible for 45 s and test compounds were applied automatically 10 s after the first picture was taken. In a standard experiment with 4 stimuli applied to one well (e.g. negative control, P2X3 agonist, TRPV1 agonist, KCl), the cells were imaged 4 times for 45 s with one stimulus applied at a time. Subsequent stimuli applied to one well were separated by a minimum interval of 5 minutes.

The images were exported as .avi video files and analysed with the CaFFEE software [24]. In brief, the time point of peak fluorescence was identified. Fluorescence data for the ground state ($F_0$) and for the peak time point ($F_1$) were assessed automatically for all cells. The difference between the two fluorescence levels, $\Delta F=F_1-F_0$, was used for further data processing [3].

##### *Establishing a thresholding strategy to define reactive cells in $Ca^{2+}$-imaging experiments*

$Ca^{2+}$ imaging data from 10 different experiments were used to explore the performance and practical application of four different threshold setting approaches. The objective was to identify the best way of defining reactive cells (Fig. S8A). Data were collected on the response to a negative control and a positive control (KCl). Only the data on Hanks' Balanced Salt Solution (HBSS, negative control) were used for thresholding. Two different methods were used:

Method **1** used a noise level-based cut off: The threshold (T) was defined at T=mean + (3 x SD). The mean fluorescence change triggered by HBSS (mean(ΔHBSS)) was considered as the noise level and 3 x SD(ΔHBSS) as the noise bandwidth. Three different approaches to determine a noise-level based cut off were tested. Method **1a** defined one general threshold that should be applicape to all future experiments. For that purpose, >9000 HBSS Δ values of the whole test set (80 PNN cultures from 10 biological replicates) were combined. In this case, a threshold of T=18 was found. Method **1b** calculated a *well-specific* threshold: to each well (technical replicate), HBSS was applied in a first step and the resulting Δ values were used to define a specific threshold for further stimulus applications to this same well. Method **1c** resulted in an *experiment-specific* threshold with additional normalization by the KCl response (which may differ in absolute height between experiments): for each biological replicate the mean of "base" values of all HBSS measurements was calculated and substracted from the mean of all "top" values of all KCl measurements. The resulting number was defined as $\Delta_{max}$ and set to 100% change in signal intensity. All Δ values of this experiment were normalized to $\Delta_{max}$. The normalized data for HBSS was then further used for threshold calculation. All normalized HBSS Δ values of one experiment (8 technical replicates) were combined and the resulting threshold was then used for all measurements of this experiment.

Method **2** used a population-based cut off: In this approach T was defined at the 95% percentile of the ΔHBSS distribution. All HBSS Δ values of the test set were combined and the highest 5% of these Δ values were defined as "reactive". We defined the next lower integer Δ value as threshold (T) for reactivity. A threshold of T=13 resulted from this population-based cut off.

Reactive cells in response to HBSS, α,β-methylene ATP (1 µM) and capsaicin (1 µM) were quantified according to all 4 threshold setting approaches. Based on the comparison of the respective results, we chose, the well-specific method **1b** as standard thresholding approach for all further experiments. Additionally, we applied the rule that T=18 (method **1a**) was the maximum value acceptable for T.

***Software used to transform images into accessible data***

Most of the data was generated using the published program CaFFEE [24]. For the evaluation of the reactions of individual cells on different stimuli a special version of this program was developed. This special version, called MultiMovie, is capable of aligning the cells in up to five movies and follow their individual behaviour through all successive treatments (Fig. S9D). This enabled us to assess cells that reacted only to some or to certain combinations of stimuli and capture these quantatively.

The accumulated amount of data of these programs exceeded the capacity of all our regularly used programs to visualize data. To accomplish this task, another program (BigData) was written, see figure 5D and S8A,B.

***Supplementary table 1: Primary antibodies used in this study***

| **Target** | **Isotype** | **Dilution** | **Supplier** | **Catalogue number** |
|---|---|---|---|---|
| βIII-tubulin (pol.) | mouse IgG1 | 1:1000 | BioLegend | 921001 |
| Peripherin | mouse IgG2a | 1:200 | Santa Cruz | sc-377093 |
| Ki67 (PE) | mouse IgG1 | 1:500 | BD Pharmingen | 556027 |
| ISL1 | rabbit | 1:200 | Abcam | ab109517 |
| BRN3A | rabbit | 1:200 | Merck Millipore | 5945 |
| P2X3 | rabbit | 1:200 | Novus | NB100-1654 |
| TRPV1 | rabbit | 1:200 | Novus | NBP1-71774 |
| NGN1 | rabbit | 1:100 | Invitrogen | MA5-24912 |
| Nanog | mouse IgG1 | 1:200 | CellSignaling | 9656 (Pluripotency Kit) |
| OCT4 | rabbit | 1:200 | | |
| SOX2 | mouse IgG1 | 1:200 | | |
| Tra1-81 | mouse IgM | 1:500 | | |
| Tra1-60 | mouse IgM | 1:200 | | |
| SSEA4 | mouse IgG3 | 1:500 | | |
| PAX6 | rabbit | 1:200 | BioLegend | 901308 |
| SOX10 | mouse IgG1 | 1:200 | Abcam | ab181466 |

***Supplementary table 2: Primers used in this study***

| **Target** | **Sequence (For)** | **Sequence (Rev)** |
|---|---|---|
| POU4F1 (BRN3A) | CCCTGAGCACAAGTACCCGTC | CGGCTTGAAAGGATGGCTCTTGC |
| GAPDH | ATGGAGAAGGCTGGGGCTCA | AGTGATGGCATGGACTGTGGTCAT |
| ISL1 | TTGGAATGGCATGCGGCATG | AGGCCACACAGCGGAAACAC |
| NANOG | GGTGAAGACCTGGTTCCAGAAC | CATCCCTGGTGGTAGGAAGAGTAAAG |
| NGN1 | GACGACACCAAGCTCACCAA | AACAAGCGGCTCAGGTATCC |
| NGN2 | AGGCCAAAGTCACAGCAACG | GGCTCCTCCTCCTCTTCTTC |
| OCT4 | GCAAAGCAGAAACCCTCGTGC | ACACTCGGACCACATCCTTCTCG |
| P2RX3 | TGACGCCACCTCAGGGCACC | AGGTCCGGAGCACAGAGCTG |
| PAX6 | CCGCCTATGCCCAGCTTCAC | AAGTGGTGCCCGAGGTGCCC |
| PRPH | GAGAGCTGGAGCTGTTGGGC | AGGCGGGACAGAGTGGCATC |
| RET | GGTCTTTTGGTGTCCTGCTGTGG | GCATCAGGCGGTACATCTCCTC |
| RPL13A | GGTATGCTGCCCCACAAAACC | CTGTCACTGCCTGGTACTTCCA |
| RUNX1 | ACCTCGAAGACATCGGCAGAAACTAG | GGAGTGGTTCAGGGAGGCAC |
| RUNX3 | AGACCCCAATCCAAGGCACC | CCACGCTGAGGCTGCTGAT |
| SCN10A | GTGGGCCTGCATGGAAGTTG | GGGCCACCTGCAGGTTGTTC |
| SCN9A | TGCTCTCTGTCTGAGGTTGGG | GCCGGTGAACGGGAAAATGC |
| SOX10 | CAGCAAAAGCAAGCCGCACG | CTTTCGTTCAGCAGCCTCCAGAG |
| TBP | GGGCACCACTCCACTGTATC | GCAGCAAACCGCTTGGGATTATATTCG |
| TRKA | GCTGTCAAGGCACTGAAGGAGG | CGGAGGAAGCGGTTGAGGTC |
| TRKB | ACAGATTTCTGCTCACTTCATGGGC | CCACAGCATAGACCGAGAGATGTTCC |
| TRKC | TCGCTGGATGCCTCCTGAAAG | CAATGACCTCCGTGTTTGAGAGTTGG |
| TRPM8 | TGGGAGCCAGCAAGCTTCTG | CCGCCAGCTCCAGACAGTTG |
| TRPV1 | AGAACGGAGCAGACGTCCAG | GTTGCCCACCGAGTCCCTGG |

***Supplementary table 3: Agonists and antagonists used in $Ca^{2+}$-imaging experiments***

| **Compound** | **Solvent** | **Concentration [µM]** | **Supplier** | **Catalogue number** |
|---|---|---|---|---|
| α,β-methylene ATP | water | 0.1, 0.3, 1, 3, 10 | Cayman Chemical | 10008956 |
| A-317491 | DMSO | 0.1, 1, 2.5, 5, 10 | Cayman Chemical | 19256 |
| AF-353 | DMSO | 0.003, 0.01, 0.03, 0.1, 0.3, 1 | Cayman Chemical | 23034 |
| capsaicin | DMSO | 0.1, 1, 10 | Cayman Chemical | 92350 |
| capsazepine | DMSO | 0.1, 1, 2.5, 5, 10 | Cayman Chemical | 10007518 |
| carbamazepine | DMSO | 25, 100 | Merck | C4024 |
| KCl | water | 40,000 | Merck | P9541 |
| olvanil | DMSO | 25 | Cayman Chemical | 90262 |
| piperine | DMSO | 100 | Merck | P49007 |
| SB-366791 | DMSO | 0.01, 0.1, 1, 10 | Adipogen | AG-CR1-0034 |
| tetrodotoxin | water | 1 | BioTechne | 1078 |
| veratridine | DMSO | 3 | alomone labs | V-110 |

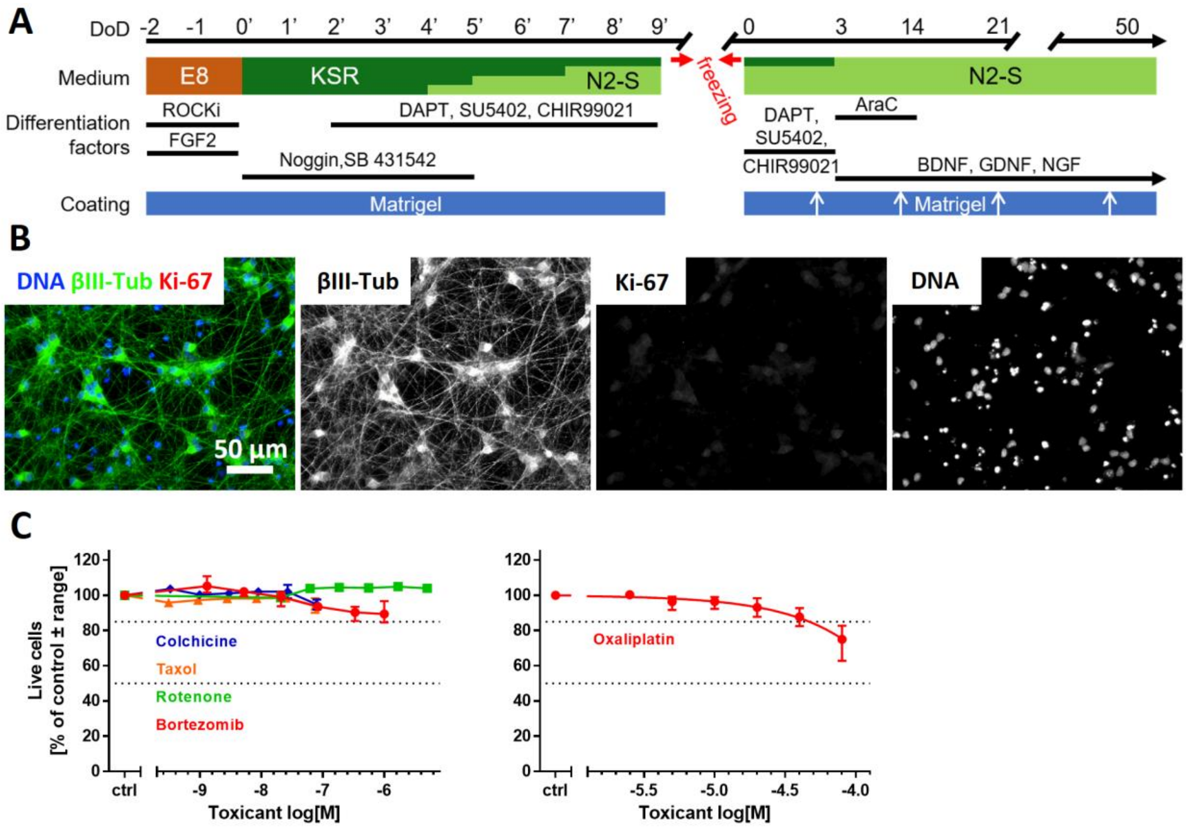

**Supplementary figure 1: Characterization of iPSC-derived sensory neurons**

**(A)** Human sensory neurons were differentiated using the media, coating and supplements indicated. The first differentiation phase is shown left, after switching of E8 stem cell medium to differentiation media containing different ratios of knockout serum replacement (KSR) and N2-S media (with Noggin and the small molecule inhibitors SB431542, DAPT, CHIR99021 and SU5402 added). Note that the timing of this step is indicated by primed numbers. During the second phase (right), starting on DoD0, cells are matured for up to 50 days. The initial medium composition of 25% KSR and 75% N2-S is followed (DoD3) by 100% N2-S medium supplemented with brain-derived neurotrophic factor (BDNF), glia-derived neurotrophic factor (GDNF) and nerve growth factor (NGF). Cells were grown on Matrigel-coated plates, white arrows indicate the addition of liquid Matrigel to the culture medium every 10 days. AraC, cytarabine; FGF2, fibroblast growth factor 2; ROCKi, ROCK inhibitor **(B)** Fixed sensory neurons were immunostained for βIII-tubulin (βIII-Tub) and the proliferation marker Ki-67. The composite image is colour coded and the single stain signals are given as b/w images. **(C)** The PeriTox-test was performed on DoD0 using immature peripheral neurons to assess the effects of toxicant exposure (24 h) on the neurite area and on general viability. Concentration-response curves of the effects on cell survival are shown. Data are given as mean ± range of 2-3 biological replicates. Errorbars smaller than the data point symbols are not shown. Data on neurite area are displayed in figure 1D.

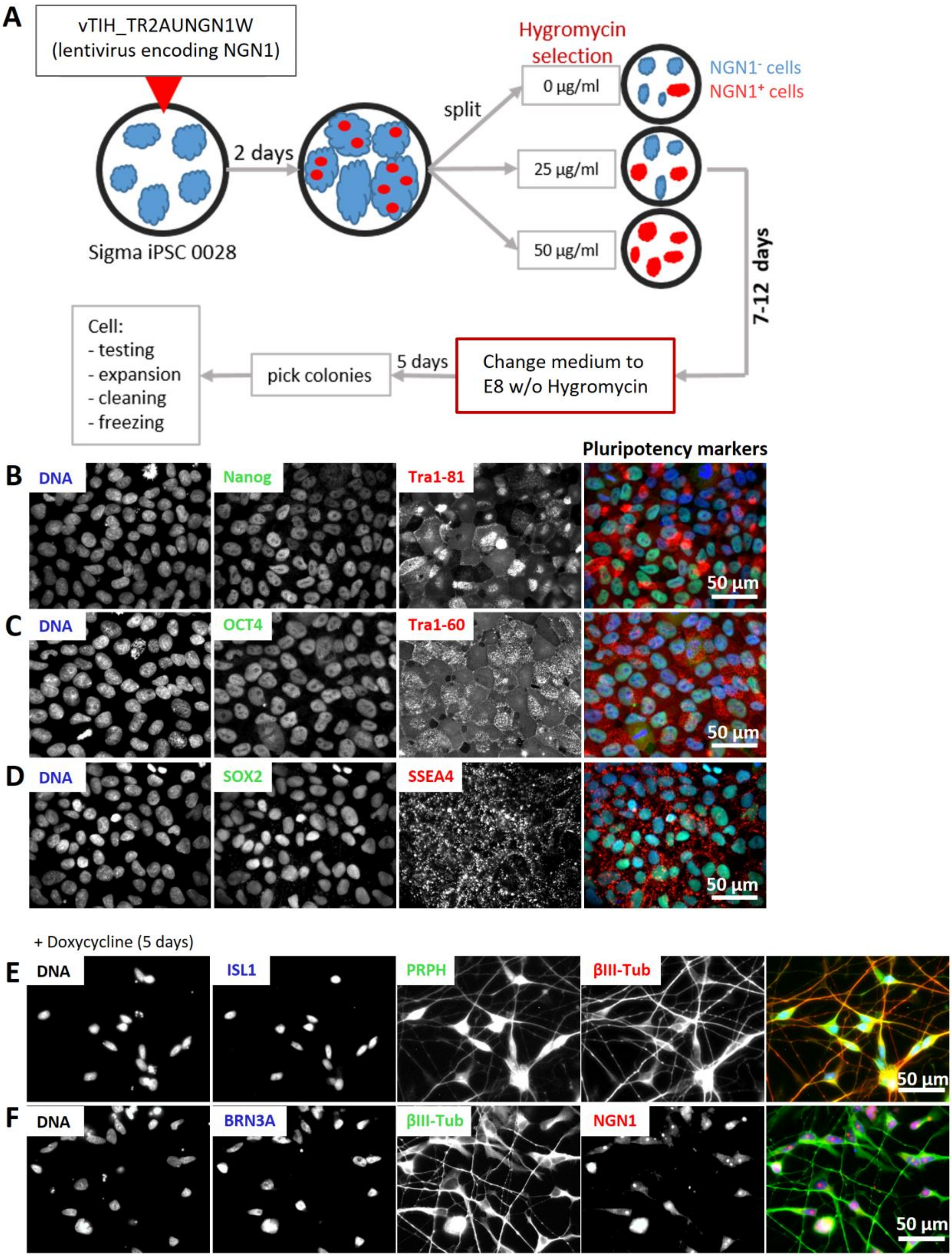
A
vTIH_TR2AUNGN1W
(lentivirus encoding NGN1)
2 days
Sigma iPSC 0028
split
Hygromycin selection
0 µg/ml
25 µg/ml
50 µg/ml
NGN1- cells
NGN1+ cells
7-12 days
Change medium to E8 w/o Hygromycin
5 days
pick colonies
Cell:
- testing
- expansion
- cleaning
- freezing
Pluripotency markers
B
DNA
Nanog
Tra1-81
50 µm
C
DNA
OCT4
Tra1-60
50 µm
D
DNA
SOX2
SSEA4
50 µm
+ Doxycycline (5 days)
E
DNA
ISL1
PRPH
βIII-Tub
50 µm
F
DNA
BRN3A
βIII-Tub
NGN1
50 µm

**Supplementary figure 2: Generation of the Sigma-NGN1 iPSC line with inducible expression of NGN1**

**(A)** Workflow for the generation of the iPSC-line Sigma-NGN1 from the commercially available Sigma iPSC 0028 line. Sigma iPSC 0028 were infected with the lentivirus vTIH_TR2AUNGN1W, encoding NGN1. Cells were grown for two days, split and then cultured under different hygromycin selection conditions (0, 25 or 50 µg/ml) for 7-12 days. Afterwards, the medium was changed to E8 medium without (w/o) hygromycin for 5 days. Single colonies were picked, expanded and cleaned followed by testing (as depicted in Fig. 2B) of the clones. Sigma-NGN1 iPSC stocks were frozen. **(B-D)** Immunofluorescence images of iPSC cultures. Cells were stained for the pluripotency markers Nanog, Tra1-81 **(B)**, OCT4, Tra1-60 **(C)**, SOX2 and SSEA4 **(D)**. Nuclei were stained with H33342 (DNA). **(E,F)** Immunofluorescence images of cells treated with doxycycline for 5 days. Fixed cells were labelled with antibodies against the pan-neuronal marker β-III tubulin (βIII-Tub), the PNS specific intermediate filament peripherin (PRPH) and the sensory neuronal transcription factors NGN1, ISL1 and BRN3A (see also Fig. 2D). **(B-F)** Images of the single stainings are shown. The composite images of the stainings are colour coded according to the detail images. Scale bars are given in the images.

STR analysis of Sigma iPSC0028

| STR loci | Data-sheet (Sigma/ Merck) | | wt (our data) | | NGN1 (our data) | |
|---|---|---|---|---|---|---|
| TH01 | 9 | 9.3 | 9 | 9.3 | 9.3 | 9.3 |
| D5S818 | 12 | 13 | 12 | 13 | 12 | 13 |
| D13S137 | 8 | 12 | 8 | 12 | 8 | 12 |
| D7S820 | 8 | 11 | 8 | 11 | 8 | 11 |
| D16S539 | 11 | 12 | 11 | 12 | 11 | 12 |
| SCF1P0 | 10 | 12 | 10 | 12 | 10 | 12 |
| AMEL | X | X | X | X | X | X |
| vWA | 16 | 16 | 16 | 16 | 16 | 16 |
| TPOX | 8 | 9 | 8 | 9 | 8 | 9 |
| D3 | | | 15 | 16 | 15 | 16 |
| D21 | | | 28 | 30 | 28 | 30 |
| D18 | | | 13 | 16 | 13 | 16 |
| PentaE | | | 7 | 10 | 7 | 10 |
| Penta D | | | 8 | 11 | 8 | 11 |
| D8 | | | 10 | 14 | 10 | 14 |
| FGA | | | 22 | 25 | 22 | 25 |
| D19 | | | 13 | 13 | 13 | 13 |
| D2 | | | 17 | 20 | 17 | 20 |

**Supplementary figure 3: STR typing of the iPSC lines Sigma iPSC 0028 and Sigma-NGN1**

Short tandem repeat (STR) analysis was used to authenticate the newly generated iPSC line Sigma-NGN1. The STR profile for Sigma iPSC 0028 provided by the vendor is given in green columns. A larger set of STR loci was investigated for Sigma iPSC 0028 (as used in our laboratory) and Sigma-NGN1 iPSC. For each STR locus, the number of repeats is given for both alleles (two sub-columns per sample).

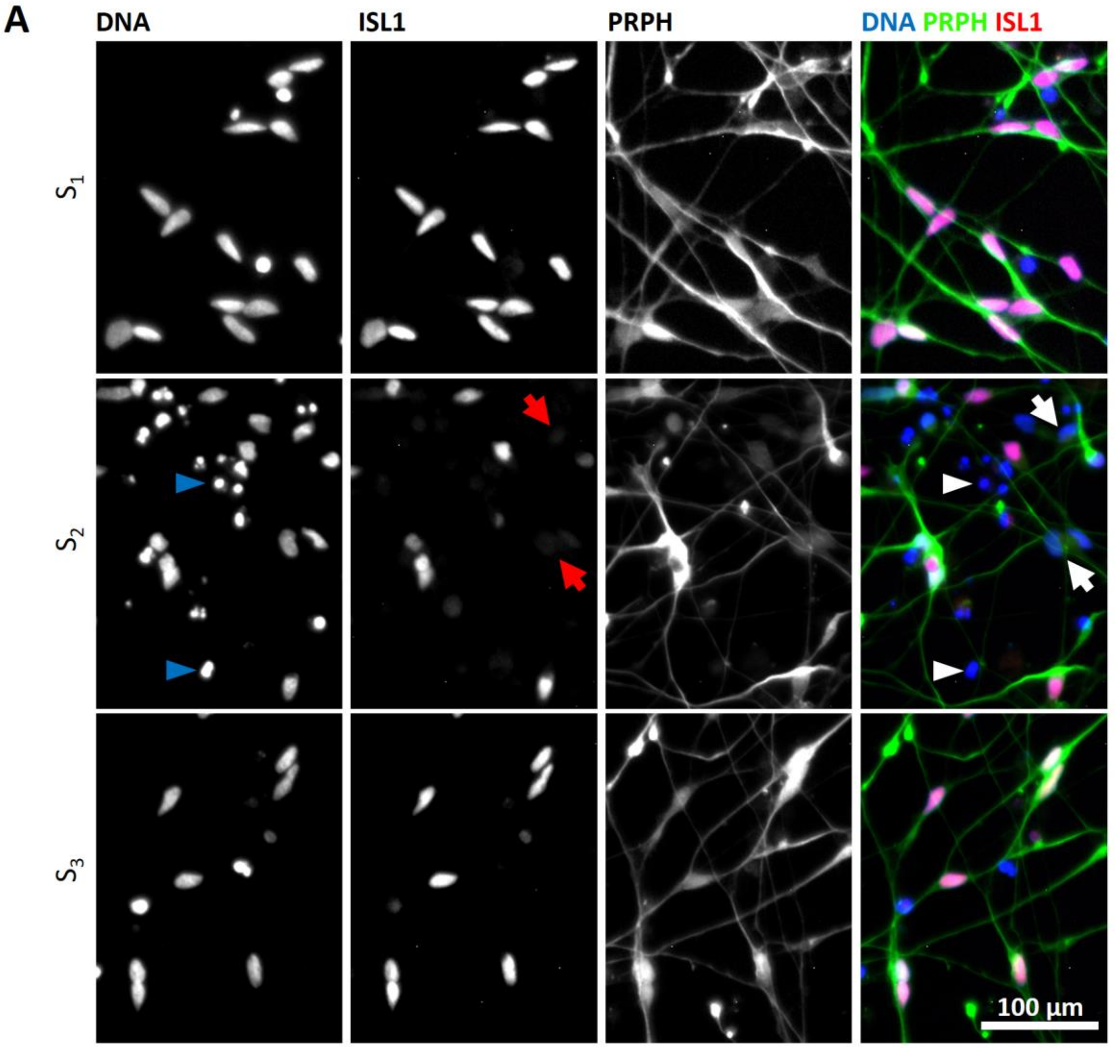

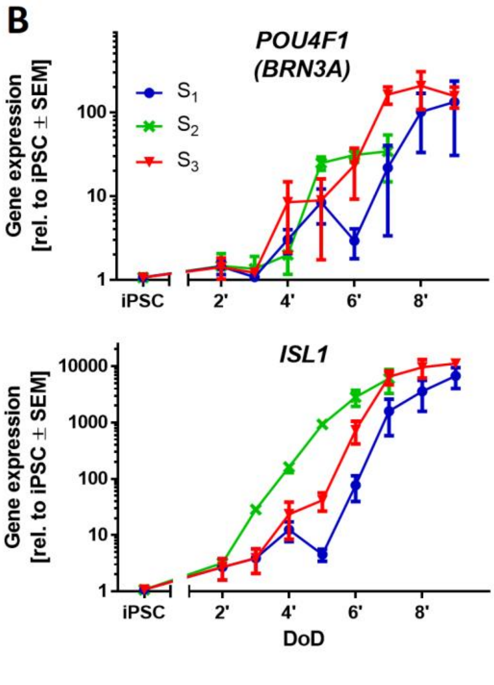

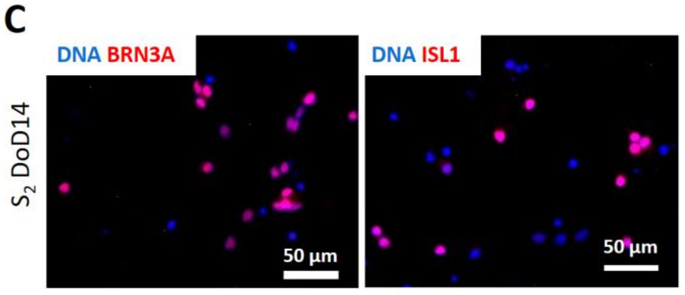

**D**

| % positive nuclei | $S_1$ | $S_2$ | $S_3$ |
|---|---|---|---|
| $BRN3A^+$ | 74-100 | 40-69 | 90-92 |
| $ISL1^+$ | 98-100 | 43-65 | 92-100 |

**Supplementary figure 4: Characterization of neurons resulting from different doxycycline exposure scenarios integrated in the standard differentiation protocol**

**(A)** Immunofluorescence images of cells differentiated according to doxycycline exposure scenarios $S_1$-$S_3$ on DoD3 (after thawing). Fixed cells were labelled with antibodies against PRPH (green) and ISL1 (red). Nuclei were stained with H33342 (blue). Single staining images of the composite images shown in figure 3C are given. The composite images are colour coded and the scale bar is given in the image. Arrowheads depict exemplary dead cells, arrows point out exemplary ISL1-negative nuclei. **(B)** Gene expression analysis of the sensory neuron marker genes *POU4F1* (*BRN3A*) and *ISL1*. Data are expressed relative to expression levels in iPSCs and given as means ± SEM of 3-4 biological replicates. **(C)** Immunofluorescence images of cells differentiated according to $S_2$. Cells were fixed on DoD14 and labelled with antibodies against BRN3A or ISL1 (red). Nuclei were stained with H33342 (blue). Scale bars are given in the images. **(D)** Quantification of ISL1-positive and BRN3A-positive nuclei in DoD14 cultures differentiated according to the three doxycycline exposure conditions described in figure 3A. The range of the percentage of positive nuclei is given.

.

A

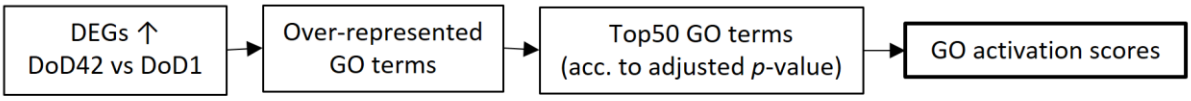

B

| | GO term | DoD 7 | DoD 14 | DoD 28 | DoD 35 | DoD 42 |
|---|---|---|---|---|---|---|
| 1 | synaptic signaling | x | x | x | x | x |
| 2 | chemical synaptic transmission | x | x | x | x | x |
| 3 | anterograde trans-synaptic signaling | x | x | x | x | x |
| 4 | trans-synaptic signaling | x | x | x | x | x |
| 5 | learning or memory | x | x | x | x | x |
| 6 | cognition | x | x | x | x | x |
| 7 | regulation of trans-synaptic signaling | x | x | x | x | x |
| 8 | modulation of chemical synaptic transmission | x | x | x | x | x |
| 9 | neurotransmitter secretion | x | x | x | x | x |
| 10 | signal release from synapse | x | x | x | x | x |
| 11 | neurotransmitter transport | x | x | x | x | x |
| 12 | regulation of cation channel activity | x | x | x | x | x |
| 13 | synaptic transmission, glutamatergic | x | x | x | x | x |
| 14 | regulation of neurotransmitter levels | x | x | x | x | x |
| 15 | synaptic vesicle cycle | x | x | x | x | x |
| 16 | synaptic vesicle exocytosis | x | x | x | x | x |
| 17 | vesicle-mediated transport in synapse | x | x | x | x | x |
| 18 | regulation of synaptic transmission, glutamatergic | x | x | x | x | x |
| 19 | regulation of transmembrane transporter activity | x | x | x | x | x |
| 20 | regulation of signaling receptor activity | x | x | x | x | x |
| 21 | neuron projection morphogenesis | x | x | x | x | x |
| 22 | regulation of ion transmembrane transporter activity | x | x | x | x | x |
| 23 | regulation of vesicle-mediated transport | x | x | x | x | x |
| 24 | plasma membrane bounded cell projection morphogenesis | x | x | x | x | x |
| 25 | cell projection morphogenesis | x | x | x | x | x |
| 26 | regulation of regulated secretory pathway | x | x | x | x | x |
| 27 | neuron cell-cell adhesion | x | x | x | x | x |
| 28 | regulation of intracellular protein transport | | | x | x | x |
| 29 | regulation of transporter activity | x | x | x | x | x |
| 30 | positive regulation of intracellular protein transport | | | x | x | x |
| 31 | dendrite morphogenesis | x | | x | x | x |
| 32 | cellular component morphogenesis | x | x | x | x | x |
| 33 | cell morphogenesis involved in neuron differentiation | x | x | x | x | x |
| 34 | cell part morphogenesis | x | x | x | x | x |
| 35 | regulation of anatomical structure size | x | x | | x | x |
| 36 | postsynapse organization | x | x | x | x | x |
| 37 | regulation of cellular component size | x | x | x | x | x |
| 38 | regulation of synaptic vesicle exocytosis | x | x | x | x | x |
| 39 | synapse organization | x | x | x | x | x |
| 40 | regulation of exocytosis | x | x | x | x | x |
| 41 | positive regulation of synaptic transmission | x | x | x | x | x |
| 42 | regulation of neurotransmitter secretion | x | x | x | x | x |
| 43 | regulation of cation transmembrane transport | x | x | x | x | x |
| 44 | neuron projection organization | x | x | x | x | x |
| 45 | regulation of neurotransmitter receptor activity | x | x | x | x | x |
| 46 | positive regulation of kinase activity | x | x | x | x | x |
| 47 | synapse assembly | x | x | x | x | x |
| 48 | regulation of intracellular transport | x | x | x | x | x |
| 49 | dendritic spine organization | x | x | x | x | x |
| 50 | positive regulation of intracellular transport | | | x | x | x |

C

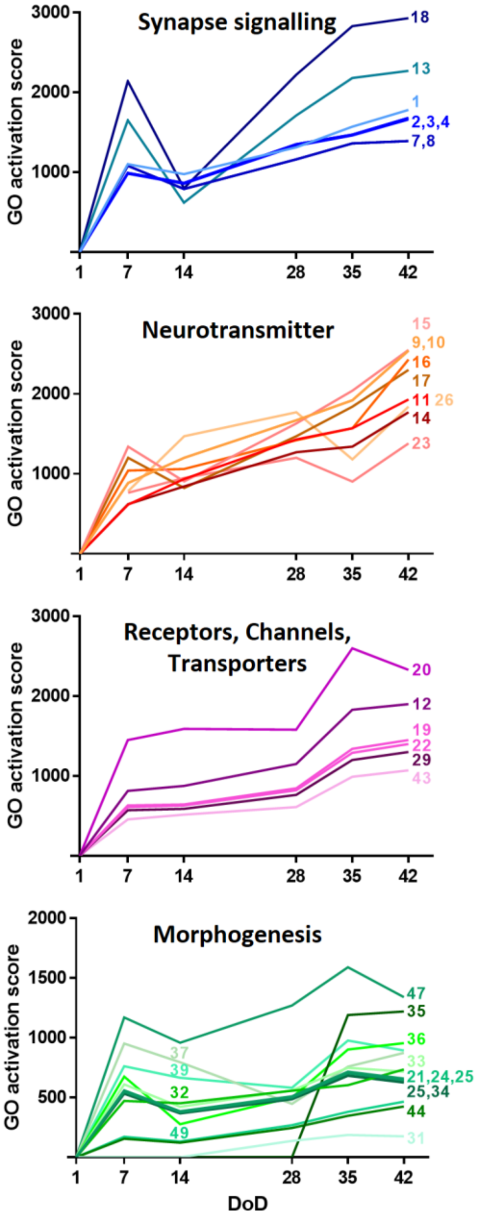

**Supplementary figure 5: Unbiased, transcriptomics-based approach to explore the differentiation of PNN**

Transcriptome data were obtained by the TempO-Seq approach for 19,000 genes and DoD1, 7, 14, 28, 35, 42. **(A)** First, DEGs were determined for DoD42 cells (vs. DoD1). All upregulated DEGS with a fold change > 2 and an adjusted *p*-value ≤ 0.05 were used for over-representation analysis of gene ontology (GO) terms [16]. GO terms of the category biological process with a maximum size of 1000 genes were taken into account for the analysis. The Top50 GO terms with the lowest adjusted *p*-value were used for calculation of quantitative activation scores [17]. **(B)** The Top50 GO terms are listed according to their adjusted *p*-value (1 = lowest *p*-value). The table gives information about the over-representation of the respective GO terms on the investigated days of differentiation (x = over-represented). The GO terms are coloured according to their assigned superordinate group: “synapse signalling” (blue), “neurotransmitter” (orange), “receptors, channels, transporters” (pink) and “morphogenesis” (green). **(C)** Quantitative activation scores for each of the Top50 GO terms are shown over time. Numbers on the right refer to the list numbers **(B)** of the respective GOs depicted in the graphs. Summary data, e.g. combined scores for the four superordinate processes colour-coded here, are displayed in figure 4.

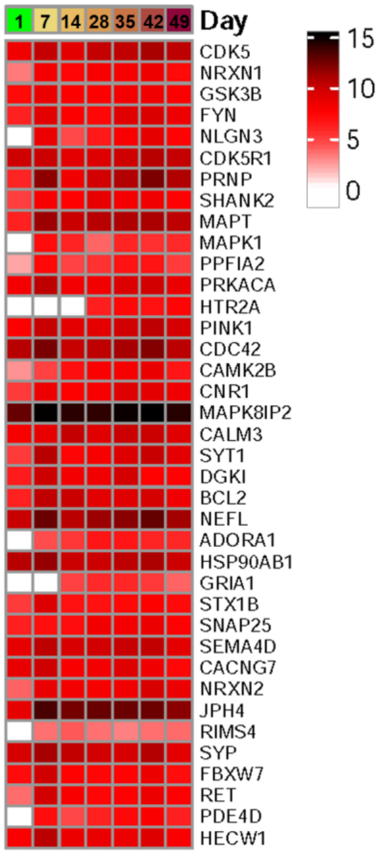

**Supplementary figure 6: Driver genes of the oGO terms**

Among the DEGs determined for DoD42 *versus* DoD1, 200 GO terms were found to be over-represented. We were interested, which genes were drivers of these GOs. The number of GO terms, in which each gene occurs, was quantified. The genes found in more than 15% of the oGO terms were further analyzed for their absolute transcript levels to confirm that the oGOs were driven by reasonably expressed genes. The heatmap shows the absolute expression levels of the GO driver genes for all timepoints examined. Data are given as counts per million (CPM) on a $\log_2$ scale. The complete set of raw data is found in supplementary file2.

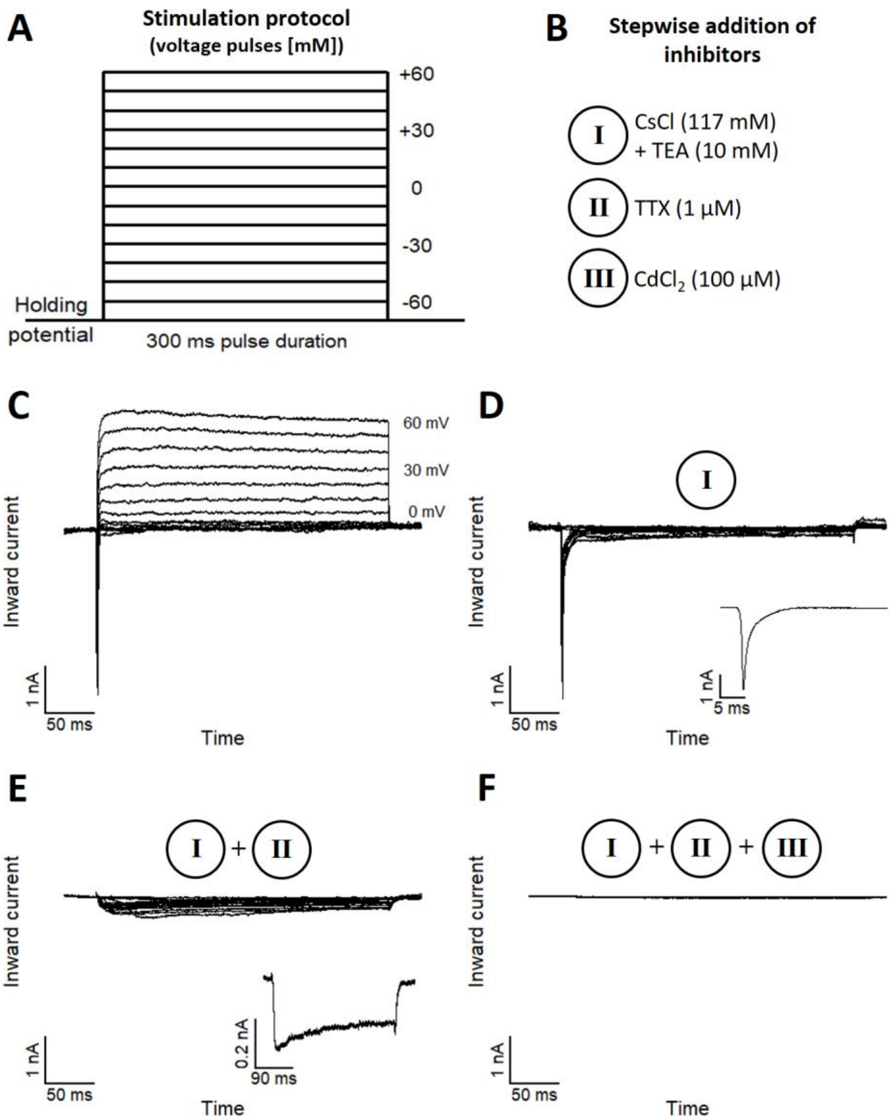

A
Stimulation protocol
(voltage pulses [mM])
+60
+30
0
-30
-60
Holding potential
300 ms pulse duration
B
Stepwise addition of inhibitors
I CsCl (117 mM) + TEA (10 mM)
II TTX (1 µM)
III CdCl2 (100 µM)
C
60 mV
30 mV
0 mV
Inward current
1 nA
50 ms
Time
D
I
Inward current
1 nA
5 ms
1 nA
50 ms
Time
E
I + II
Inward current
0.2 nA
90 ms
1 nA
50 ms
Time
F
I + II + III
Inward current
1 nA
50 ms
Time

**Supplementary figure 7: Expression of functional voltage-gated ion channels**

PNN used on DoD25-35 for voltage-clamp experiments to investigate the expression of voltage-gated ion channels by a step-wise blocking approach. **(A)** Schematic representation of the stimulation protocol used for voltage-clamp recordings. Voltage pulses were applied with a pulse duration of 300 ms and a pulse frequency of 0.2 Hz, starting at -70 mV and increasing in steps of +10 mV. **(B)** Overview of the step-wise blocking approach to differentially assess major cation channel classes. Inhibitors used for the step-wise blocking approach: (I) $K_V$ channel currents were blocked by tetraethylammonium (TEA) combined with the intracellular replacement of $K^+$ ions by $Cs^+$. (II) Block of $Na_V$ channel currents was achieved by tetrodotoxin (TTX). (III) $Ca_V$ channel currents were blocked by $CdCl_2$. **(C)** Representative current responses of a cell in normal recording buffer without any inhibitors added. **(D)** Recording of $Na_V$ currents (with small contribution of $Ca_V$ channels) after the block of $K_V$ channel currents. Inset shows a single trace of a remaining $Na_V$ current at a -30 mV pulse. **(E)** Conditions as in **(D)**, but additional block of $Na_V$ channel currents. Inset shows a single trace of a $Ca_V$ channel current (at V=0 mV) with characteristic, slow inactivation kinetics. **(F)** Condition as in **(E)**, but additional block of $Ca_V$ channel currents, resulting in a complete block of all current responses at all voltage pulses.

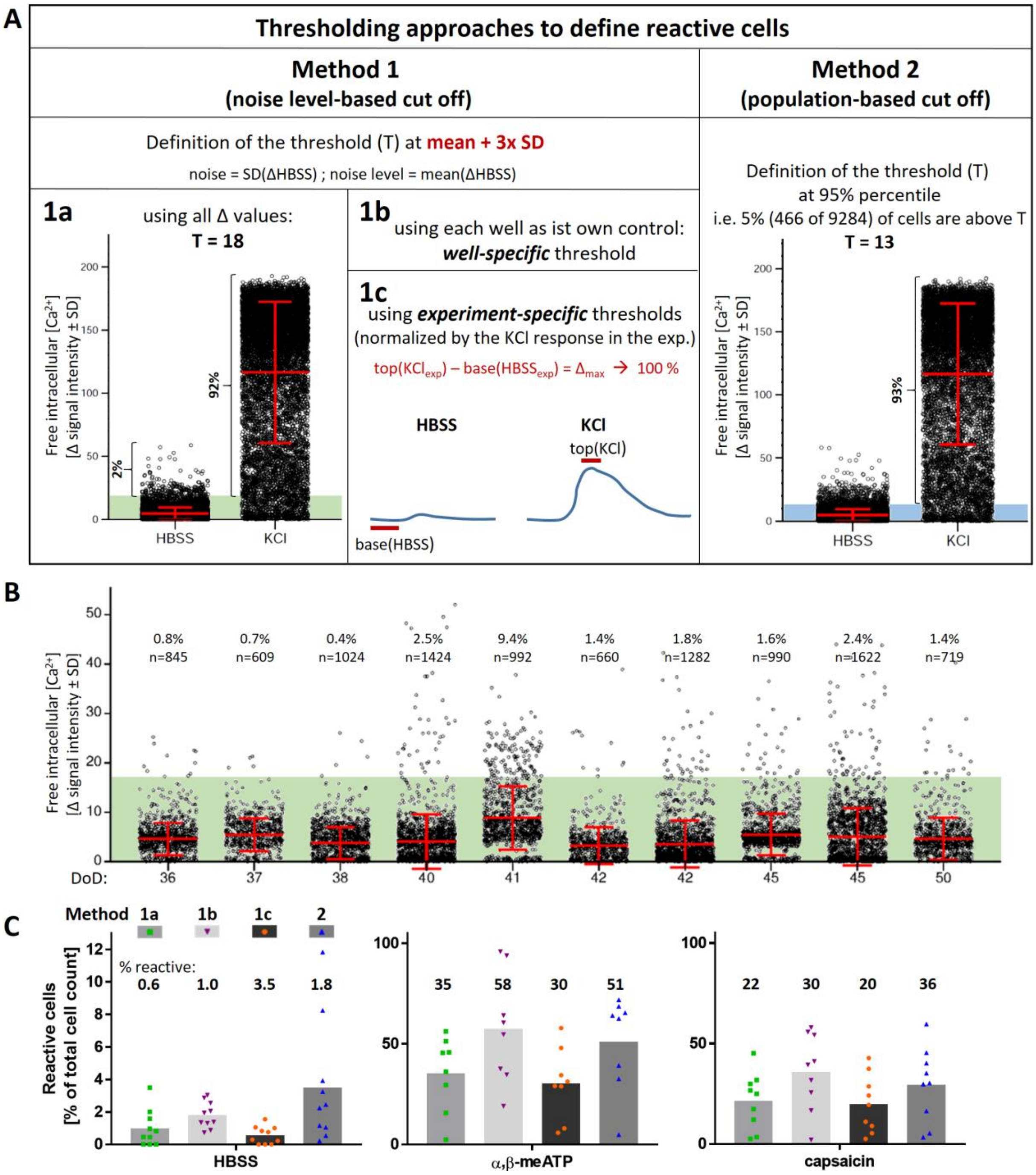

A
Thresholding approaches to define reactive cells
Method 1
(noise level-based cut off)
Method 2
(population-based cut off)
Definition of the threshold (T) at mean + 3x SD
noise = SD(ΔHBSS) ; noise level = mean(ΔHBSS)
1a
using all Δ values:
T = 18
1b
using each well as ist own control:
well-specific threshold
1c
using experiment-specific thresholds
(normalized by the KCl response in the exp.)
top(KClexp) – base(HBSSexp) = Δmax → 100 %
HBSS
KCl
top(KCl)
base(HBSS)
Definition of the threshold (T)
at 95% percentile
i.e. 5% (466 of 9284) of cells are above T
T = 13
Free intracellular [Ca2+]
[Δ signal intensity ± SD]
2%
92%
93%
HBSS
KCl
B
0.8% n=845
0.7% n=609
0.4% n=1024
2.5% n=1424
9.4% n=992
1.4% n=660
1.8% n=1282
1.6% n=990
2.4% n=1622
1.4% n=719
DoD:
36 37 38 40 41 42 42 45 45 50
C
Method 1a 1b 1c 2
% reactive:
0.6 1.0 3.5 1.8
Reactive cells
[% of total cell count]
HBSS
35 58 30 51
α,β-meATP
22 30 20 36
capsaicin

**Supplementary figure 8: Thresholding approaches for $Ca^{2+}$ imaging evaluation to define reactive cells**

**(A)** Summary and illustration of the approaches as described in the supplementary methods that were investigated for threshold setting. Thresholding was always based on the response to the negative control (Hanks' Balanced Salt Solution (HBSS)). The green area in **1a** covers all cells with ΔHBSS<18, while the blue area in **2** covers all cells with ΔHBSS<13. (B) The application of method 1a (T=18) on the HBSS Δ values of the single biological replicates of the test set is exemplified. The percentage of reactive cells is given. The number of all cells measured in each experiment is also indicated. Each dot represents a single cell measured. The means ± SD are indicated graphically in red for each experiment. The green area covers all cells defined as "non-reactive". DoD: day of differentiation (C) Evaluation of reactive cells according to all 4 threshold setting methods using all technical replicates available for each of the 10 biological replicates (also including wells that were not part of the test set): Quantification of reactive cells in response to HBSS, α,β-methylene ATP (1 µM) and capsaicin (1 µM) is shown as mean (grey bars) in % of the total cell number. Each data point represents one replicate (each with several hundred cells). The well-specific method **1b** was chosen as standard thresholding approach for all experiments with an upper threshold limit of T=18 (method **1a**).

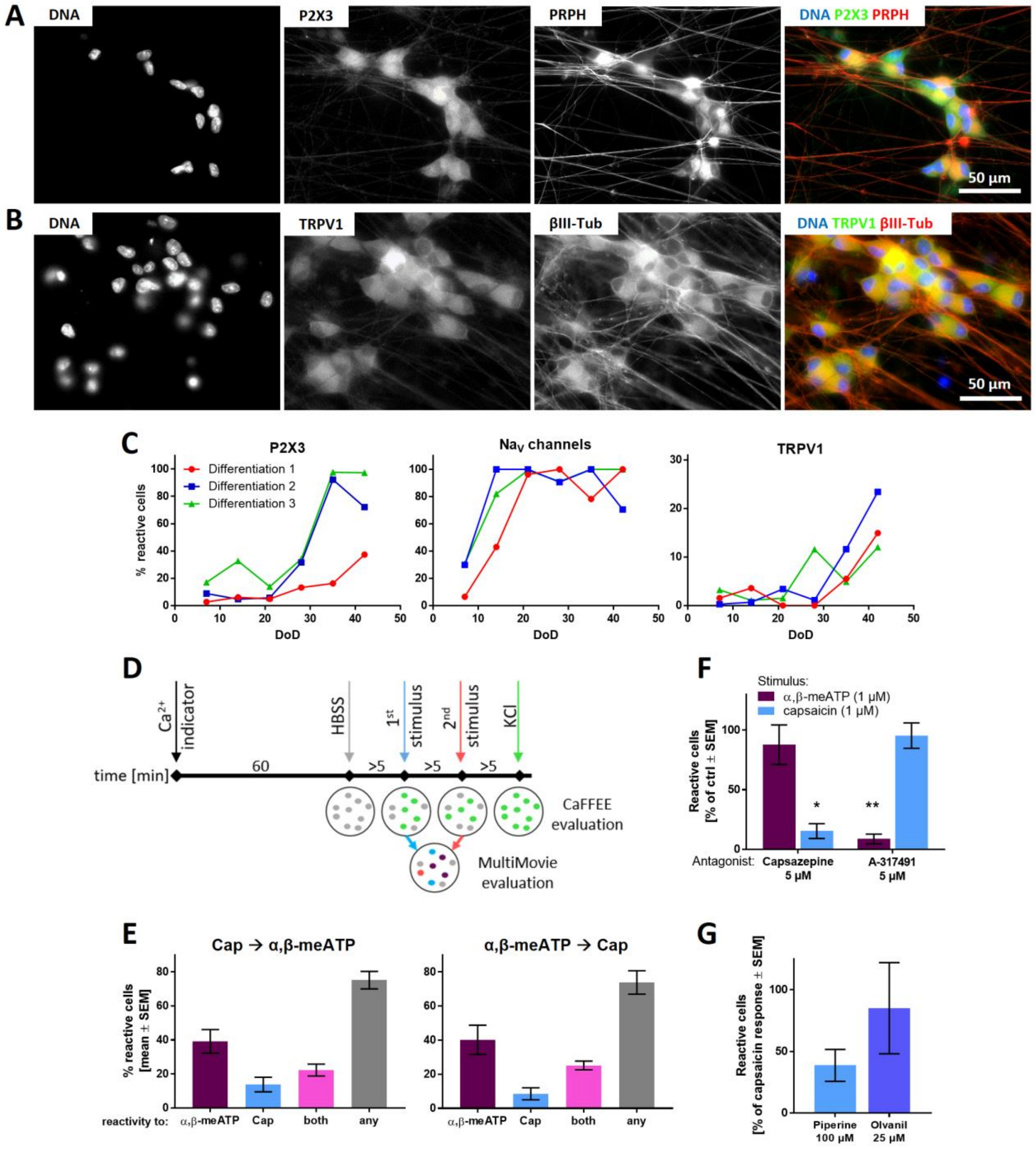
A
DNA
P2X3
PRPH
DNA P2X3 PRPH
50 µm
B
DNA
TRPV1
βIII-Tub
DNA TRPV1 βIII-Tub
50 µm
C
P2X3
Differentiation 1
Differentiation 2
Differentiation 3
% reactive cells
DoD
Na$_V$ channels
TRPV1
D
Ca$^{2+}$ indicator
HBSS
1$^{st}$ stimulus
2$^{nd}$ stimulus
KCl
time [min]
60
>5
CaFFEE evaluation
MultiMovie evaluation
F
Stimulus:
α,β-meATP (1 µM)
capsaicin (1 µM)
Reactive cells [% of ctrl ± SEM]
Antagonist:
Capsazepine 5 µM
A-317491 5 µM
E
Cap → α,β-meATP
α,β-meATP → Cap
% reactive cells [mean ± SEM]
reactivity to:
α,β-meATP
Cap
both
any
G
Reactive cells [% of capsaicin response ± SEM]
Piperine 100 µM
Olvanil 25 µM

**Supplementary figure 9: Characterization of PNN and their use in $Ca^{2+}$ imaging experiments**

**(A,B)** Immunofluorescence images of cells stained for P2X3 and peripherin (PRPH) **(A)**, or TRPV1 and βIII-tubulin (βIII-Tub) **(B)**. Single staining images of the composite images shown in Fig. 6A are given and the composite image is color-coded. **(C)** Weekly $Ca^{2+}$-measurements monitored the development of three independently differentiated cultures from DoD7-42. Responses towards the specific agonists of P2X3 (α,β-meATP, 1 µM) and TRPV1 (capsaicin, 1 µM), and towards a $Na_V$ channel modulator (veratridine, 3 µM) were quantified. **(D)** Sequence of a typical $Ca^{2+}$ imaging experiment conducted in one well. Four different stimuli were applied to the same well with >5 min intervals inbetween. Each measurement can be evaluated for itself using the CaFFEE program. Combination of several measurements for evaluation in the MultiMovie program allows determination of possible polymodality. **(E)** Characterization of cells regarding P2X3- and TRPV1-polymodality using the MutliMovie program. Additionally, the influence of the sequence of applied stimuli was investigated. **(F)** The specificity of the TRPV1-antagonist capsazepine and the P2X3-antagonist A-317491 was investigated with α,β-meATP as the first, and capsaicin as the second stimulus. Data are normalized to the respective stimulus response of cells not pre-incubated with an inhibitor and given as means ± SEM. Significance was tested for treatments ± antagonist. * $P < 0.05$, ** $P < 0.005$ **(G)** Response of sensory neuronal cultures towards the TRPV1 agonists piperine [and olvanil. Data are normalized by the culture response to capsaicin (=100%) and given as means ± SEM.

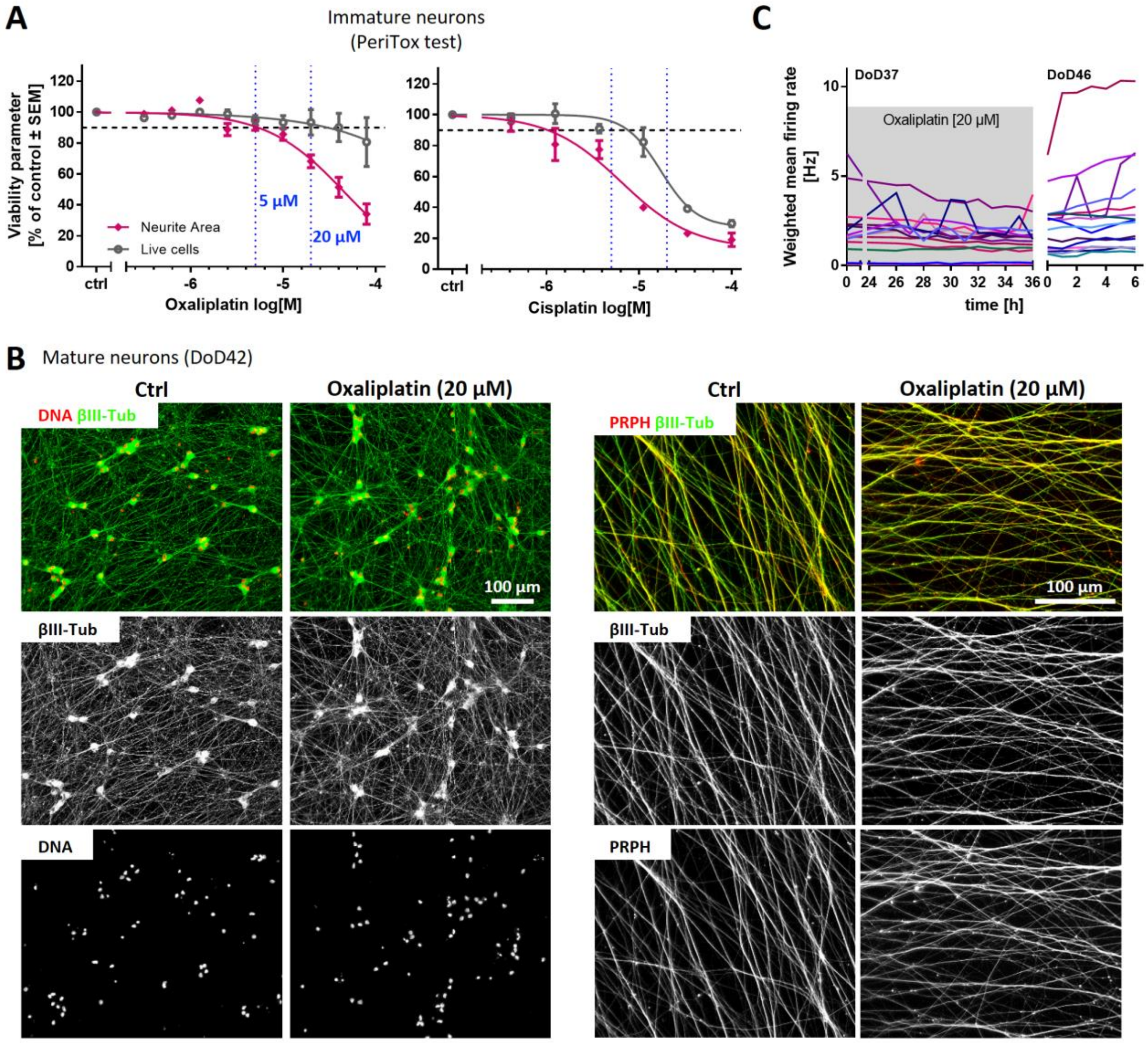

A
Immature neurons
(PeriTox test)
Viability parameter [% of control ± SEM]
Neurite Area
Live cells
5 µM
20 µM
Oxaliplatin log[M]
Cisplatin log[M]
ctrl
B
Mature neurons (DoD42)
Ctrl
Oxaliplatin (20 µM)
DNA βIII-Tub
PRPH βIII-Tub
βIII-Tub
DNA
PRPH
100 µm
C
DoD37
DoD46
Oxaliplatin [20 µM]
Weighted mean firing rate [Hz]
time [h]

**Supplementary figure 10: Effects of platinum compounds on viability parameters**

**(A)** The PeriTox-test was used to assess concentration-dependent effects on neurite growth (pink) and cell survival (grey) of oxaliplatin (left) and cisplatin (right). Horizontal, dashed line: 'no effect' cut-off (90%) for live cells. Vertical, dotted lines: concentrations (5 µM, 20 µM) used in $Ca^{2+}$ imaging experiments. **(B)** PNN on DoD42 were either exposed to oxaliplatin for 24 h before fixation or not (Ctrl). PNN imaged at low magnification (left) after staining for βIII-tubulin (βIII-Tub) and DNA did not reveal significant impairments of either viability or neurite structure. The representative image data were confirmed by qualitative high content imaging of the neurite area. Images on the right focus on neurites at higher magnification. Microtubules (βIII-Tub) and intermediate filaments (PRPH) were stained to visualize potential damage to neurites. Blinded observers were not able to distinguish Ctrl PNN from oxaliplatin-treated PNN. Composite images are colour coded and the single stains are shown in b/w. **(C)** PNN were cultured on multi-electrode array (MEA) plates. On DoD37, oxaliplatin was added for 36 h. Right after oxaliplatin addition and during the last 12 h of the treatment (hourly), the mean firing rate was recorded. Data are given as the 15 min averaged weighted mean firing rates. The grey box indicates oxaliplatin treatment. Presence of oxaliplatin for 24 h did not alter the mean firing rate. After 36 h, oxaliplartin was washed-out. On DoD46, integrity was assessed visually amd firing signals were recorded for 6 h, demonstrating PNN viability.

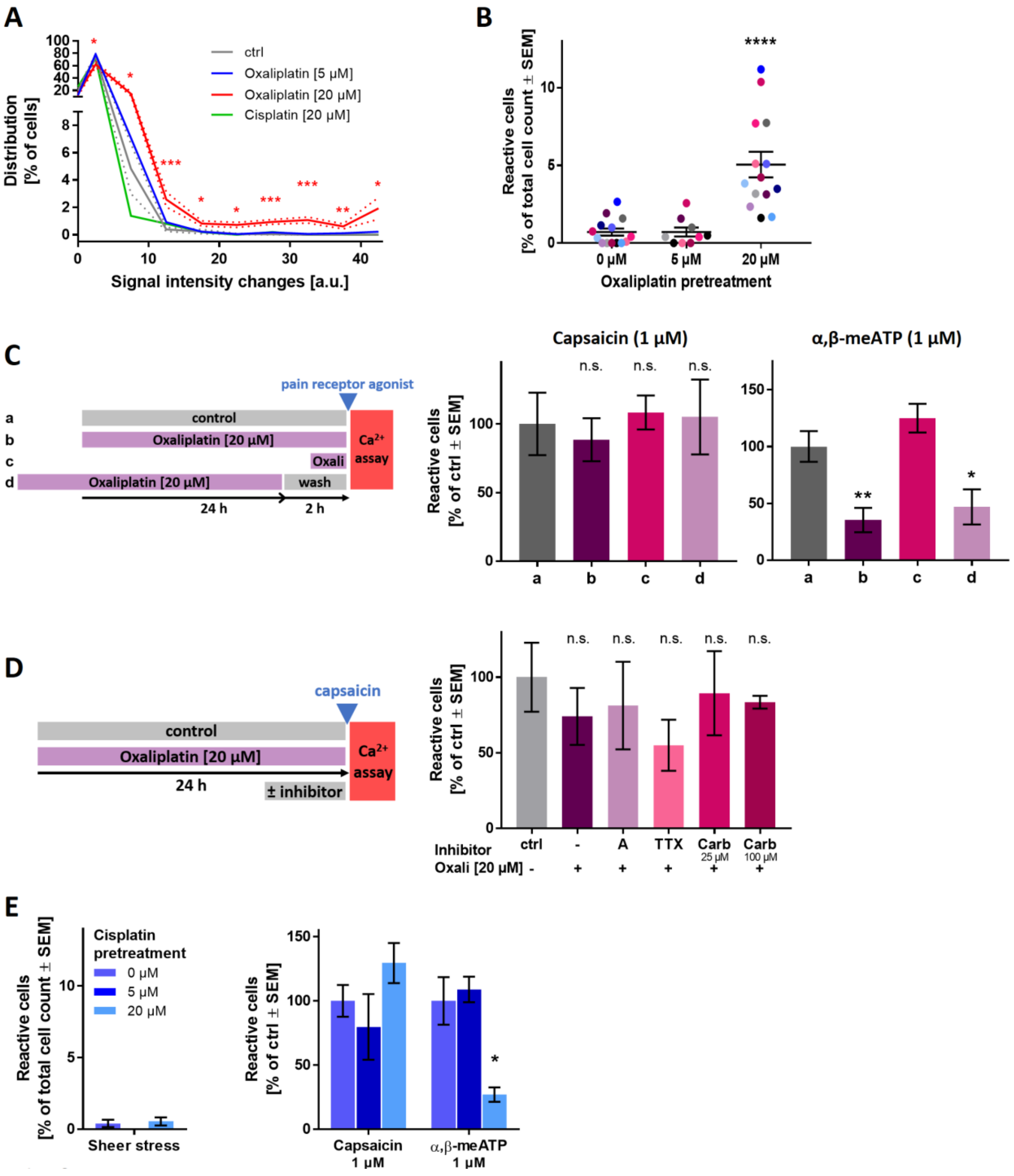
A
ctrl
Oxaliplatin [5 µM]
Oxaliplatin [20 µM]
Cisplatin [20 µM]
Distribution [% of cells]
Signal intensity changes [a.u.]
B
Reactive cells [% of total cell count ± SEM]
0 µM
5 µM
20 µM
Oxaliplatin pretreatment
C
pain receptor agonist
control
Oxaliplatin [20 µM]
Oxali
wash
Ca2+ assay
24 h
2 h
Capsaicin (1 µM)
α,β-meATP (1 µM)
Reactive cells [% of ctrl ± SEM]
n.s.
D
capsaicin
control
Oxaliplatin [20 µM]
24 h
± inhibitor
Ca2+ assay
Reactive cells [% of ctrl ± SEM]
Inhibitor
Oxali [20 µM]
ctrl
A
TTX
Carb 25 µM
Carb 100 µM
E
Cisplatin pretreatment
0 µM
5 µM
20 µM
Reactive cells [% of total cell count ± SEM]
Sheer stress
Reactive cells [% of ctrl ± SEM]
Capsaicin 1 µM
α,β-meATP 1 µM

**Supplementary figure 11: Functional changes in mature PNN exposed to platinum compounds**

**(A)** PNN were pre-treated with oxaliplatin or cisplatin for 24 h and their response to mechanical stress was assessed (see figure 7A). For each signal intensity change value (in arbitrary units, a.u.), the number of cells responding accordingly was determined. This was normalized to the total number of cells. This way, signal intensity change distribution curves (=relative frequencies of $[Ca^{2+}]_i$ changes) were obtained for all conditions. This revealed a significant subpopulation of oxaliplatin pre-treated PNN exhibiting increases in the $[Ca^{2+}]_i$ change. The data from these curves formed the rationale for us to choose the general threshold of T=18 for the quantifications shown in figures 7B and S11B. **(B)** PNN were pre-treated with oxaliplatin for 24 h and their response to mechanical stress was assessed. Each dot represents one biological replicate. Experiments are represented by matching colours. **(C)** The influence of different oxaliplatin exposure schedules (**a-d**) on the responsiveness towards capsaicin (left) or α,β-meATP (right) was investigated. **(D)** Cells were pre-treated with oxaliplatin for 24 h. A-317491 (A, 10 µM), TTX (3 µM) or carbamazepine (Carb) were added to the PNN 1 h before stimulation. Then, the response towards capsaicin (1 µM) application was assessed. **(E)** PNN were pre-treated with cisplatin for 24 h. Responses towards mechanical stress (left), capsaicin and α,β-meATP (right) were assessed. **(A-E)** Data are means ± SEM. Significance was tested against the respective control. * $P < 0.05$, ** $P < 0.005$, *** $P > 0.0005$

### Supplementary References

[1] Chen G, Gulbranson DR, Hou Z et al. (2011) Chemically defined conditions for human iPSC derivation and culture. Nat Methods 8, 424–429.

[2] Dreser N, Madjar K, Holzer A-K et al. (2020) Development of a neural rosette formation assay (RoFA) to identify neurodevelopmental toxicants and to characterize their transcriptome disturbances. Arch Toxicol 94, 151–171.

[3] Klima S, Brüll M, Spreng A-S et al. (2021) A human stem cell-derived test system for agents modifying neuronal N-methyl-D-aspartate-type glutamate receptor Ca2+-signalling. Arch Toxicol 95, 1703–1722.

[4] Fattorelli N, Martinez-Muriana A, Wolfs L et al. (2021) Stem-cell-derived human microglia transplanted into mouse brain to study human disease. Nat Protoc 16, 1013–1033.

[5] García-León JA, García-Díaz B, Eggermont K et al. (2020) Generation of oligodendrocytes and establishment of an all-human myelinating platform from human pluripotent stem cells. Nat Protoc 15, 3716–3744.

[6] Nikasa P, Tricot T, Mahdieh N et al. (2021) Patient-Specific Induced Pluripotent Stem Cell-Derived Hepatocyte-Like Cells as a Model to Study Autosomal Recessive Hypercholesterolemia. Stem Cells Dev 30, 714–724.

[7] Shih P-Y, Kreir M, Kumar D et al. (2021) Development of a fully human assay combining NGN2-inducible neurons co-cultured with iPSC-derived astrocytes amenable for electrophysiological studies. Stem Cell Res 54, 102386.

[8] Terryn J, Welkenhuysen M, Krylychkina O et al. (2018) Topographical Guidance of PSC-Derived Cortical Neurons. J Nanomater 2018, 1–10.

[9] an Verheyen, Diels A, Dijkmans J et al. (2015) Using Human iPSC-Derived Neurons to Model TAU Aggregation. PLoS One 10, e0146127.

[10] Hoelting L, Klima S, Karreman C et al. (2016) Stem Cell-Derived Immature Human Dorsal Root Ganglia Neurons to Identify Peripheral Neurotoxicants. Stem Cells Transl Med 5, 476–487.

[11] Stiegler NV, Krug AK, Matt F et al. (2011) Assessment of chemical-induced impairment of human neurite outgrowth by multiparametric live cell imaging in high-density cultures. Toxicol Sci 121, 73–87.

[12] Schildknecht S, Karreman C, Pöltl D et al. (2013) Generation of genetically-modified human differentiated cells for toxicological tests and the study of neurodegenerative diseases. ALTEX 30, 427–444.

[13] Livak KJ, Schmittgen TD (2001) Analysis of relative gene expression data using real-time quantitative PCR and the 2(-Delta Delta C(T)) Method. Methods (San Diego, Calif.) 25, 402–408.

[14] House JS, Grimm FA, Jima DD et al. (2017) A Pipeline for High-Throughput Concentration Response Modeling of Gene Expression for Toxicogenomics. Front Genet 8, 168.

[15] Love MI, Huber W, Anders S (2014) Moderated estimation of fold change and dispersion for RNA-seq data with DESeq2. Genome Biol 15, 550.

[16] Raudvere U, Kolberg L, Kuzmin I et al. (2019) g:Profiler: a web server for functional enrichment analysis and conversions of gene lists (2019 update). Nucleic Acids Res 47, W191-W198.

[17] Waldmann T, Rempel E, Balmer NV et al. (2014) Design principles of concentration-dependent transcriptome deviations in drug-exposed differentiating stem cells. Chem Res Toxicol 27, 408–420.

[18] Hamill OP, Marty A, Neher E et al. (1981) Improved patch-clamp techniques for high-resolution current recording from cells and cell-free membrane patches. Pflugers Arch 391, 85–100.

[19] R Core Team R: A language and environment for statistical computing. R Foundation for Statistical Computing, Vienna, Austria 2020. https://www.R-project.org.

[20] Wilke CO (2020) Streamlined Plot Theme and Plot Annotations for 'ggplot2' [R package cowplot version 1.1.1]: Comprehensive R Archive Network (CRAN).

[21] Danker T (2019) tdanker/ephys2: Read, analyze and plot HEKA Patchmaster files, 11.08.2019. https://rdrr.io/github/tdanker/ephys2/.

[22] Wickham H (2016) ggplot2. Elegant graphics for data analysis: Springer international publishing.

[23] Edwards SM (2020) Freshing Up your 'ggplot2' Plots [R package lemon version 0.4.5]: Comprehensive R Archive Network (CRAN).

[24] Karreman C, Klima S, Holzer A-K et al. (2020) CaFFEE: A program for evaluating time courses of Ca2+ dependent signal changes of complex cells loaded with fluorescent indicator dyes. ALTEX 37, 332–336.

### Appendix to supplementary information

DB-ALM Protocol:

Differentiation of human iPSC-derived immature dorsal root ganglia-like cells and their use in the PeriTox-test to test neurotoxicants

### DB-ALM Protocol

EURL ECVAM DATABASE SERVICE
ON ALTERNATIVE METHODS TO
ANIMAL EXPERIMENTATION

© European Union 2006-2013

### INDEX

### Part A. Protocol Introduction

Protocol Name: Differentiation of human iPSC-derived immature dorsal root ganglia-like cells and their use in the PeriTox-test to test neurotoxicants

**Abstract:** This protocol provides information about the differentiation of human induced pluripotent stem cells into immature human dorsal root ganglia-like cells. These cells provide the basis for the PeriTox-test, which allows the identification of specific (peripheral) neurotoxicants.

##### *Résumé*

The current protocol describes the differentiation of human induced pluripotent stem cells (hiPSCs) to immature human dorsal root ganglia-like (iDRG) cells performed according to a previously published protocol (Chambers, Qi et al. 2012) with modifications like the introduction of a freeze-and-thaw step.

These iDRG neurons provide the basis for the PeriTox-test. Besides cytotoxicity, the PeriTox-test allows the identification of neurotoxicants, as it includes the functional endpoint of neurite growth inhibition. The test includes single dose treatment of the cells for 24 hours and a direct image-based readout via Cellomics Array Scan VTI HCS high content reader. Treatment of the cells is only possible in concentration ranges where no precipitation of the compound is visually observed.

##### *Experimental Description*

**Biological Endpoint and Endpoint Measurement:**

The PeriTox-test enables the assessment of peripheral neurotoxicity using human iPSC-derived immature human dorsal root ganglia-like cells. Neurite outgrowth as a functional endpoint as well as cytotoxicity are investigated via high content imaging.

**Endpoint Value:**

Concentration-response-curve (including $EC_{10}$, $EC_{25}$, $EC_{50}$, NOEL…)

**Experimental System:**

The human iPSC line Sigma iPSC0028 (EPITHELIAL-1) (derived from adult human epithelial cells by OSKM retrovirus reprogramming, purchased from Sigma-Aldrich (Cat# IPSC0028)) is used to generate immature human dorsal root ganglia (iDRG) neurons.

Sigma iPSC0028-derived iDRG neurons are further used for the PeriTox-test.

#### Discussion

Work requires S1 cell culture laboratories (genetically modified cells).

The current protocol describes the differentiation of human induced pluripotent stem cells (hiPSC) to immature dorsal root ganglia-like (iDRG) cells performed according to a previously published protocol (Chambers, Qi et al. 2012) with modifications like the introduction of a freeze-and-thaw step. The hiPSC line Sigma iPSC0028 is used for the iDRG cell differentiation in this protocol. The Sigma iPSC0028 line is maintained feeder-free on Laminin-521 in E8 medium and during the first two days of differentiation the cells are maintained E8 medium.

In the PeriTox-test human iPSC-derived immature dorsal root ganglia-like cells are used. No specific ethical approval is required.

The duration of the PeriTox-test is 25 h including 24 h of toxicant exposure.

For the performance of the PeriTox-test special equipment like a high content imaging microscope (e.g. Cellomics ArrayScanVTI, Thermo Fisher) equipped with a 10x lens is needed for image acquisition.

To prevent negative edge effects, the edge wells of a 96-well plate are filled with water and only the inner 60 wells are used for testing.

Compounds that reveal autofluorescence can interfere with the detection of calcein-AM or H-33342 in the microscope read-out. In this case, images are blurry and cells cannot be detected by the software. The PeriTox-test can not be used to assess neurite growth and viability endpoints for such compounds.

Training of operators of approximately 4 weeks is needed. During the differentiation, medium change should be performed as fast as possible to keep cells as short as possible at room temperature. From day of differentiation 7' on, medium should be added carefully and slowly so that cells are not washed away from the plate as the cells detach easier the more dense the culture is. Thawing of the cells has to be performed as fast as possible so that cells are no longer exposed to high DMSO concentrations than necessary.

The PeriTox-test is a high throughput assay. Reproducible and robust results are gained with three technical replicates. With this assay neurite specific compound effects can be identified. The comparison to results of a complementary assay using central neurons, e.g. the NeuriTox test (Krug, Balmer et al. 2013), gives insight about the peripheral specificity of an effect.

#### *Status*

**In Development:**

The general method is published in Hoelting et al. 2016. (Hoelting, Klima et al. 2016)

The exact feeder-free protocol for maintenance of iPSC and their differentiation to iDRG neurons as described here, is not published yet, however, the basic procedure highly comparable to the previously published protocol.

**Known Laboratory Use:**

University of Konstanz (used by different operators in this laboratory)

**Participation in Evaluation Study:**

The test system was not tested in different laboratories yet. But intra-laboratory testing revealed a high robustness across different operators and assay runs.

Different classes of chemotherapeutics, which are known to cause peripheral neuropathies, were identified in the PeriTox test whereas testing on human central neurons missed these compounds. This demonstrates the relevance of this test system for safety testing of drugs.

**Participation in Validation Study:**

No

**Regulatory Accepted:**

No

#### *Proprietary and/or Confidentiality Issues*

The distribution of the protocol or any protocol components is not limited.

#### *Health and Safety Issues*

**General precautions:**

No general precautions.

**MSDS Information:**

In addition to the safety measures regarding the compounds in use, there are no safety measures needed for the performance of this method

#### *Abbreviations and Definitions*

FGF: fibroblast growth factor

DAPT: γ-Secretase Inhibitor IX

DMEM: Dulbecco's minimum essential medium

DMSO: dimethylsulfoxide

E8: Essential 8

FBS: fetal bovine serum

FITC: fluorescein isothiocyanate

HSA: human serum albumine

iDRG: immature dorsal root ganglia-like

iPSC: induced pluripotent stem cell

KSR: knockout serum replacement

MEM NEAA: minimum essential medium non-essential amino acids

PBS: phosphate-buffered saline

RT: room temperature

**Last Update:** 04.01.2021

### Part B. Technical Description

**Procedure Details, Latest Version:** 04/01/2021

**Protocol Name:** PeriTox-test to test neurotoxicants on human iPSC-derived immature dorsal root ganglia-like cells

Final modified protocol

Contact person

Marcel Leist, University of Konstanz, Universitätsstraße 10, 78464 Konstanz

+49 7531 885038

Contact person

Anna-Katharina Holzer, University of Konstanz, Universitätsstraße 10, 78464 Konstanz

+49 7531 885331

#### *Materials and Preparations*

##### CELL OR EXPERIMENTAL SYSTEM

The human iPSC line Sigma iPSC0028 (derived from adult human epithelial cells by OSKM retrovirus reprogramming, purchased from Sigma-Aldrich (Cat# IPSC0028)) is used to generate immature human dorsal root ganglia (iDRG) neurons.

Sigma iPSC0028 -derived iDRG neurons are further used for the PeriTox-test.

##### EQUIPMENT

###### Fixed Equipment

Cellomics Array Scan VTI HCS high content reader (Thermo Fisher)

Centrifuge

Freezer (-20°C and -80°C)

Fridge (4°C)

Humidified incubator (37°C, 5% $CO_2$ in air)

Ice machine

Laminar flow hood for sterile atmosphere

Light microscope

liquid nitrogen storage

Mr.Frosty™ freezing container (Thermo Fisher)

Multichannel pipettes

Micropipettes

Neubauer counting chamber

Water bath

**Consumables**

Gloves

Sterile cryovials

Sterile eppendorf tubes (1.5 ml)

Sterile plastic tubes (15 ml, 50 ml)

Sterile single wrapped pipettes

Sterile 6-well plates (Falcon)

Sterile 96-well plates (Falcon)

Sterile 6 cm cell culture dishes (Sarstedt)

Sterile 0.22 mm bottle-top vacuum filter system (Corning)

Sterile 500 ml storage bottle (Corning)

#### MEDIA, REAGENTS, SERA, OTHERS

| **Material** | **Supplier** | **Catalogue Number** |
|---|---|---|
| 0.5M EDTA UltraPure pH 8.0 | Invitrogen | 15575 |
| 2-mercaptoethanol | Gibco | 31350 |
| Accutase | PAA | L11-007 |
| Apotransferin | Sigma | T-2036 |
| Basic fibroblast growth factor (bFGF) | Invitrogen | 13256029 |
| Calcein-AM | Sigma | 17783 |
| CHIR99021 | Axon Medchem | 1386 |

| | | |
|---|---|---|
| Dulbecco's minimum essential medium (DMEM) F12 | Invitrogen | 21331 |
| DMEM/F12, 15 mM HEPES | Gibco | 11330 |
| DMEM / GlutaMax | Gibco | 31966-047 |
| Dimethyl sulfoxide (DMSO) | Sigma | D2650 |
| Fetal bovine serum (FBS) | PAA | A15-101 |
| Glucose | Sigma | G7201 |
| GlutaMax® | Invitrogen | 35050-061 |
| Hoechst H-33342 | Sigma | 14533-100MG |
| Holo-Transferrin | Sigma | T0665 |
| Human Serum Albumin | Sigma | A6608 |
| Insulin | Sigma | I9278 |
| Knock out DMEM | Gibco | 10829 |
| Knockout serum replacement (KSR) | Gibco | 10208 |
| L-Ascorbic Acid | Sigma | A8960 |
| Laminin-521 | BioLamina | LN521-25 |
| Matrigel | Corning | 354234 |
| Minimum essential medium non-essential amino acids (MEM NEAA) | Invitrogen | 11140 |
| Narciclasine | Sigma | N9789 |
| Sodium chloride (NaCl) | Roth | 3957.1 |
| Noggin | R&D | 719-NG |
| Phosphate buffered saline (PBS) (-$Ca^{2+}$, -$Mg^{2+}$) | Gibco | 14190-169 |
| Phosphate buffered saline (PBS) (+$Ca^{2+}$, +$Mg^{2+}$) | Gibco | 14040-091 |
| Progesterone | Sigma | P-7556 |
| Putrescine | Sigma | P-5780 |

| | | |
|---|---|---|
| Recombinant human FGF | PeproTech | 100-18B |
| ROCK inhibitor Y-27632 | Tocris | 1254 |
| TGF-beta inhibitor SB 43154 | Tocris | 1614 |
| Sodium Selenite | Sigma | S-5261 |
| SU5402 | Tocris | 3300 |
| TGF-β1 | R&D | 240-B/CF |
| Trypan blue stain (0.4%) | Gibco | 15250061 |
| γ-Secretase Inhibitor IX (DAPT) | Merck | 565784 |

**PREPARATIONS**

**Media and Endpoint Assay Solutions**

- 250X-media

| **Components** | **Volume required per 100 ml** |
|---|---|
| DMEM/F12, 15 mM Hepes | 99.515 ml |
| L-Ascorbic Acid | 1.6 g |
| Sodium selenite (0.7 mg/ml) stock solution | 0.485 ml |

Sterile filter and prepare aliquots (2 ml). Freeze at -20°C.

- Holo-Transferrin (1000x)
  Dissolve 10 mg in 1 ml PBS and store aliquots at -20°C. The solution is stable for at least 3 months under these conditions
- 0.05% HSA-Solution
  Add 50 µl of a 10% HSA stock solution (in PBS; A6608-100mg) to 10 ml PBS
- Recombinant human FGF Aliquots (1000x)
  Dissolve 1 mg FGF in 10 ml of 0.05% HSA in PBS. Prepare aliquots and store at -80°C for a maximum of 3 months.
- TGF-β1 Aliquots (1000x)
  Prepare a solution of 0.05% HSA, 4 mM HCl in PBS (add 3.5 µl of 25% (8M) HCl to 7 ml 0.05% HSA-solution). Dissolve 10 µg TGF-β1 in 5.75 ml of the prepared solution. Aliquot and store at -80°C for a maximum of 3 months.
- SB 43154 Aliquots

Add 2404 µl ethanol (100%) to 10 mg of SB 43154 to a final concentration of 10 mM. Aliquot SB 43154 solution in 200 µl aliquots and store at -80°C for a maximum of 1 month.

- Noggin Aliquots

  Add 2 ml 0.1% (m/v) BSA in PBS to 500 µg of noggin to a final concentration of 250 µg/ml. Aliquot noggin solution in 25 µl aliquots and store at -80°C for a maximum of 3 months.
- CHIR 99021 Aliquots

  Add 3.58 ml DMSO to 5 mg of CHIR 99021 to a final concentration of 3 mM. Aliquot CHIR 99021 solution in 120 µl aliquots and store at -20°C.
- Rock inhibitor Y-27632 Aliquots

  Add 15.185 ml sterile water to 50 mg of Y-27632 dihydrochloride to a final concentration of 10 mM. Aliquot Y-27632 solution in 500 µl aliquots and store at -20°C for a maximum of 1 month.
- γ-Secretase Inhibitor IX (DAPT) Aliquots

  Thaw InSolution γ-Secretase Inhibitor IX, which is a 25 mM solution of γ-Secretase Inhibitor IX in DMSO. Aliquot γ-Secretase Inhibitor IX in 40 µl aliquots and store at -20°C. This solution is stable for at least 3 months under these conditions.
- SU5402

  Add 336 µl DMSO to 1mg of SU5402 to a final concentration of 10 mM. Aliquot SU5402 solution in 50 µl aliquots and store at -20°C for a maximum of 1 month.
- Putrescine Aliquots

  Add 31.04 ml of sterile water to 5 g of Putrescine dihydrochloride to a final concentration of 1 M. Aliquot putrescine in 300 µl aliquots and store at -80°C. This solution is stable for at least 3 months.
- Sodium Selenite Aliquots

  *E8 stock solution:*

  Dissolve 0.7 mg sodium selenite in 1 ml destilled water. Sterile filter the solution and freeze aliquots at -20°C.

  *N2-S stock solution*:

  Add 5.78 ml sterile water to 100 mg of sodium selenite to a concentration of 100 mM for the master solution. Either freeze the master solution or dilute master solution 1:200 in sterile water to obtain a final stock concentration of 500 µM. Aliquot the selenium stock solution in 500 µl aliquots and store at -80°C.
- Progesterone Aliquots

  Add 12.08 ml sterile water to 100 mg of progesterone-water soluble powder (contains 76 mg progesterone per 1 g powder, Lot # 048K1642) to a progesterone concentration of 2 mM for the master solution. Either freeze the master solution or dilute master solution 1:20 in sterile water to obtain a final stock concentration of 100 µM. Aliquot the progesterone stock solution in 500 µl aliquots and store at -80°C.

- Calcein-AM Aliquots
  Add 251 µl DMSO to 1 mg of Calcein-AM to a final concentration of 4 mM. Aliquot Calcein-AM solution in 10 µl aliquots and store at -20°C. This solution is stable for at least 1 month under these conditions.
- H-33342 Aliquots
  Add 5 ml sterile water to 5 mg of H-33342 to a final concentration of 1 mg/ml. Aliquot H-33342 solution in 500 µl aliquots and store at +4°C. This solution is stable for at least 6 months under these conditions.
- Accutase Aliquots
  Thaw out Accutase bottle, prepare 10 ml aliquots and store at -20°C. The aliquots are stable for at least 1 month under these conditions.
- Matrigel Aliquots
  Thaw out Matrigel by placing Matrigel bottle on ice until Matrigel becomes liquid. Aliquot Matrigel in 330 µl and 1 ml aliquots and store at -20°C. The aliquots are stable for at least 2 months under these conditions
- KnockOut serum replacement Aliquots
  Thaw out KnockOut serum replacement bottle, prepare 50 ml aliquots and store at -20°C. The aliquots are stable for at least 3 months under these conditions

- Preparation of EDTA dissociation solution
  Dissolve 0.45 g NaCl in 250 ml PBS (-$Ca^{2+}$, -$Mg^{2+}$). Add 250 µl of 0.5M EDTA UltraPure pH 8.0. Sterile filter and store at 4°C.
- Preparation of full E8 medium (E8)
  Mix all the components in the table below in a bottle and sterilize it by filtering through a 0.22 µm filter bottle.Label the bottle with content and date keep, it at 4°C for two weeks. Medium should not be re-warmed, it is imperative to only warm up the medium amount to be used immediatly and not keeping it for more than 2 weeks after the preparation day.

| **Components of medium** | **Volume required per 100 ml** |
|---|---|
| DMEM/F12, 15 mM Hepes | 99.1 ml |
| 250X-media | 0.4 ml |
| Insulin | 0.2 ml |
| Holo-Transferrin | 0.1 ml |

| | |
|---|---|
| Recombinant human FGF | 0.1 ml |
| TGF-β1 | 0.1 ml |

- <u>Preparation of N2-S medium (N2-S)</u>
  The stock solutions of the supplements putrescine (1 M), selenium (500 µM) and progesterone (100 µM) are prepared and aliquoted upon first use and stored at -80°C. Thawed aliquots can be stored at 4°C for up to two weeks. Weigh apotransferin and glucose into a plastic bottle and dissolve in DMEM/F12 medium. Add all medium components and sterilize by filtering through a 0.22 µm filter bottle. Wrap the bottle in aluminium foil, label with content and date keep it at 4°C for two weeks. Medium should not be re-warmed, it is imperative to only warm up the medium amount to be used immediatly and not keeping it for more than 2 weeks after the preparation day.

| **Components of medium** | **Volume required per 100 ml** |
|---|---|
| DMEM/F12 | 98.6 ml |
| Apotransferin | 10 mg |
| Glucose | 155 mg |
| Insulin | 400 µl |
| Putrescine (1 M) | 10 µl |
| Selenium (500 µM) | 6 µl |
| Progesterone (100 µM) | 20 µl |
| GlutaMax® | 1.0 ml |

- <u>Preparation of Knockout serum replacement (KSR) medium</u>
  Mix all the components in the table below in a bottle and sterilize it by filtering through a 0.22 µm filter bottle. Wrap the bottle in aluminium foil, label with content and date. Keep it at 4°C. Medium should not be re-warmed, it is imperative to only warm up the medium amount to be used immediately and not keeping it for more than 2 weeks after the preparation day.

| **Components of medium** | **Volume required per 100ml** |
|---|---|
| Knock out DMEM | 83 ml |
| Knock out serum replacement | 15 ml |
| GlutaMax® | 1 ml |

| | |
|---|---|
| MEM NEAA | 1 ml |
| 2-mercaptoethanol | 100 µl |

- <u>Wash medium</u>
  25% KSR medium and 75% N2-S medium._Medium should not be re-warmed, it is imperative to only prepare and warm up the medium amount to be used immediately.
- <u>Freezing medium</u>
  Add 10% DMSO to fetal bovine serum (FBS) and sterilize by filtering through a 0.22 µm filter. Medium can be used for up to 1 week when stored at 4°C.
- <u>Staining solution</u>
  calcein-AM (11 µM), Hoechst-33342 (11 µg/ml) in PBS. Use within 2 days and avoid light exposure.

**Test Compounds**

- Test and control compounds are stored according to the manufacturer's instructions (e.g. 4°C, room temperature, -20°C), compound aliquots at -80°C.

- Stock solutions should be dissolved in sterile water or DMSO, if possible 1000x more concentrated than the working solution. The used DMSO is stored in a lightproof, air-tight bottle at room temperature.

- Final DMSO concentration on the cells is 0.1% (v/v)

- After dissolving the compounds which are delivered in a solid/powder form, all compound solutions are aliquoted into volumes sufficient for one experiment (i.e. one biological replicate). In this way repeated freezing and thawing which can damage the compound's stability and efficiency can be avoided.

- For conducting an experiment, a compound aliquot is thawed and diluted with *day 0* culture medium with supplements in a separate deepwell-plate.

- All compound dilutions in the deepwell-plate contain 0.4% (v/v) DMSO, so that a final concentration of 0.1% (v/v) DMSO is reached on the cells when 25 µl of test or control compound solution is added to the cells (75 µl of day 0 medium with supplements, without DMSO). The highest compound concentration is diluted with medium 1:250 without DMSO as 0.4% (v/v) is already reached with the DMSO the compound is solved in. The serial dilution (normally 1:3) is done with *day 0* culture medium (75% N2-S, 25% KSR supplemented with CHIR99021 (1.5 µM), SU5402 (5 µM) and DAPT (5 µM)) supplemented with 0.4% (v/v) DMSO.

- The compound dilutions (25 µl each) are added to the cells (75 µl) using a multichannel pipette, 6 filter tips at a time

**Positive Control(s)**

- As positive control, Narciclasine is used in a final concentration of 50 nM.
- Therefore, the Narciclasine preparation follows the same indications as for the test compound: stock solution in DMSO (50 µM) and dilution in day 0 culture medium (75% N2-S, 25% KSR supplemented with CHIR99021 (1.5 µM), SU5402 (5 µM) and DAPT (5 µM)) to a concentration of 200 nM in the deepwell-plate. These controls should have the same amount of solvent as the test compounds
- 25 µl of the compound dilution are added to the cells (75 µl).

**Negative Control(s)**

- As negative control, *day 0* culture medium supplemented with 0.1% (v/v) DMSO is used.
- Therefore, *day 0* culture medium supplemented with 0.4% (v/v) DMSO is prepared in the deepwell-plate
- 25 µl of the DMSO dilution are added to the cells (75 µl)

***Method***

**EXPERIMENTAL SYSTEM PROCUREMENT**

The human iPSC line Sigma iPSC0028 (EPITHELIAL-1) is purchased from Sigma-Aldrich (Cat# IPSC0028).

**ROUTINE PROCEDURES**

**Maintenance of Sigma iPSC0028**

Day -3: Thawing of Sigma iPSC0028 cells (6 cm dish)

(1) Prepare one Laminin-521-coated 6 cm dish:

  a. 4 ml of coating solution are needed per 6 cm dish, solution should cover the dish

  b. Dilute Laminin-521 in PBS (+$Mg^{2+}$, +$Ca^{2+}$) 1:40

  c. Add 4 ml coating solution to the dish, distribute equally and incubate at 37°C for 2 h or at 4°C over night

(2) Pre-warm 10 ml DMEM/F12 and 4 ml E8 medium at 37°C

(3) Thaw 1 vial of frozen Sigma iPSC0028 iPS cells in the waterbath (37°C) until a raisin sized frozen part is still left

(4) Transfer the cell suspension immediately into a 15 ml plastic tube with 10 ml of prewarmed DMEM/F12

(5) Rinse the cryovial with 1 ml DMEM/F12 and combine with other cell suspension

(6) Spin 3 min at 500 x g

(7) Discard supernatant and resuspend the cell pellet in 4 ml E8 medium sothat cell clumps (not single cells!) are floating in the medium

(8) Aspirate the coating solution from one 6 cm dish and add the cell clump solution to the dish. Distribute the clups equally by shaking the dish slowly.

Day 0: Splitting of Sigma iPSC0028 (6 cm dish) (repeat every 7 days / when cells get >70% confluent)

(1) Prepare one Laminin-521-coated 6 cm dish:
   - 4 ml of coating solution are needed per 6 cm dish, solution should cover the dish
   - Dilute Laminin-521 in PBS (+$Mg^{2+}$, +$Ca^{2+}$) 1:40
   - Add 4 ml coating solution to the dish, distribute equally and incubate at 37°C for 2 h or at 4°C over night

(2) Pre-warm 7 ml EDTA dissociation solution, 10 ml DMEM/F12 medium and 4 ml E8 medium at 37°C

(3) Aspirate medium from the Sigma iPSC0028 cells

(4) Wash the cells twice with 2 ml EDTA dissociation solution and aspirate immediately.

(5) Add 1 ml EDTA dissociation solution and incubate for 2 min in the incubator (37°C / 5% CO2)
   - Meanwhile aspirate coating solution from prepared 6 cm dishand add 4 ml E8 (pre-warmed)

(6) Aspirate EDTA dissociation solution carefully from the cells. Add 2 ml pre-warmed DMEM/F12 to the cells ) and pipet 4 - 5 x up and down with a sterile 2 ml plastic pipette. The cells should stay in clumps.

(7) Transfer the cells into a 15 ml plastic tube.

(8) Rinse the Sigma iPSC0028 dish with 8 ml DMEM/F12 medium and transfer to the medium into the same 15 ml plastic tube.Distribute 0.25 ml of the Sigma iPSC0028 cell suspension per per prepared 6 cm dish

**Differentiation of Sigma iPSC0028 cells into immature dorsal root ganglia-like cells**

Day -2:

(1) Prepare DMEM/F12 medium, E8 medium and EDTA dissociation solution and prewarm.

(2) Prepare Matrigel coated plate(s)

- **a.** Matrigel has to cover the plate bottom (therefore 1 ml suspension for one well of a 6-well plate is required)
- **b.** Add cold DMEM/F12 to frozen Matrigel pellet and resolve it so that it is diluted 1:40
- **c.** Add solution to the plate(s) and incubate for 1.5 h at RT or 30 min at 37°C
- **d.** After the incubation time, remove Matrigel solution (you can leave DMEM/F12 on the plate(s) as long as you need to prepare your cells)

(3) Wash Sigma iPSC0028 cells twice gently with 2 ml EDTA dissociation solution

(4) After discarding the EDTA add 1 ml of prewarmed EDTA dissociation solution and incubate 4 min at 37°C.

(5) Add 1 ml of DMEM/F12 medium and detach Sigma iPSC0028 cells from the plate by pipetting with a P1000. Cells should not be clumped, single cell suspension should be ensured by pipetting gently up and down

(6) Collect the cell suspension in a 50 ml plastic tube

(7) Spin 3 min at 500 x g

(8) Remove supernatant

(9) Resuspend pellet in 10 ml DMEM/F12

(10) Spin again 3 min at 500 x g

(11) Remove supernatant

(12) Resuspend cells in 1 ml E8 containing 10 µM ROCK inhibitor

(13) count cells in a Neubauer chamber using Trypan blue

(14) plate 90 000 cells/$cm^2$ on Matrigel coated plate(s) in E8 medium containing 10 µM ROCK inhibitor (for 6 well plate use 1.5 ml medium per well)

Day -1:

(15) Change medium to fresh prewarmed E8 medium containing 10 µM ROCK inhibitor

Day 0':

(16) Cells should have 90% confluency today

(17) Change Medium to prewarmed KSR supplemented with Noggin (17.5 ng/ml) and the special TGF-beta inhibitor SB 431542 (10 µM)

Day 1'-8':

| Day | Medium | | Supplements | | | | |
|---|---|---|---|---|---|---|---|
| | KSR | N2-S | Noggin (17.5 ng/ml) | SB431542 (10 µM) | CHIR99021 (1.5 µM) | SU5402 (5 µM) | DAPT (5 µM) |
| 1' | x | | x | x | | | |
| 2' | x | | x | x | x | x | x |
| 3' | x | | x | x | x | x | x |
| 4' | x (75%) | x (25 %) | x | x | x | x | x |
| 5' | x (50%) | x (50%) | | | x | x | x |
| 6' | x (50%) | x (50%) | | | x | x | x |
| 7' | x (25%) | x (75%) | | | x | x | x |
| 8' | x (25%) | x (75%) | | | x | x | x |

Day 9': (Freezing of cells)

(18) Prepare freezing medium and wash medium

(19) Discard the cell supernatant, add 500 µl Accutase per each of the 6 wells and incubate 25 min at 37°C

(20) Add 1 ml wash medium to each well of the 6 well-plate and detach cells from the plate by pipetting with a P1000 (ensure a single cell suspension by pipetting gently up and down). Transfer the cell suspension to a 50 ml plastic tube.

(21) Wash the whole 6 well plate with 1 ml wash medium in total and pool remaining cells in the 50 ml tube

(22) Spin the tube 3 min at 500 x g

(23) Discard supernatant and resuspend the cell pellet in 10 ml of wash medium

(24) Take 10 µl of the cell suspension for counting the cells in a Neubauer chamber using Trypan blue

(25) While counting the cells, spin the remaining cell suspension 3 min at 500 x g

(26) Discard supernatant and incubate the cell pellet on ice for 3 min

*From now on all steps are performed on ice*

(27) Resuspend the cell pellet in freeze medium in a concentration of 8 x $10^6$ cells per ml

(28) Distribute 1 ml of cell suspension per cryovial and put them in a Mr. Frosty freezing container and store them overnight at -80°C

(29) After 24 h store the vials in a cardboard storage box and transfer to liquid nitrogen

#### TEST MATERIAL EXPOSURE PROCEDURES

##### Thawing of dorsal root ganglia-like cells

Day 0:

(1) Prepare wash medium and prewarm
(2) Prepare Matrigel coated plate(s)
   - Matrigel has to cover the plate bottom (therefore 50 µl suspension for one well of a 96-well plate for PeriTox is required)
   - Add cold DMEM/F12 to frozen Matrigel pellet and resolve it so that it is diluted 1:40
   - Add solution to plate(s) and incubate for 1.5 h at RT or 30 min at 37°C
   - After incubation time, remove Matrigel solution (you can leave DMEM/F12 on the plate(s) as long as you need to prepare your cells)
(3) For two 96 well plates thaw one vial of Day 9' cells in the waterbath until a raisin sized frozen part is still left
(4) Transfer the cell suspension immediately into a 15 ml plastic tube with 10 ml of prewarmed wash medium
(5) Rinse the cryovial with 1 ml wash medium and combine with other cell suspension
(6) Spin 3 min at 500 x g
(7) Discard supernatant and resuspend the cell pellet in 1 ml wash medium supplemented with CHIR99021 (1.5 µM), SU5402 (5 µM) and DAPT (5 µM)
(8) count cells in a Neubauer chamber using Trypan blue
(9) plate 100 000 cells/cm$^2$ on Matrigel coated plate(s) in wash medium containing CHIR99021 (1.5 µM), SU5402 (5 µM) and DAPT (5 µM) with a Multipette at speed "3" (for PeriTox use 75 µl medium/well)

##### PeriTox assay

Day 0:

(1) One hour after seeding add 25 µl of prewarmed wash medium supplemented with CHIR99021 (1.5 µM), SU5402 (5 µM) and DAPT (5 µM) containing the test compounds to each well.

Day 1:

(2) 23 h after adding the test compounds, stain cells by adding 10 µl of staining solution to the cells (100 µl medium per well) and incubate 1 h at 37°C.
(3) Read the fluorescence signal using an ArrayScan VTI HCS microscope (Cellomics) to measure cell viability and neurite growth.

**Typical plate layout:**

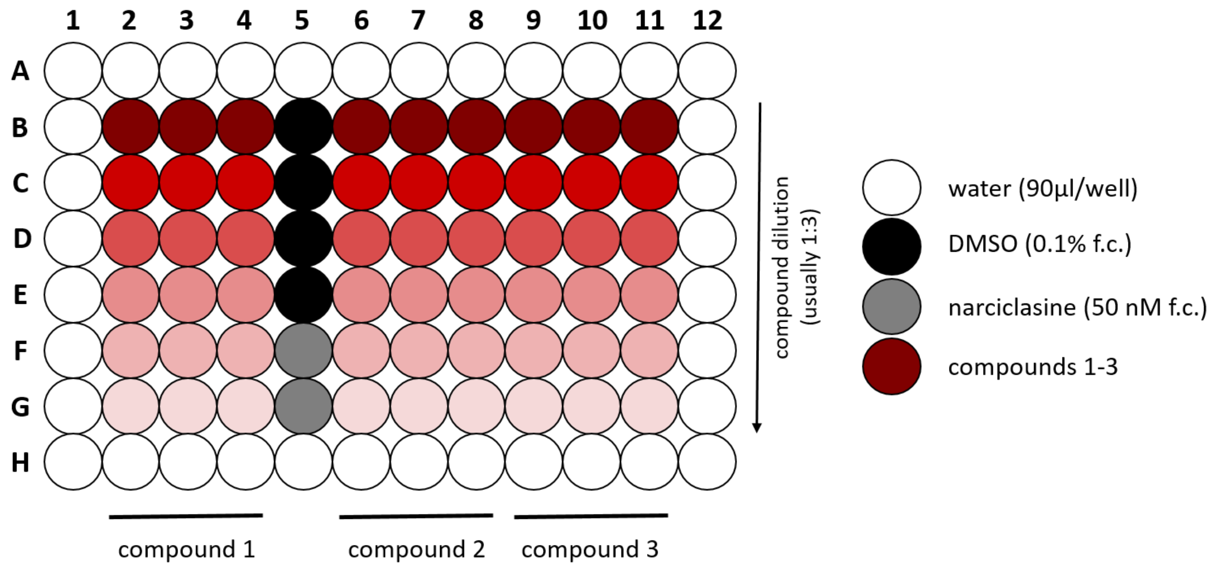

**Figure 1.** Each data point (= one biological replicate) includes usually 3 technical replicates measured on the same plate (see plate layout). For data analysis, at least 3 biological replicates are required.

#### ENDPOINT MEASUREMENT

Cells are stained with calcein-AM to mark viable cells. Co-staining with Hoechst H-33342 allows the identification of any cell.

Cells are stained for 60 min at 37°C and 5% CO2 in the incubator.

The cell staining is imaged in a Cellomics Array Scan VTI HCS reader, using the proper channels for each staining filter. Exposure times are set manually.

To measure the neurite area, the software acquires the Hoechst to identify the cells as objects (via identification of the nuclei), and the calcein-AM in a different channel to measure neurite area. Double positive cells are counted as viable.

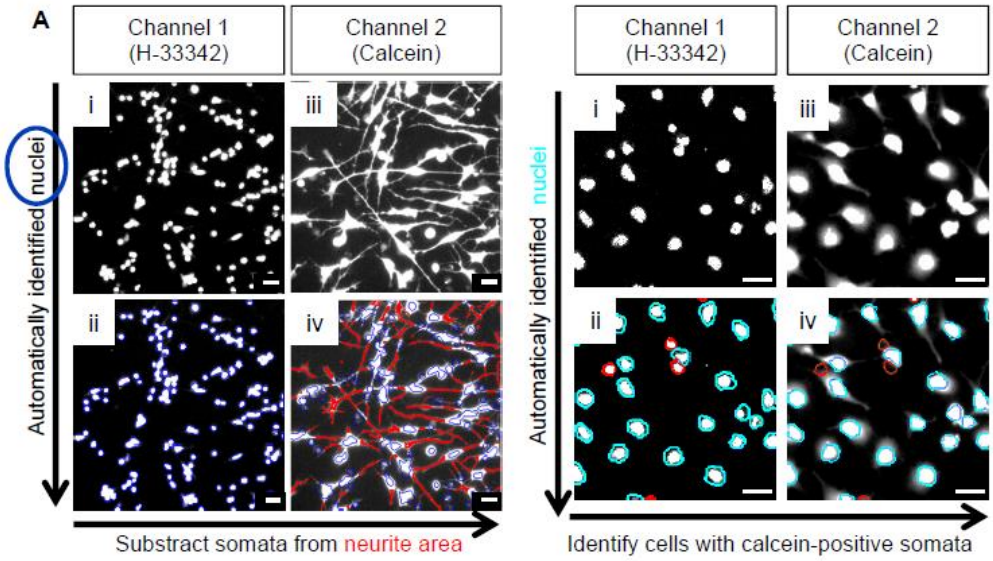

**Figure 2.** Basic principle of the imaging algorithm used for neurite area and viabilityquantification. (**Left side neurite area**): Untreated hESC-derived iDRG are stained withH-33342 and calcein. The nuclei are detected by their H-33342 staining in channel 1 (i);they are automatically identified and marked (indicated by a blue circle) (ii). All viablecells are stained by calcein, which is detected in channel 2 (iii). The algorithmautomatically expands the nuclear outline to define a "virtual soma area". All calcein-positivepixels outside the virtual soma area are defined as neurite area (shown in red)and they are automatically quantified (iv). (**Right side - viability**) Cells of the examplepictures were treated for 24 h with 50 nM epothilone A. (panel i) H-33342 staining; (panelii) automatic identification of cell nuclei, displayed with a color-coded outline of their shape(cyan for normal nuclei, red for objects that are not normal nuclei (e.g. apoptotic nuclei orfragments)). (panel iii) Live cell labelling by calcein. (panel iv) The algorithm quantifies thecalcein intensity in the cells' "virtual soma areas" (cyan circled). Cells with calcein staining below athreshold value are classified by the program as not viable (red circles). (from (Hoelting, Klima etal. 2016)

#### ACCEPTANCE CRITERIA

A rough qualitative evaluation considers the following endpoints on day 1:

Control cells are attached to the plate and neurites are visible under the microscope in phase-contrast.

For toxicity testing:

Positive control narciclasine:

Neurite area $\leq$ 75% of DMSO control

Viability $\geq$ 90% of DMSO control (or not significantly changed)

Negative control DMSO:

Neurite area $\geq$ 150.000

##### *Data Analysis*

The data are analyzed and represented with GraphPad Prism.

For the concentration curve, a nonlinear regression fit is calculated. The fitting method is least squares. If a non-linear curve fit is not possible, a linear curve fit is performed. The curve deriving from the fit is a 4-parameter log function. To calculate the EC50 value, this log-function is solved for y=50% of the total scale, not for 50% of the min-max scale (see example below). Treated concentrations are analyzed for deviation from control. Statistics applied are one-way ANOVA (and nonparametric) with Dunnett's post-test.

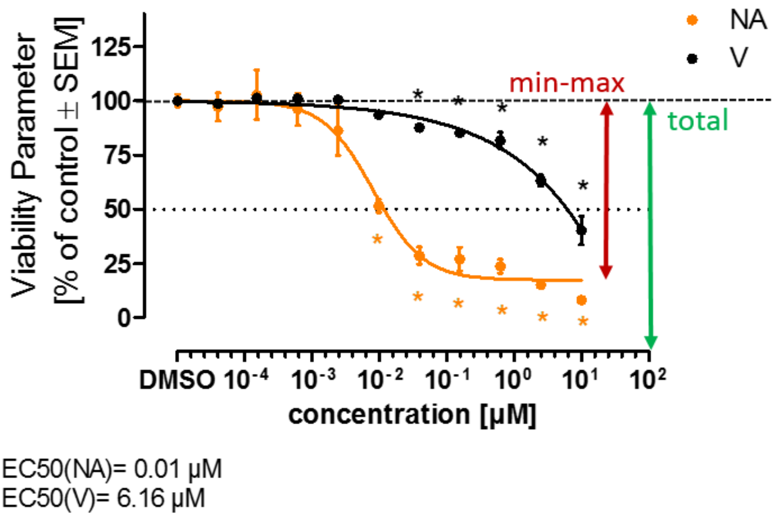

**Figure 3.** Example graph showing the standard representation of neurite area and viability data as % of control using GraphPad Prism. The min-max scale as well as the total scale are depicted in this graph for better understanding of the EC50 calculation. Statistics applied are one-way ANOVA with Dunnett's post-test, statistically significant data points are highlighted by '*'.

***Prediction Model***

**Two different prediction models are used:**

**1. prediction model for screening**:

hit = decrease/increase in neurite area while viability is not changed

(compare to narciclasine positive control:

Neurite area ≤ 75% of DMSO control

Viability ≥ 90% of DMSO control )

**2. prediction model for compound hazard evaluation**:

hit confirmation testing;

EC50 Viability (V) / Neurite Area (NA) ≥ 3 → specifically neurotoxic

A prediction model for the PeriTox test has been developed that allows the classification of toxicants as either specifically neurotoxic, unspecifically cytotoxic or inactive. Specifically neurotoxic compounds decrease the neurite area at concentrations that do not affect the viability of the cells. Unspecifically cytotoxic compounds decrease the neurite area and the viability in the same concentration range to a similar extent. Inactive compounds have no effect on the cells.

To design a prediction model for the PeriTox test, the following steps were taken: (a) use of the "ratio" of EC50 (viability)/EC50 (neurites) as the primary endpoint; (b) measurement of this value for "unspecific toxicants" (the uncoupler CCCP, SDS, Triton-X100, and the topoisomerase inhibitor etoposide; the average ratio was 1.3760.39); (c) definition of a "noise band" (4SD from the average of the ratios of these compounds); and (d) definition of compounds with a ratio outside the noise band (EC50 ratio of.3) as "neurite specific."

This test definition was used to screen three dozen chemicals, preselected for their potential interest for neurite toxicity. 21 positive hits were identified. All tested microtubule drugs, epothilones and proteasome inhibitors were classified as positive hits. All these data were in good agreement with recent clinical findings. (McDonald, Randon et al. 2005, Argyriou, Iconomou et al. 2008, Argyriou, Marmiroli et al. 2011, Miltenburg and Boogerd 2014) Several histone deacetylase (HDAC) inhibitors (HDACi; MS275, SAHA, and TSA) showed neurite toxicity. Among the ROCK pathway modulators, the ρ activator narciclasine was toxic to neurites and inhibitors, such as blebbistatin, accelerated neurite growth. Furthermore, various classes of chemotherapeutic agents and also acrylamide known to cause human peripheral neuropathies were identified using the PeriTox test, although they were missed when tested on human central neurons and vice versa. (Delp, Gutbier et al. 2018)

The US national toxicology program (NTP) assembled a screening library, which consists of different substance classes such as organophosphates, organochlorines, drug-like compounds, pesticides and polycyclic aromatic hydrocarbons (PAHs). This screening library was screened with the PeriTox test and the high level of confirmation (88%) in the hit-confirmation phase indicates that the PeriTox test is technically robust. (Delp, Gutbier et al. 2018)

#### *Annexes*

None.

#### *Bibliography*

Argyriou, A. A., G. Iconomou and H. P. Kalofonos (2008). "Bortezomib-induced peripheral neuropathy in multiple myeloma: a comprehensive review of the literature." Blood **112**(5): 1593-1599.

Argyriou, A. A., P. Marmiroli, G. Cavaletti and H. P. Kalofonos (2011). "Epothilone-Induced Peripheral Neuropathy: A Review of Current Knowledge." Journal of Pain and Symptom Management **42**(6): 931-940.

Chambers, S. M., Y. Qi, Y. Mica, G. Lee, X. J. Zhang, L. Niu, J. Bilsland, L. Cao, E. Stevens, P. Whiting, S. H. Shi and L. Studer (2012). "Combined small-molecule inhibition accelerates developmental timing and converts human pluripotent stem cells into nociceptors." Nat Biotechnol **30**(7): 715-720.

Delp, J., S. Gutbier, S. Klima, L. Hoelting, K. Pinto-Gil, J. H. Hsieh, M. Aichem, K. Klein, F. Schreiber, R. R. Tice, M. Pastor, M. Behl and M. Leist (2018). "A high-throughput approach to identify specific neurotoxicants/ developmental toxicants in human neuronal cell function assays." Altex.

Hoelting, L., S. Klima, C. Karreman, M. Grinberg, J. Meisig, M. Henry, T. Rotshteyn, J. Rahnenfuhrer, N. Bluthgen, A. Sachinidis, T. Waldmann and M. Leist (2016). "Stem Cell-Derived Immature Human Dorsal Root Ganglia Neurons to Identify Peripheral Neurotoxicants." Stem Cells Transl Med **5**(4): 476-487.

Krug, A. K., N. V. Balmer, F. Matt, F. Schönenberger, D. Merhof and M. Leist (2013). "Evaluation of a human neurite growth assay as specific screen for developmental neurotoxicants." Archives of Toxicology **87**(12): 2215-2231.

McDonald, E. S., K. R. Randon, A. Knight and A. J. Windebank (2005). "Cisplatin preferentially binds to DNA in dorsal root ganglion neurons in vitro and in vivo: a potential mechanism for neurotoxicity." Neurobiology of Disease **18**(2): 305-313.

Miltenburg, N. C. and W. Boogerd (2014). "Chemotherapy-induced neuropathy: A comprehensive survey." Cancer Treatment Reviews **40**(7): 872-882.
